# Description and Validation of the Colorectal Cancer and Adenoma Incidence & Mortality (CRC-AIM) Microsimulation Model

**DOI:** 10.1101/2020.03.02.966838

**Authors:** Andrew Piscitello, Leila Saoud, Michael Matney, Bijan J Borah, A Mark Fendrick, Kristen Hassmiller Lich, Harald Rinde, Paul J Limburg

## Abstract

**Background:** Microsimulation models of colorectal cancer (CRC) have helped inform national screening guidelines and health policy decision-making. However, detailed descriptions of particular underlying assumptions are not published, limiting access to robust platforms for exploratory analyses. We describe the development and validation of the Colorectal Cancer and Adenoma Incidence and Mortality (CRC-AIM) microsimulation model, a robust model built to facilitate collaborative simulation studies on disease progression and early detection through screening interventions.

**Design:** We used the Cancer Intervention and Surveillance Modeling Network (CISNET) CRC models, specifically CRC-SPIN, as a foundation for CRC-AIM’s formulas and parameters. In addition, we developed novel submodels and recalibrated various parameters to address gaps and discrepancies in publicly available information. Along with evaluating the natural history and screening detection outcomes from CRC-AIM, we determined the impact of using different life tables (cohort versus period) on natural history outcomes.

**Results:** CRC-AIM demonstrated substantial cross-model validity when comparing multiple natural history and screening outputs and probability curves to those from CISNET models, particularly CRC-SPIN. Additionally, using period life tables, CRC-AIM’s cumulative probability of developing CRC from ages 40 to 100 (7.1%) lies within the range of the CISNET models (6.7% to 7.2%). Using cohort tables, that probability increases to 8.0%. One notable difference is that, regardless of life table used, the cumulative probability of dying from CRC (3.2% for period; 3.8% for cohort) is slightly higher in CRC-AIM than the CISNET models (2.7% to 2.8%), due to CRC-AIM’s different methodology for determining survival. Additionally, there is substantial overlap (e.g. 94-95% overall agreement for strategies on and off the efficient frontier for stool-based strategies) across multiple screening overlay outputs between CRC-AIM and the CISNET models, especially CRC-SPIN.

**Conclusions:** We developed and validated a robust CRC microsimulation model, CRC-AIM, and demonstrate the influence of life table choice on downstream outputs. We further describe CRC-AIM’s parameters and include complete component tables to enhance transparency and encourage collaboration.

## Introduction

The U.S. Preventive Services Task Force (USPSTF), an independent panel of national experts in disease prevention and evidence-based medicine, supplement empirical data with microsimulation models to inform national preventive and screening guidelines. For colorectal cancer (CRC), the USPSTF relies on information such as model outputs from the Cancer Intervention and Surveillance Modeling Network (CISNET) Colorectal Working Group (CWG). The CWG comprises a coordinating center and three independent modeling groups: (1) the Colorectal Cancer Simulated Population model for Incidence and Natural history (CRC-SPIN) from the RAND Corporation; (2) the Simulation Model of Colorectal Cancer (SimCRC) from the University of Minnesota; and (3) Microsimulation Screening Analysis (MISCAN) from Memorial Sloan Kettering and Erasmus University, Rotterdam.^1,2^

These highly sophisticated microsimulation models are powerful tools to simulate the complicated natural history of CRC and precursor lesions in individual patients and identify effective screening interventions for the early detection of CRC.^3–5^ Over time, the model components have evolved and been recalibrated and reparametrized with the emergence of new clinical data (eg, CRC-SPIN v2.x^6^). Although CISNET has documented many changes to the models throughout the years, due to their complexity, it can be difficult to fully determine the downstream effects of recalibration based only on publicly accessible information.^5^

To provide additional transparency and generate an alternative platform for collaborative modeling analyses, we created a robust microsimulation model—the Colorectal Cancer and Adenoma Incidence & Mortality (CRC-AIM) microsimulation model—based on previously reported parameters from CRC-SPIN.^7^ We selected CRC-SPIN as a foundational natural history model because of its comparatively high degree of parsimony, documentation and transparency. Nevertheless, complete details of various components of CRC-SPIN, such as the submodel to determine CRC stage based on size, could not be found in the published literature. This required us to make informed assumptions and create additional submodels to produce a functioning microsimulation model. (See **Supplementary Material** for a detailed description of the differences between CRC-AIM and CRC-SPIN.)

In this publication, we describe the methods used to develop CRC-AIM and demonstrate its robustness by thoroughly comparing its outputs to the published natural history and screening outputs of the three CISNET CWG models (CRC-SPIN, SimCRC, and MISCAN). Furthermore, we examine the consequence of selecting different life tables on natural history outputs. The CISNET models use period life tables, which may underestimate survival because they only describe mortality conditions at a particular time.^1,8^ We explore the consequences of cohort life tables, which build in differential risk across generations by assuming improved mortality rates over time.^8,9^ The CRC-AIM microsimulation model can help inform and address long-standing clinical questions, such as exploring alternate surveillance scenarios and varied colonoscopy performance data.

Ultimately, CRC-AIM not only demonstrates the approach by which existing CRC models can be reproduced from publicly available information, but also provides a ready opportunity for interested researchers to leverage the model for future collaborative projects or further adaptation and testing. To promote transparency and credibility of this new model, we have made available CRC-AIM’s formulas and parameters on a public repository (https://github.com/CRCAIM/CRC-AIM-Public).

## Methods

### Natural History

Natural history modeling for colorectal cancer (CRC) describes the adenoma-carcinoma sequence in the absence of screening for a large population of individuals (**Figure 1**). During one’s lifetime, an individual may generate one or more adenomas. These adenomas independently grow at different rates and may transition into preclinical CRC. The time between adenoma initiation and its transition to preclinical cancer is defined as the *adenoma dwell time*, which is different for each adenoma. In the absence of screening, the time required for a preclinical cancer to become clinically detected, meaning the appearance of disease signs and symptoms, is defined as the *sojourn time* (ST) of the cancer. Each preclinical cancer generated in CRC-AIM is assigned an ST. Upon completion of the ST, the CRC becomes clinically detected and a CRC survival methodology is used to determine the age of cancer cause-specific mortality. If multiple preclinical CRCs exist within an individual, the first cancer to become clinically detected determines survival.^6,10^ The cause of death is considered to be cancer-cause-specific or other-cause-specific, depending on which occurred first in the simulation.

**Figure 1.**
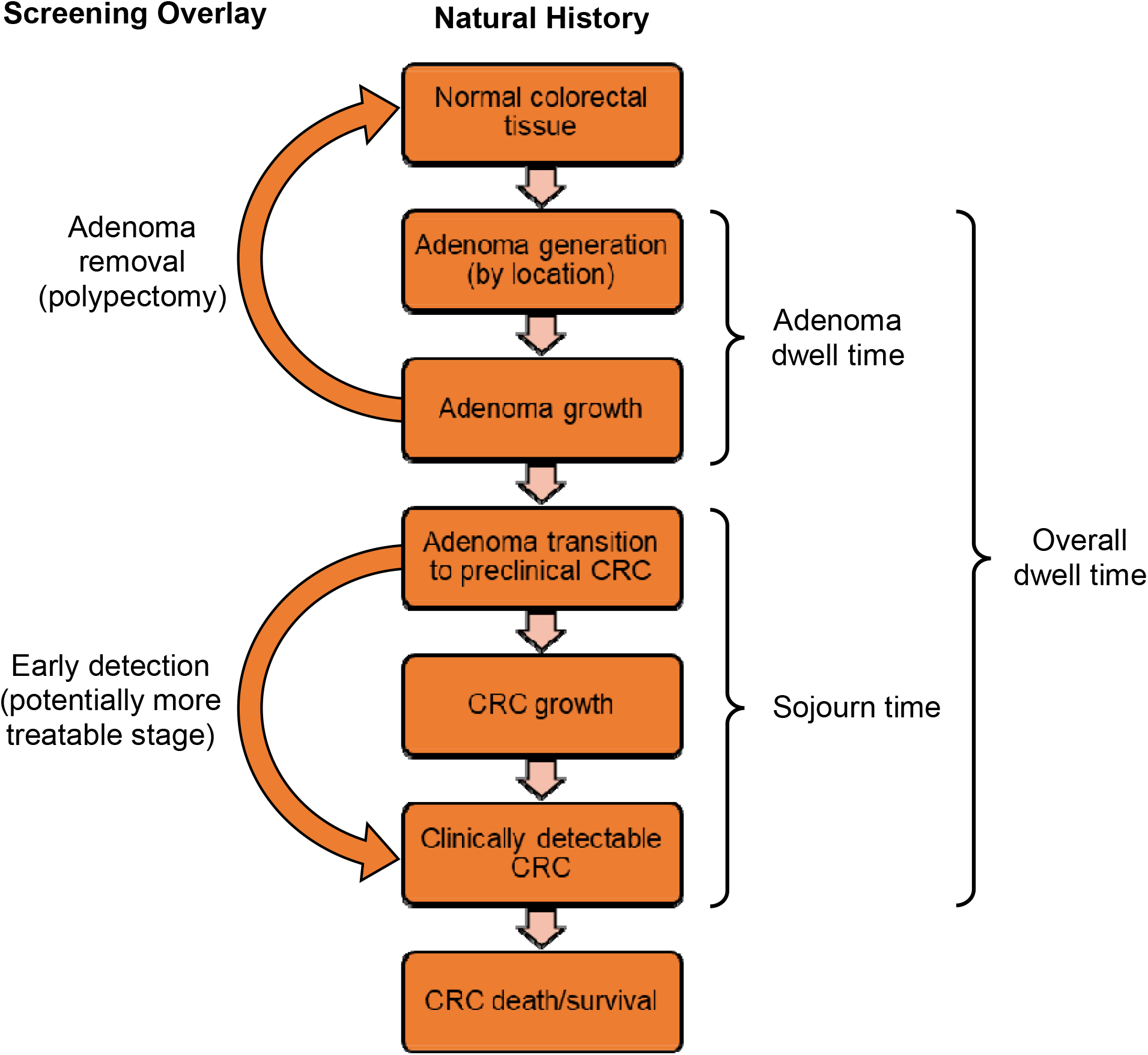
Steps in the natural history of colorectal cancer (CRC) with description of screening consequences.

The model simulates all events until an individual reaches their other-cause mortality date. This approach is analogous to the “parallel universe” model described in CRC-SPIN.^6,10^

See **Supplemental Material** for detailed descriptions of formulas and assumptions related to CRC-AIM’s natural history.

#### Simulating all-cause mortality

To simulate death from other causes, the CISNET models use period life tables, which use mortality rates from a particular period in time to represent mortality rates throughout an individual’s lifetime. CRC-AIM, however, uses cohort (generation) life tables, which determine annual survival from past mortality rates and/or from projected mortality rates. The cohort life tables provide decennial survival information for cohorts born between 1900 and 2010 and project mortality for decennial years 2010 through 2100. We interpolate across birth cohort decade to obtain survival estimates for a specific cohort year and interpolate within a year-of-death interval for specific age at death. The model uses a birth-year-specific sex ratio based on data from the U.S. Census Bureau.^11^

#### Adenoma generation and location

CRC-AIM uses the non-homogenous Poisson process from CRC-SPIN v1.0, specifically the instantaneous risk function, to generate adenomas in an individual.^10^ Adenoma risk is based on an individual’s per-person (inherent) risk, sex, and age. This process assumes no risk of generating adenomas prior to age 20, after which risk generally increases with age. Nonadenomatous polyps are not explicitly modeled because they are not considered to progress to CRC.^12^ Adenomas are localized either to the colon (91% total: 8% cecum, 23% ascending, 24% transverse, 12% descending, 24% sigmoid) or the rectum (9% total), similar to CRC-SPIN v1.0.^10^ Location (colon versus rectum) is used for other natural history components of the model.

#### Adenoma growth

CRC-AIM uses the adenoma growth function from CRC-SPIN v1.0 to describe adenoma size at any time after its generation. The growth function is based on a Janoschek growth curve and assumes that newly generated adenomas are 1 mm in size and can grow to a maximum of 50 mm. Adenomas are not allowed to regress in size, although some grow very slowly, and most of which will never transition to CRC.

#### Transition from adenoma to preclinical CRC

CRC-AIM uses the log-normal function from CRC-SPIN v1.0 that allows adenomas to transition to preclinical cancer.^10^ This function describes the cumulative transition probability of an adenoma transitioning to preclinical cancer at or before adenoma size (*s*) is a function of the adenoma size, age at adenoma initiation (*a*), sex (male vs female), and the location of the adenoma (colon vs rectum), and is defined as:

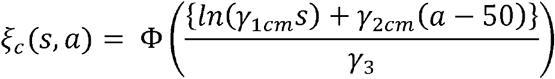

where Φ is a standard normal cumulative distribution function (CDF). There are separate parameters based on adenoma location and sex: colon-male (*cm*), colonfemale (*cf*), rectum-male (*rm*), and rectum-female (*rf*).

CRC-AIM uses a traditional cycle-based approach to aid in interpretability, whereas CRC-SPIN is a continuous time model.^10^ We calculate adenoma size at the start of an interval, the size at the end of an interval (t+1), and determine the probability of transitioning to CRC within the interval conditioned on the probability of having not yet transitioned by the adenoma size at time *t*, defined as:

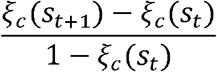

#### Preclinical cancer initiation

When an adenoma transitions to preclinical CRC, multiple events in the model are triggered:

- Similar to CRC-SPIN, CRC-AIM assigns the preclinical CRC an initial size of 0.5 mm and assumes that all CRCs grow exponentially within an adenoma until they overtake the adenoma. As a result, all CRCs are hypothetically detectable, since the minimum adenoma size is 1 mm and colonoscopy can detect 1 mm lesions. (Lesions are defined as the maximum of the adenoma and CRC sizes).^10^
- The sojourn time (ST) is determined.
- Independent of ST, the size of the preclinical cancer upon reaching ST is determined.
- The ST and the initial/final cancer sizes are used to enable the calculation of cancer size at any time.
- Cancer stage at diagnosis is determined based on cancer size upon reaching ST.

#### Transition to clinically detectable CRC

ST is modeled for the *i-*th individual’s *j-*th preclinical cancer using a lognormal distribution that is conditional on location (colon versus rectum), similar to CRC-SPIN.^10^

#### CRC size at clinical detection

Similar to CRC-SPIN, when an adenoma transitions to preclinical CRC, CRC-AIM samples CRC size upon reaching ST.^10^ The distribution of CRC sizes is based on Surveillance, Epidemiology, and End Results Program (SEER) data from 1975-1979 (prior to widespread CRC screening).^10^ However, CRC-SPIN does not describe the form or parameterization of the smoothed distribution for size sampling.

CRC-AIM implements CRC size (*s*) at clinical detection as a generalized log distribution, parameterized by location (μ), scale (σ), and shape (λ), with a maximum CRC size of 140 mm (see **Supplemental Material**). The probability density function is defined as:

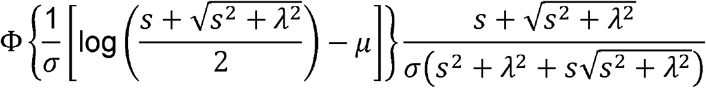

#### CRC growth

During the preclinical CRC phase, CRC-AIM assigns the cancer an initial size of 0.5 mm. We replicated the methodology of CRC-SPIN that describes “flat spots” in the adenoma growth trajectory upon CRC initiation,^10^ which we interpret to mean that upon initiation of a preclinical cancer, the preclinical cancer’s originating adenoma stops growing. We define “lesion size” as the larger value between adenoma size and preclinical cancer size.

Similar to CRC-SPIN, preclinical cancer follows an exponential growth curve in CRC-AIM, with size at time *t* from CRC initiation described as:

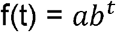

where *a* is the initial CRC size.

If we express initial CRC size as *s_i_*, since the initial (*s_i_*) and final CRC size (*s_f_*) is known, and the time (*t_sf_*) required to reach the final CRC size is known (the independently sampled ST), then the rate (*b*) can be calculated as:

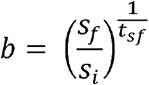

#### CRC stage upon detection

In CRC-AIM’s natural history process, CRC is detected upon reaching ST and the presentation of disease signs and symptoms. (CRC can also be detected in the preclinical cancer growth phase, ie, during ST, during screening)

When CRC is detected, the American Joint Committee on Cancer (AJCC) CRC stage is determined based on its size. Similar to CRC-SPIN v1.0, we derived a multinomial logistic regression model fit to SEER 1975-1979 data to determine AJCC stage based on size (see **Supplemental Materials**). However, the actual form and parameterization of the multinomial logistic regression model used in CRC-SPIN v1.0 is not described. CRC-AIM uses the approach described below.

If we denote CRC size in millimeters as *s* in the *i*-th individual, *k* stage where *k*=1,…,4, then the logit (*g*) for the *i*th individual for the *k*-th category, for categories *k*=1, 2,…*K*-1, is:

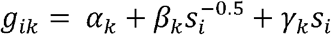

The probability of the *i*-th individual belonging to *k*-th category for categories *k*=1,…,*K* is:

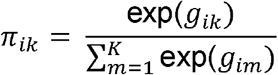

where *g_i,κ_* = 0.

For each individual *i*, the sum of the *k* probabilities adds to 1 and the cumulative probability across increasing stages (ie, *π*_*i*1_, *π*_*i*1_ + *π*_*i*2_, *π*_*i*1_,+ *π*_*i*2_ + *π*_*i*3_) serve as thresholds to define CRC stage based on a uniform (0,1) CDF lookup. If an individual’s CRC is identified prior to clinical diagnosis, its size would be smaller, the thresholds would shift, and the sampled uniform (0,1) particular to that CRC would more likely yield an earlier CRC stage.

#### CRC survival

CRC-AIM implements cause-specific survival as a set of parametric regression equations that model survival probabilities, stratified by location and AJCC CRC stage, as a function of sex and age at diagnosis. To compare the survival outcomes of CRC-AIM to those of the CISNET models, we generated survival curves that mimicked the timeframes of SEER data used by CISNET both before and after their survival update in 2013 (1975-1979 and 2000-2003, respectively). We used code developed by Deborah Schrag to convert pre-1988 SEER registry data from SEER historic staging criteria to AJCC staging categories^13^

We fit five separate parametric linear regression models for each AJCC stage and location (colon versus rectum), based on different distributions to describe survival time: Weibull, lognormal, exponential, Fréchet, and loglogistic (see **Supplemental Materials**). Model effects were sex and age at CRC diagnosis. We based model selection on the smallest Akaike information criterion (AICc) value for the fitted distribution across the five models for each AJCC stage and location (**Table S1**). CRC-AIM uses the CDFs described by the selected model to determine the age at CRC-specific death as a function of age at diagnosis and/or sex. The regression-based coefficients are multiplied by an indicator function if sex or age at diagnosis criteria are met (sex indicator is 1 if met, −1 if not met; age at diagnosis indicator is 1 if met, 0 if not met).

### Screening Overlay

The screening component of CRC-AIM is derived from basic assumptions about CRC screening. In general, CRC screening facilitates the detection and removal of adenomas and preclinical lesions. Each time screening is due, the chance of a lesion to be detected is dependent on the screening test’s sensitivity and reach. The overall effectiveness of screening is dependent on screening frequency, adherence rates, and the sensitivity and specificity of the screening test. False positives can occur in relation to a screening test’s specificity and lead to unnecessary follow-up colonoscopies or unnecessary polypectomies. Complications, including fatal ones, can arise due to polypectomies. Thus, adverse events related to colonoscopies are part of the screening component.

For the purposes of this analysis, the screening test characteristic input assumptions were the same as those used in CISNET models (**Table 2**),^2,14^ which we have reproduced here in greater detail. We assumed that the same test characteristics for screening colonoscopies applied to colonoscopies for diagnostic follow-up or for surveillance, and that there was no correlation in findings between CTC or sigmoidoscopy and subsequent diagnostic colonoscopy. For colonoscopy and sigmoidoscopy, the lack of specificity with endoscopy reflects the detection of nonadenomatous polyps. For sigmoidoscopy, this may lead to an unnecessary diagnostic colonoscopy, and for colonoscopy, this may lead to an unnecessary polypectomy. For computed tomographic colonography, the lack of specificity reflects the detection of ≥6 mm nonadenomatous lesions, artifacts, stool, and adenomas smaller than the 6-mm threshold for referral to colonoscopy that are measured as ≥6 mm. For FIT, we assumed a positivity cutoff of ≥100 ng of hemoglobin (Hb) per mL of buffer (≥20 mcg Hb/g of feces). For FIT and mt-sDNA, the sensitivity for adenomas <10 mm was considered the sensitivity for nonadvanced adenomas. For the high sensitivity guaiac based fecal occult blood test (HSgFOBT), we assumed that 1-5 mm adenomas do not bleed and therefore cannot cause a positive stool test. It was also assumed that HSgFOBT can be positive because of bleeding from other causes, the probability of which is equal to positivity rate in individuals without adenomas.

The sensitivity inputs for stool-based tests are per person and are based on the characteristics of the most advanced lesion. The sensitivity inputs for structural tests are per lesion and potential detection of lesions depend on the reach of the test. It is assumed that sigmoidoscopy completely visualizes the rectum for all individuals, visualizes the sigmoid colon in 88% of individuals, and visualizes the descending colon in 6% of individuals.^3,15,16^ With colonoscopy, full reach (to the cecum) is assumed to be achieved 95% of the time and if reach is only partial, a second colonoscopy is performed.^2^

It is assumed that a follow-up colonoscopy occurs after any positive non-colonoscopy screening test. If the follow-up colonoscopy is negative, individuals return to their original non-colonoscopy screening test and the next screening is due in 10 years. If the follow-up colonoscopy is positive for an adenoma of any size, individuals enter a surveillance colonoscopy period where the next colonoscopy is based on the findings of the latest colonoscopy (3 years if a detected adenoma is ≥10 mm or if ≥3 adenomas of any size are detected; 5 years if 1-2 adenomas <10 mm are detected; these are the same histology assumptions used by CISNET). Surveillance continues until at least age 85 and then halts if no adenomas or CRC are detected on the last surveillance exam or continues past 85 until no adenomas or CRC are detected on a surveillance exam. Individuals with preclinical lesions that become symptomatic based on sojourn time expiration receive a colonoscopy to clinically diagnose interval cancers (CRCs that grow and develop after a screening or surveillance exam but before the next recommended exam).

Three types of complications related to colonoscopies are included in the model: 1) serious gastrointestinal events (e.g., perforations, gastrointestinal bleeding, or transfusions) of which 8.97% were perforations^17^ and 5.19% of perforations led to death^18^; 2) other gastrointestinal events (e.g., paralytic ileus, nausea and vomiting, dehydration, abdominal pain); and 3) cardiovascular events (e.g., myocardial infarction or angina, arrhythmia, congestive heart failure, cardiac or respiratory arrest, syncope, hypotension, or shock). Risk of complications increases by age.^17,18^ (For additional information, see eFigure1 of Knudsen et al.^2^) Because the risk of complications with colonoscopy is conditional on polypectomy,^17,19^ and sigmoidoscopy is modeled without a biopsy or polypectomy of detected lesions, the risk of complications with sigmoidoscopy was therefore assumed to be none.

The screening modalities evaluated for the main validation analysis were FIT, HSgFOBT, mt-sDNA, and colonoscopy.

### CRC-AIM Comparison and Cross-Validation

To evaluate the performance of CRC-AIM, we compared the outputs of CRC-AIM against the natural history experiment outputs of CRC-SPIN, SimCRC, and MISCAN.

We compared the CRC-AIM natural history outputs to those from CRC-SPIN v1.0, as well as the other CISNET models’ outputs, that were generated prior to 2013 and the associated CISNET CRC survival update. For comparisons to these historical (pre-2013) outputs, we selected the 1975-1979 time period for CRC cause-specific survival curves, because this time period had been described as previously used by CRC-SPIN.^10^ We also compared the outputs from CRC-AIM to natural history outputs presented in the 2015 CISNET technical report.^14^ For these comparisons, we used the 2000-2003 time period for CRC cause-specific survival curves, similar to what CISNET used for recently diagnosed CRC.^20^

All CRC-AIM analyses use cohort life tables for non-CRC mortality unless otherwise specified. We attempted to match birth cohorts to those from the original CISNET analyses, but this information was not always reported. Data from reference publications^12,14,21,22^ were extracted using a custom Python package that assigns numeric values based on the pixel location of data within a figure and the corresponding X and Y scale values.

#### Comparison to historic (pre-2013) outputs

Adenoma and CRC Prevalence/Incidence: We compared the adenoma and CRC prevalence/incidence values for individuals at the age of 65, as described by Knudsen et al.^21^ Results were reported either as a rate per thousand individuals or by percentage. Because the cost-basis year used by Knudsen et al was 2007,^21^ and we wanted to replicate their simulation of 65-year-olds, we assumed a birth cohort of 1942 (2007 minus 65). The number of simulated individuals was not reported but we assumed 3 million. Furthermore, we excluded individuals with existing preclinical cancer at age 65 from the CRC cumulative incidence calculations under the assumption that CRC incidence measures clinically diagnosed CRC. In addition, we compared the multiplicity of adenomas (the average number of adenomas in individuals with one or more adenomas), as described in Kuntz et al.^12^

Adenoma Dwell Time, Sojourn Time (ST), and Overall Dwell Time: We replicated the retrospective analysis from Kuntz et al^12^ that calculated the mean, median, and interquartile range (IQR) for adenoma dwell time, the ST, and the overall dwell time (adenoma dwell time plus ST) for individuals with clinically diagnosed CRC. We simulated a cohort of 30 million individuals born in 1944. We also compared CRC-AIM’s outputs to different mean values of these characteristics as reported one year earlier by CRC-SPIN,^22^ although the birth cohort for that analysis was different (CRC-SPIN had simulated 30 million individuals born in 1928).

Estimated Annual Transition Probabilities: We replicated a modeling experiment estimating transition probabilities from Rutter and Savarino.^22^ We simulated 30 million individuals born in 1928 and estimated state-transition probabilities for a cohort of 60-year-olds, which are based on the proportion of individuals making state transitions as they progress from age 60 to 61.

#### Comparison to USPSTF outputs

##### Natural History Comparison

We compared the natural history outputs from CRC-AIM to those described in the 2015 CISNET technical report by Zauber et al.^14^ For each comparison, we replicated the reference population by simulating 2 million individuals using a birth cohort year of 1975.

Prevalence of Preclinical CRC/Adenomas: We replicated the outputs describing the prevalence of preclinical CRC and adenomas for individuals over 40 years old. In our calculation of preclinical cancer prevalence, once an individual developed clinically detectable cancer, they no longer had preclinical cancer and were removed from the numerator. Therefore, this analysis does not represent the cumulative probability of acquiring preclinical cancer by a certain age. In our calculation of adenoma prevalence, we only considered adenomas that were present until clinical CRC developed.

Location/Size of Adenomas: We replicated the location and size of adenomas for individuals over 40 years old. For the size analysis, only the size of the most advanced adenoma (ie, largest lesion) was considered for the adenoma distribution. Again, adenomas were only included until clinical CRC developed.

CRC Incidence by Age: We replicated the incidence of clinically diagnosed CRC cases per 100,000 individuals by age, which was calibrated to 1975-1979 SEER incidence (without screening). This incidence was compared to the empirical SEER incidence rates prior to the diffusion of screening (1975-1979) and after over half of the population had been screened (2007-2011). We calculated the incidence of new clinically diagnosed CRC that occurred between yearly intervals, starting at 2015-2016 (age 40) and projected through 2075-2076 (age 100). Individuals with clinically diagnosed CRC for a given interval were excluded for future intervals.

CRC Stage Distribution at Diagnosis: We generated a stage distribution of clinically diagnosed CRC in individuals aged 40 and older in the absence of screening.

Cumulative Probability of Developing/Dying from CRC: We determined the cumulative probability of developing CRC and dying from CRC, by age, in the absence of screening using a birth year of 1975. This was performed using both cohort life tables, described in detail above, and period life tables, which were obtained from the Centers for Disease Control and Prevention.^8^

Cumulative Probability of Developing CRC in Individuals with/without Underlying Lesions: We replicated an experiment by Kuntz et al^12^ that determined the cumulative probability of developing CRC, stratified by individuals who either did or did not have an existing adenoma or undiagnosed preclinical cancer at age 55. The analysis extended to age 85. The weighted average of the two groups reflects the population-level risk of cumulative CRC incidence across age. Competing cause of mortality (ie, death from other causes) was removed, and the cumulative probability of developing CRC in this analysis was therefore not impacted by choice of life table.

##### Screening Outcomes

We replicated the screening outcomes supplemental tables from the CISNET technical report for each screening modality.^14^ We wanted to reproduce CISNET model outputs as faithfully as possible, and therefore we used the modified CRC-AIM that employs period life tables. Predicted screening outcomes were simulated for a birth year cohort of 1975. The main benefits of CRC screening are life-years gained (LYG) from prevention of CRC cases and delay of CRC deaths compared with no screening. Number of colonoscopies was used to represent burden and harms. The number of tests, complications from colonoscopies, CRC cases, CRC deaths, life-years with CRC, incidence reduction, and mortality reduction were additional screening outcomes. All outcomes were reported per 1,000 individuals free of diagnosed CRC at age 40.

Efficient Frontiers: We replicated the efficient frontier figures from the Figure 3 (colonoscopy) and Figure 4 (stool-based—FIT, HSgFOBT, and mt-sDNA) and corresponding tables from Knudsen et al.^2^ Like CISNET, for both colonoscopy and stool-based efficient frontiers, the screening stop ages were 75, 80, or 85 years. We included only screening start ages of 50 and 55 for colonoscopy and replicated the screening start ages of 50 and 55 for stool-based modalities. Strongly dominated strategies (ie, strategies that colonoscopies for fewer LYG) were discarded. The incremental number of LYG per 1000 (ΔLYG) and incremental number of colonoscopies per 1000 (ΔCOL) were computed. The efficiency ratio (ΔCOL/ΔLYG) for each remaining strategy was calculated. Strategies with fewer LYG but a higher efficiency ratio than another strategy were discarded as weakly dominated. The efficient frontier was the line that connected the efficient strategies; strategies that had LYG within 98% of the efficient frontier were considered near-efficient.

#### Cross-validation of screening outcomes

The purposes of the validation analysis were to cross-validate CRC-AIM’s screening component with other CISNET screening comparative effectiveness results assuming perfect adherence. Colonoscopy and stool CISNET screening modalities were analyzed in the validation analysis. For more information on the comparative analyses, see **Supplemental Materials**.

## Results

We compared the natural history outputs from CRC-AIM with the published outputs from the three CISNET models—CRC-SPIN, SimCRC, and MISCAN—and found considerable similarity among the models.

First, CRC-AIM produced similar natural history outputs compared to CRC-SPIN v1.0, SimCRC, and MISCAN prior to the 2013 CISNET CRC survival update. The adenoma prevalence at age 65 for CRC-AIM is 29.2%, which is similar to the CISNET models (30.7% for CRC-SPIN, 37.2% for SimCRC, and 39.8% for MISCAN) (**Table 1**). Moreover, the multiplicity of adenomas at age 65 for CRC-AIM (1.7) was within the range of values for the CISNET models (1.8 for CRC-SPIN, 1.6 for SimCRC, and 2.0 for MISCAN) (**Table 1**). In addition, the location (proximal colon, distal colon, and rectum) of the number of size-stratified adenomas (1-5 mm, 6-9 mm, and ≥10 mm), along with the cumulative incidence (10-year, 20-year, and lifetime) across CRC stages, was comparable between CRC-AIM and the CISNET models (**Figure 2, Table 1**).

**Figure 2.**
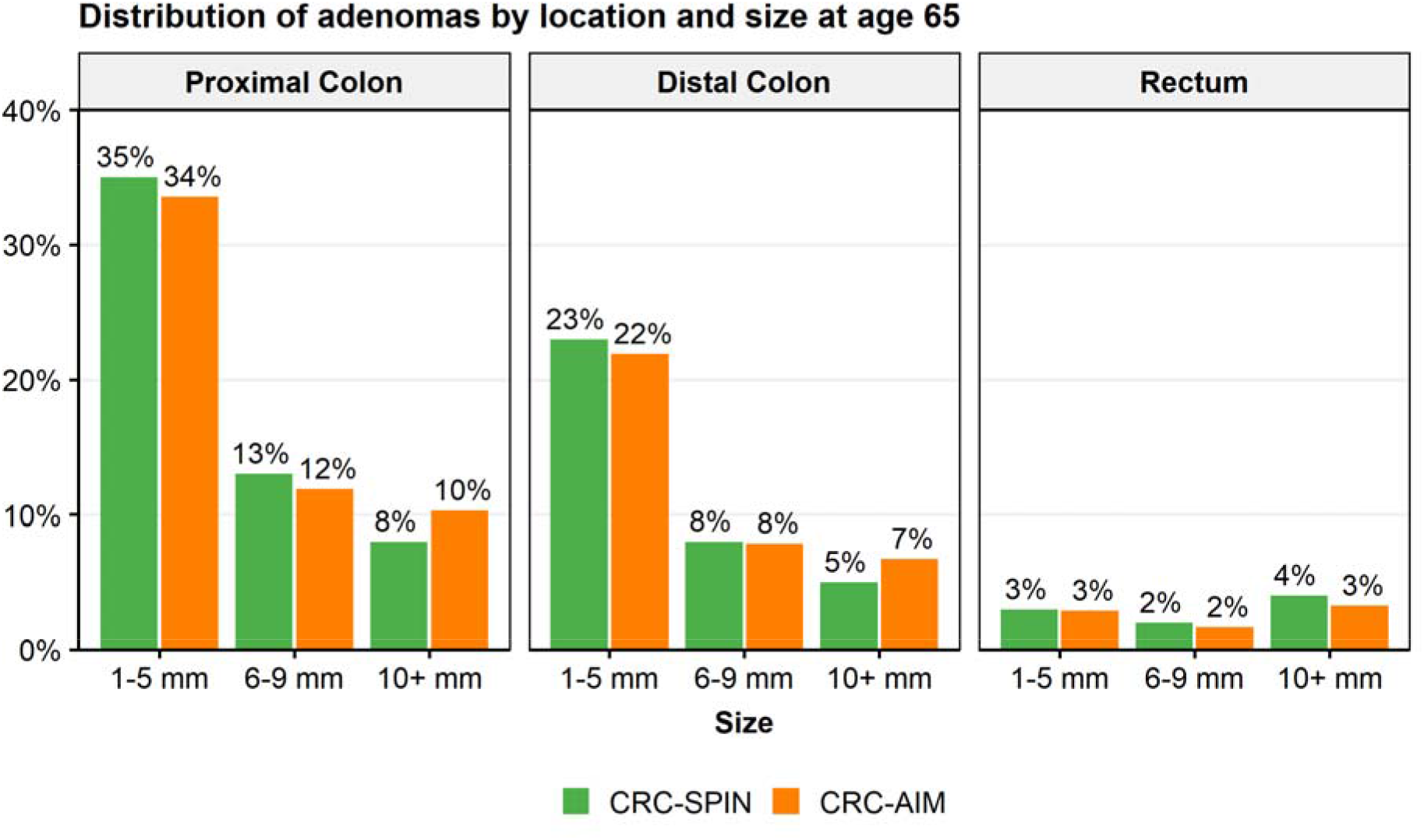
Distribution of adenomas by size and location at age 65 in CRC-SPIN and CRC-AIM. Sizes are subdivided into small (1-5 mm), medium (6-9 mm), and large (10+ mm) adenomas. Data adapted from Knudsen et al.^21^

**Table 1.**
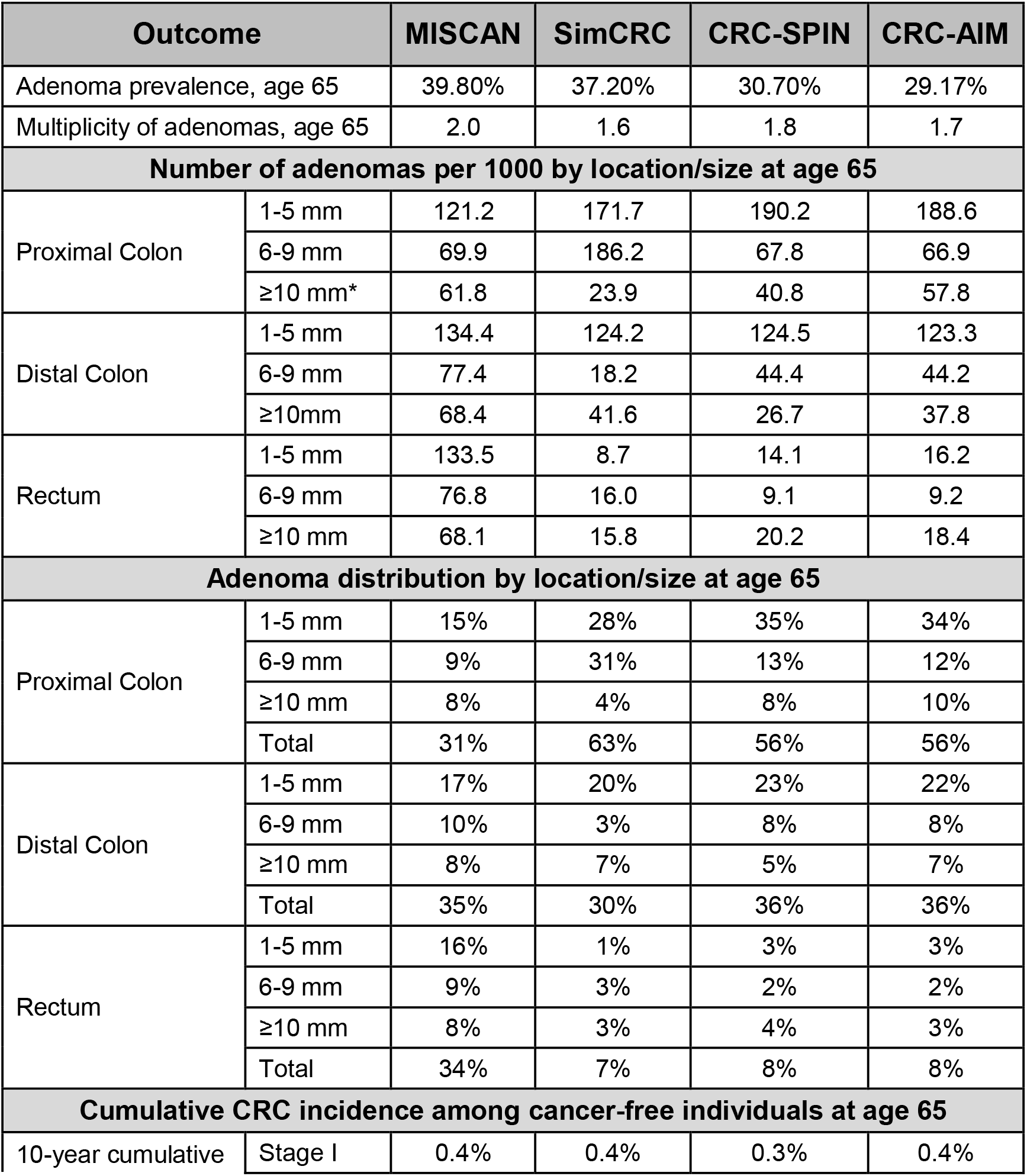
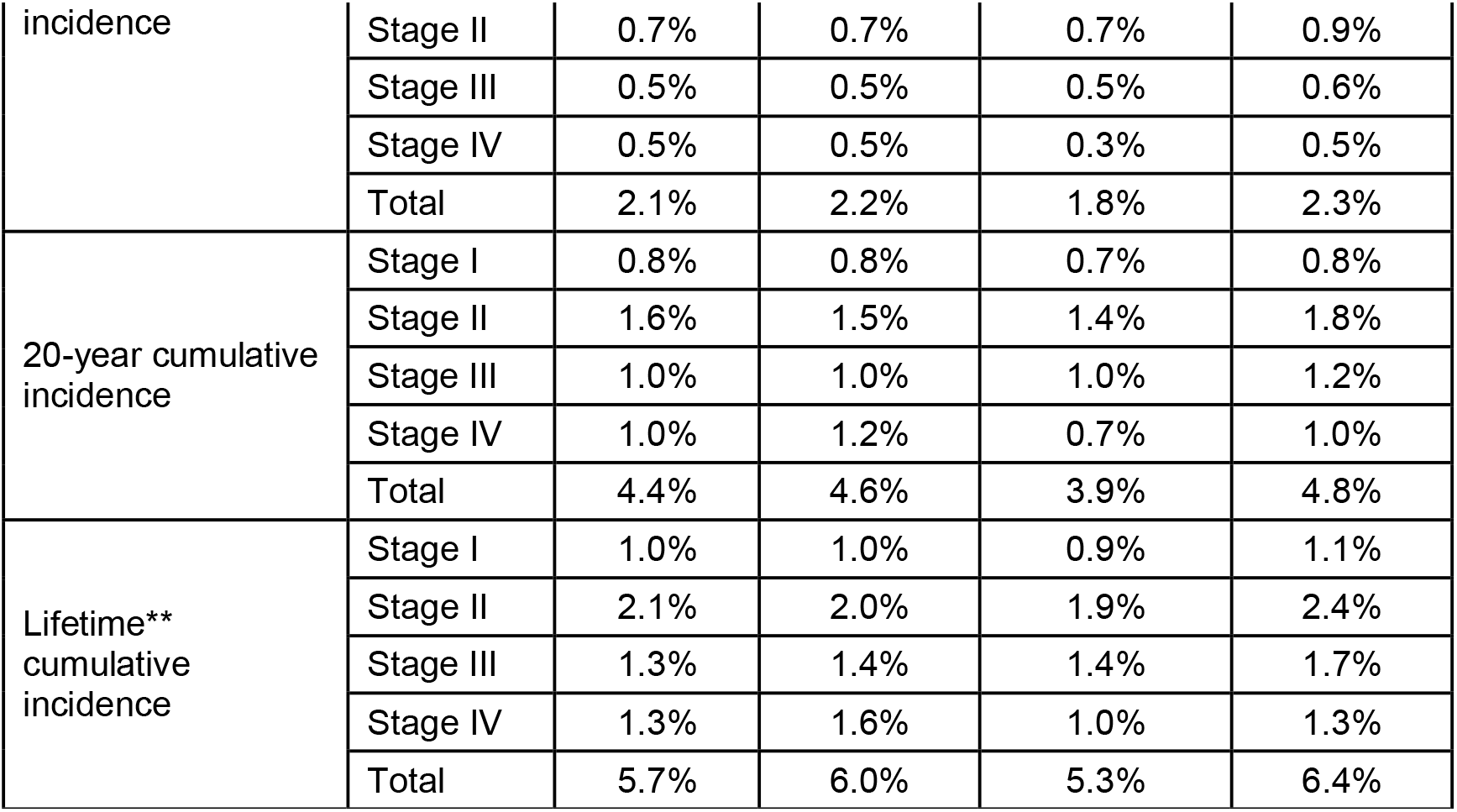
Prevalence and incidence of adenoma and colorectal cancer (CRC) by model. Multiplicity of adenomas data adapted from Kuntz et al^12^; other data adapted from Knudsen et al.^21^ *The table from Knudsen et al^21^ labels this category as “1-10 mm”. **The maximum lifespan age was not explicitly mentioned in Knudsen et al.^21^ CRC-AIM simulation stop-age was 120 years.

There was a comparable percentage of CRCs among CRC-AIM, CRC-SPIN, and SimCRC that developed from adenomas generated within 10 and 20 years of the clinical CRC diagnosis (**Figure S1**, **Table 2**). Although CRC-AIM’s overall dwell time is longer than the ranges reported for CRC-SPIN and SimCRC by Kuntz et al,^12^ it is less than another published estimate for CRC-SPIN^22^ (**Figure 3, Table S2)**. CRC-AIM’s derived annual transition probabilities align closely with those of CRC-SPIN (**Table S3**), except for preclinical CRC transition probabilities.

**Figure 3.**
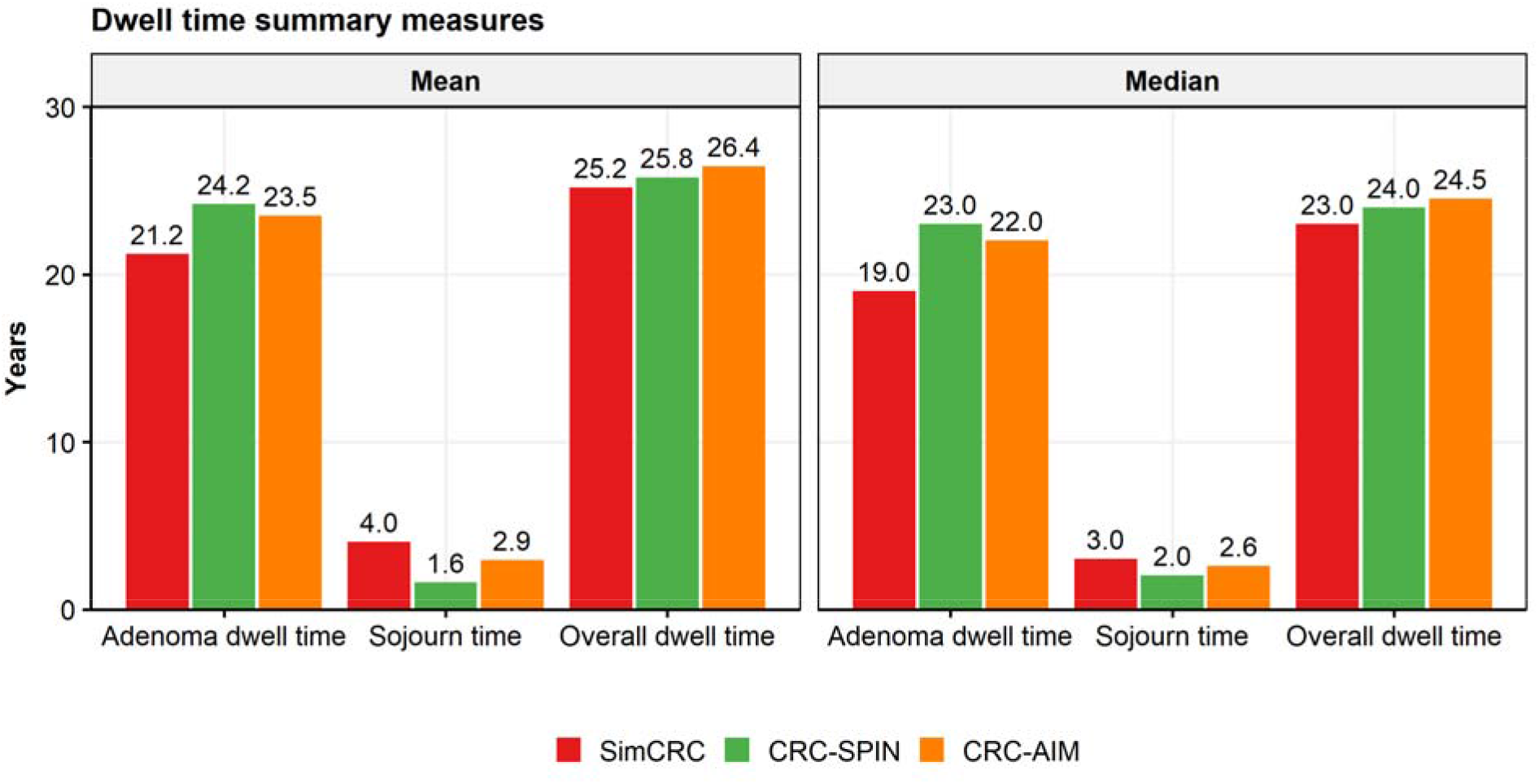
Median and mean adenoma dwell time, sojourn time, and overall dwell time in SimCRC, CRC-SPIN, and CRC-AIM. Data adapted from Kuntz et al;^12^ results from MISCAN are not included as the model has been subsequently recalibrated.

**Table 2.** Screening test characteristic inputs. Reproduced and adapted with permission from Knudsen et al, 2016.^2^ CRC, colorectal cancer.

| Screening Test | Value | Source |
| --- | --- | --- |
| **Colonoscopy (within reach, per lesion)** |  |  |
| Specificity | 86%<sup>b</sup> | (2013) Schroy et al<sup>28</sup> |
| Sensitivity for adenomas 1-5 mm | 75% | (2006) Van Rijn et al<sup>29</sup> |
| Sensitivity for adenomas 6-9 mm | 85% | (2006) Van Rijn et al<sup>29</sup> |
| Sensitivity for adenomas ≥10 mm | 95% | (2006) Van Rijn et al<sup>29</sup> |
| Sensitivity for CRC | 95% | Assumed |
| Reach | 95% (to end of cecum, remainder between rectum and cecum)<sup>c</sup> | Assumed |
| Risk of complications (serious/other gastrointestinal and cardiovascular) | age-specific risks<sup>d</sup> | (2014) Van Hees et al<sup>19</sup>;<br>(2009) Warren et al<sup>17</sup>;<br>(2003) Gatto et al<sup>18</sup> |
| **Fecal immunochemical test (per person)** |  |  |
| Specificity | 96.5% | (2014) Imperiale et al<sup>30</sup> |
| Sensitivity for adenomas 1-5 mm | 7.6%<sup>e</sup> | (2014) Imperiale et al<sup>30</sup> |
| Sensitivity for adenomas 6-9 mm |  | (2014) Imperiale et al<sup>30</sup> |
| Sensitivity for adenomas ≥10 mm | 23.8%<sup>f</sup> | (2014) Imperiale et al<sup>30</sup> |
| Sensitivity for CRC | 73.8% | (2014) Imperiale et al<sup>30</sup> |
| Reach | whole colorectum | Assumed |
| Risk of complications | 0% | (2016) Lin et al<sup>31</sup> |
| **High sensitivity guaiac based fecal occult blood test (per person)** |  |  |
| Specificity | 92.5% | (2008) Zauber et al<sup>32</sup> |
| Sensitivity for adenomas 1-5 mm | 7.5%<sup>g</sup> | (2008) Zauber et al<sup>32</sup> |
| Sensitivity for adenomas 6-9 mm | 12.4% | (2008) Zauber et al<sup>32</sup> |
| Sensitivity for adenomas ≥10 mm | 23.9% | (2008) Zauber et al<sup>32</sup> |
| Sensitivity for CRC | 70% | (2008) Zauber et al<sup>32</sup> |
| Reach | whole colorectum | Assumed |
| Risk of complications | 0% | (2016) Lin et al<sup>31</sup> |
| **Multi-target stool DNA (per person)** |  |  |
| Specificity | 89.8% | (2014) Imperiale et al<sup>30</sup> |
| Sensitivity for adenomas 1-5 mm | 17.2%<sup>e</sup> | (2014) Imperiale et al<sup>30</sup> |
| Sensitivity for adenomas 6-9 mm |  | (2014) Imperiale et al<sup>30</sup> |
| Sensitivity for adenomas ≥10 mm | 42.4%<sup>f</sup> | (2014) Imperiale et al<sup>30</sup> |
| Sensitivity for CRC | 92.3% | (2014) Imperiale et al<sup>30</sup> |
| Reach | Whole colorectum | Assumed |
| Risk of complications | 0% | (2016) Lin et al<sup>31</sup> |

| **SIG Flexible sigmoidoscopy (within reach, per lesion)** |  |  |
| --- | --- | --- |
| Specificity | 87%<sup>b</sup> | (2005) Weissfeld et al<sup>33</sup> |
| Sensitivity for adenomas 1-5 mm | 75% | Assumed |
| Sensitivity for adenomas 6-9 mm | 85% | Assumed |
| Sensitivity for adenomas ≥10 mm | 95% | Assumed |
| Sensitivity for CRC | 95% | Assumed |
| Reach | 76-88% to sigmoid-descending junction; 0% beyond splenic flexure | (2003) Atkin et al<sup>15</sup>;<br>(1999) Painter et al<sup>16</sup> |
| Risk of complications | 0% | Assumed<sup>h</sup>;<br>(2014) van Hees et al<sup>19</sup>;<br>(2009) Warren et al<sup>17</sup> |
| **CTC Computed tomographic colonography (CTC) (per lesion)** |  |  |
| Specificity | 88%<sup>i</sup> | (2008) Johnson et al<sup>34</sup> |
| Sensitivity for adenomas 1-5 mm | NPnot provided (only individuals with ≥6 mm lesions are determined to have positive CTC tests) | (2008) Johnson et al<sup>34</sup> |
| Sensitivity for adenomas 6-9 mm | 57% | (2008) Johnson et al<sup>34</sup> |
| Sensitivity for adenomas ≥10 mm | 84% | (2008) Johnson et al<sup>34</sup> |
| Sensitivity for CRC | 84% | (2008) Johnson et al<sup>34</sup> |
| Reach | Whole colorectum | Assumed |
| Risk of complications | 0% | (2016) Lin et al<sup>31</sup> |

**Table 3.** Percent of adenomas that had developed within 10 years or 20 years of clinical colorectal cancer diagnosis by model. Data adapted from Kuntz et al.^12^

| Age at cancer diagnosis (years) | Adenomas developed (%) |  |  |  |
| --- | --- | --- | --- | --- |
|  | MISCAN | CRC-SPIN | SimCRC | CRC-AIM |
| Within 10 years of clinical cancer diagnosis |  |  |  |  |
| 55 | 72% | 3% | 10% | 6.5% |
| 65 | 67% | 4% | 9% | 6.7% |
| 75 | 62% | 4% | 9% | 7.5% |
| Within 20 years of clinical cancer diagnosis |  |  |  |  |
| 55 | 94% | 24% | 39% | 42.2% |
| 65 | 92% | 25% | 37% | 37.5% |
| 75 | 89% | 28% | 33% | 36.2% |

**Table 4.** Screening strategies evaluated at perfect adherence. COL, colonoscopy; CTC, computed tomographic colonography; FIT, fecal immunochemical test; HSgFOBT, high sensitivity guaiac based fecal occult blood test; mt-sDNA, multi-target stool DNA test; SIG, flexible sigmoidoscopy

| Screening Modality | Screening Interval, y | Age to Begin Screening, y | Age to End Screening, y | No. of (Unique) Strategies |
| --- | --- | --- | --- | --- |
| no screening | NA | NA | NA | 1 (1) |
| COL | 5, 10, 15 | 45, 50, 55 | 75, 80, 85 | 27 (20) |
| mt-sDNA | 1, 3, 5 | 45, 50, 55 | 75, 80, 85 | 27 (27) |
| FIT | 1, 2, 3 | 45, 50, 55 | 75, 80, 85 | 27 (27) |
| HSgFOBT | 1, 2, 3 | 45, 50, 55 | 75, 80, 85 | 27 (27) |
| SIG | 5, 10 | 45, 50, 55 | 75, 80, 85 | 18 (15) |
| SIG and FIT (SIG_FIT) | 5_2, 5_3, 10_1, 10_2 | 45, 50, 55 | 75, 80, 85 | 36 (36) |
| SIG and HSgFOBT (SIG_HSgFOBT) | 5_2, 5_3, 10_1, 10_2 | 45, 50, 55 | 75, 80, 85 | 36 (36) |
| CTC | 5, 10 | 45, 50, 55 | 75, 80, 85 | 18 (15) |

Second, we observed consistent natural history outputs between CRC-AIM and the CISNET-based outputs described in the CISNET CRC technical report.^14^ The prevalence of preclinical CRC estimated by CRC-AIM is within the range of the CISNET models (**Figure 4A**). Because preclinical cancer prevalence is sensitive to sojourn time,^14^ if CRC-AIM used the original sojourn time parameter estimates from CRC-SPIN,^7^ then CRC-AIM’s prevalence would align with the prevalence reported by Berg et al^23^ (see **Supplemental Material**).

**Figure 4.**
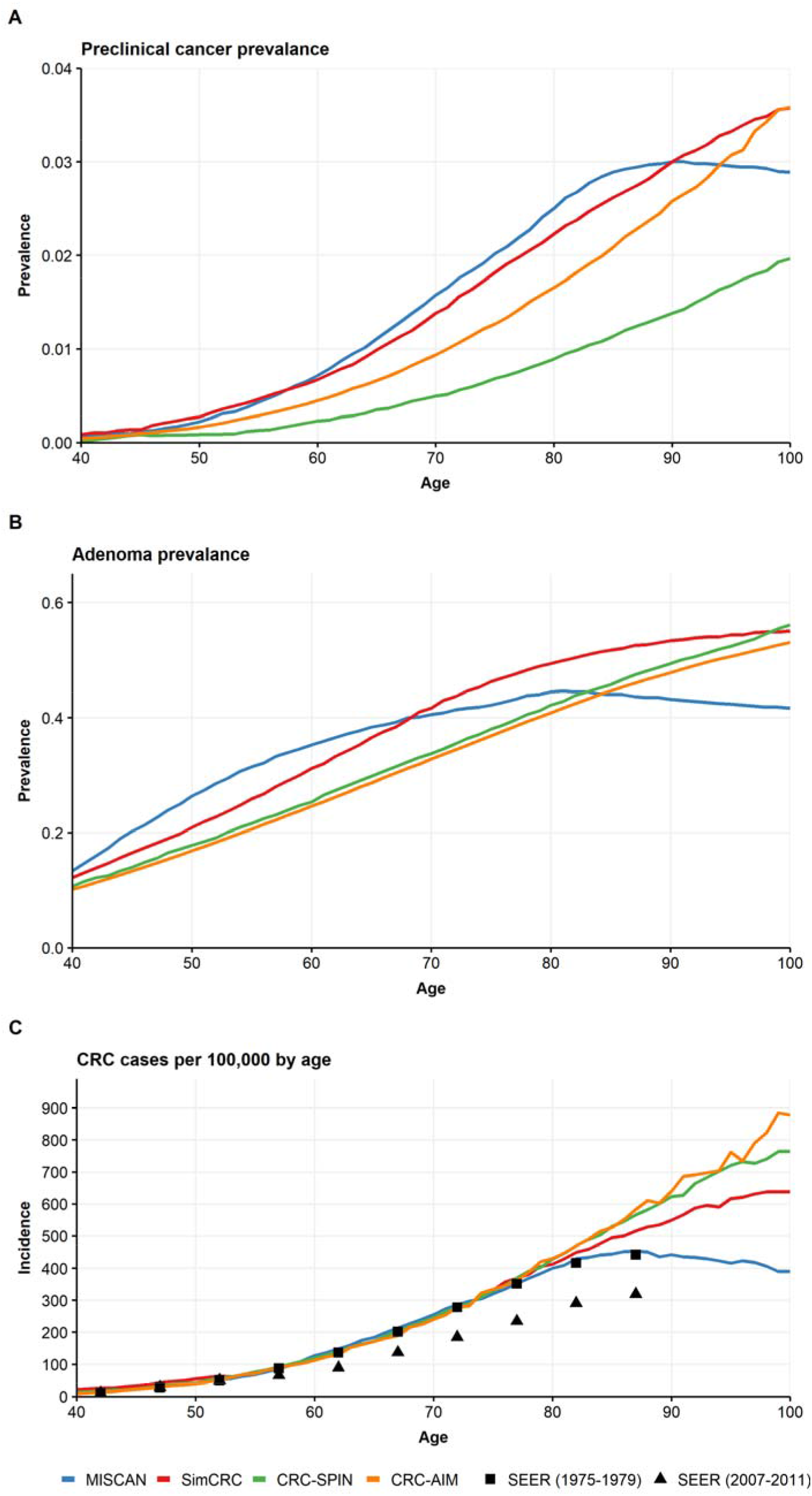
Preclinical cancer prevalence, adenoma prevalence, and colorectal cancer (CRC) incidence by model. Prevalence of (A) preclinical cancers and (B) adenoma by age. (C) Incidence of clinically diagnosed CRC per 100,000 individuals by age. Black squares represent CRC incidence for largely unscreened population according to the Surveillance, Epidemiology, and End Results data (SEER 1975-1979); black triangles represent CRC incidence with majority of screening-adherent individuals (SEER 2007-2011). Data adapted from Zauber et al.^14^

CRC-AIM’s adenoma prevalence almost overlaps that of CRC-SPIN (**Figure 4B, Table S4**). Both CRC-AIM and CRC-SPIN estimate lower adenoma prevalence compared to SimCRC and MISCAN, until approximately age 80 (**Figure 4B, Table S4**). The CRC-AIM CRC incidence curve overlaps the three CISNET models and 1975-1979 SEER estimates, trending more closely with CRC-SPIN than SimCRC or MISCAN after age 85 (**Figure 4C**). The distribution of adenoma location in CRC-AIM is identical to that of CRC-SPIN (**Figure S2**), and the distribution of adenomas by size of the most advanced adenoma among individuals aged 40, 60, and 80 is similar to CRC-SPIN (**Figure 5**). The CRC-AIM cancer stage distribution is almost identical to CRC-SPIN, with both models producing lower Stage IV cancer estimates compared to SimCRC and MISCAN due to a similar method to assign cancer stage probability^1^ (**Figure 6**).

**Figure 5.**
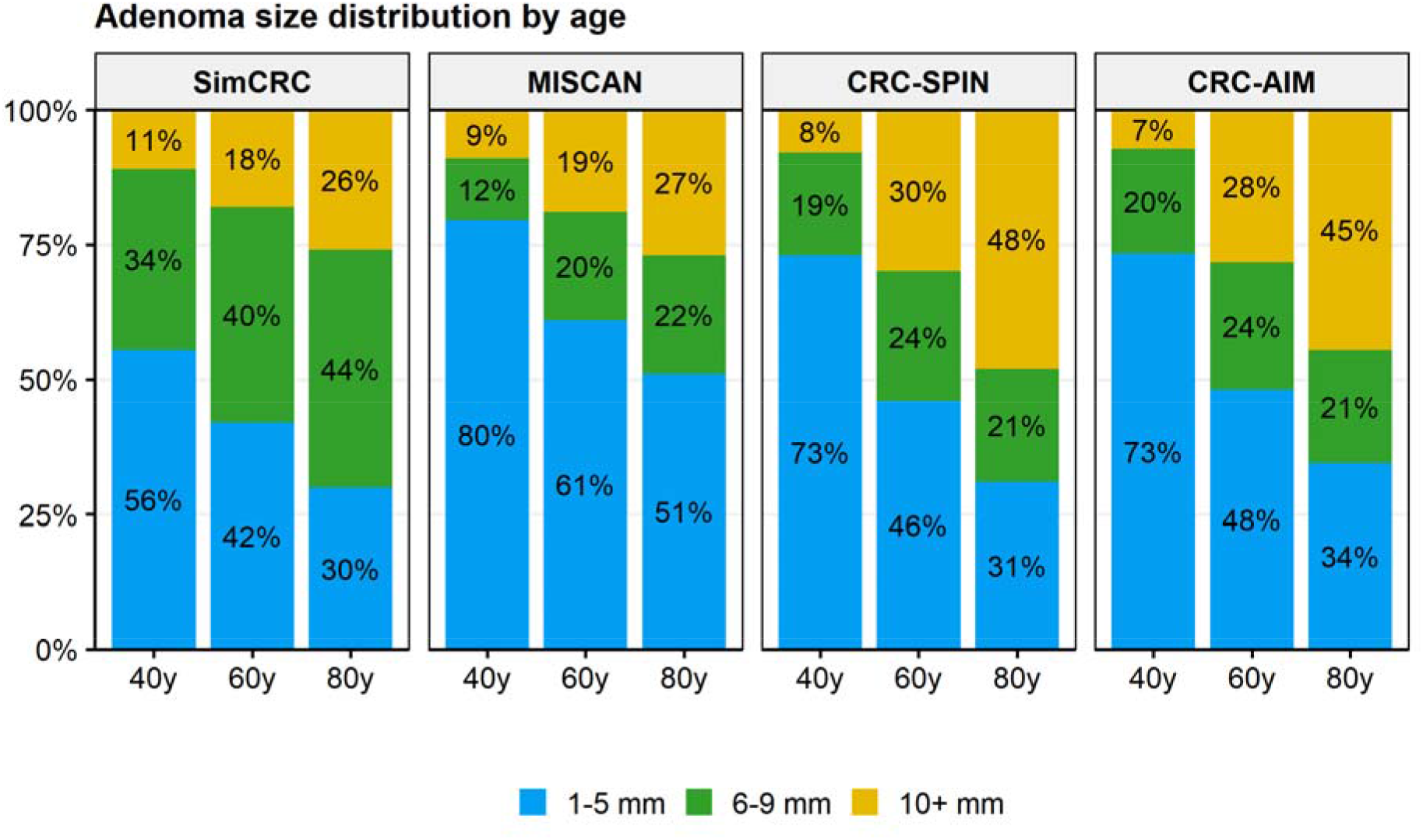
Adenoma size distribution by age of the most advanced adenoma by model. Data adapted from Zauber et al.^14^

**Figure 6.**
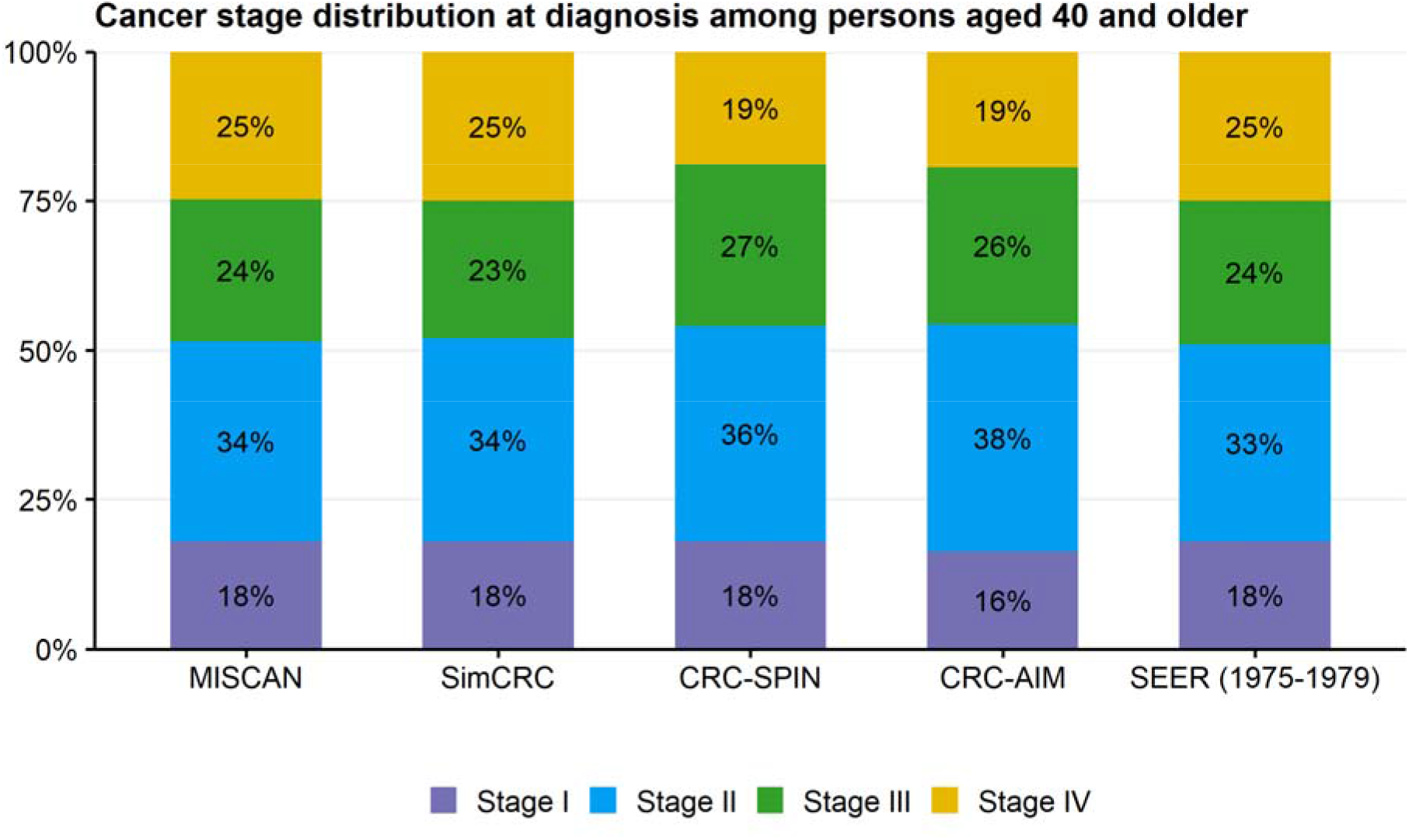
Colorectal cancer (CRC) stage distribution at clinical diagnosis among individuals aged 40 or older by model. Natural history stage distribution in the absence of screening is represented by Surveillance, Epidemiology, and End Results data (“SEER 1975-1979”). Stage was defined according to the American Joint Committee on Cancer (AJCC) staging system. Data adapted from Zauber et al.^14^

Additionally, we used CRC-AIM to explore the sensitivity of natural history modeling outputs (cumulative probability of developing CRC, cumulative risk of dying from CRC, and life expectancy) in an unscreened population based on the choice of life tables (cohort versus period; see **Methods** for details) for non-CRC-related mortality. CRC-AIM uses cohort life tables, which are preferred over period life tables for predicting future mortality.^8,9^ Although the CISNET models use period life tables, multiple natural history outputs between CRC-AIM and the CISNET models are generally comparable, with the exceptions of cumulative CRC risk and CRC mortality.

The curves describing the cumulative probability of developing CRC between ages 40 and 100 are similar between default CRC-AIM (with cohort life tables) and the CISNET models until age 80, after which CRC-AIM estimates more CRC (**Figure 7**). The cumulative probability is 8.0% for default CRC-AIM compared to 7.2% for CRC-SPIN, 7.0% for SimCRC, and 6.7% for MISCAN. Using period life tables, CRC-AIM generates an overlapping probability curve and a comparable cumulative probability of 7.1% (**Figure 7**). Similarly, the cumulative probability curve of dying from CRC between ages 40 and 100 is nearly identical to the CISNET models until age 80, after which CRC-AIM estimates more CRC deaths (**Figure 7**). The cumulative probability of dying from CRC is 3.2% in CRC-AIM using period tables and 3.7% in CRC-AIM using cohort tables, compared to 2.7% for CRC-SPIN and 2.8% for SimCRC and MISCAN.^14^For life expectancy, we replicated the CISNET model analysis by using period tables and assuming a simulation stop-age of 100 years. The life expectancy among 40-year-olds was 39.63 years for the modified CRC-AIM, which was almost identical to the CISNET models (39.6 years for SimCRC, 40.0 years for both MISCAN and CRC-SPIN). All three CISNET models simulate out to 100 years due to the unreliability of period life table estimates after age 100. Using the default cohort life tables, CRC-AIM simulated a life expectancy among 40-year-olds of 41.1 years, assuming a stop-age of 100 years. Because cohort life tables allow for the simulation of stop-ages beyond 100 years, we evaluated the simulated life expectancy for CRC-AIM given an assumed stop-age of 120 years, which was 41.7 years.

**Figure 7.**
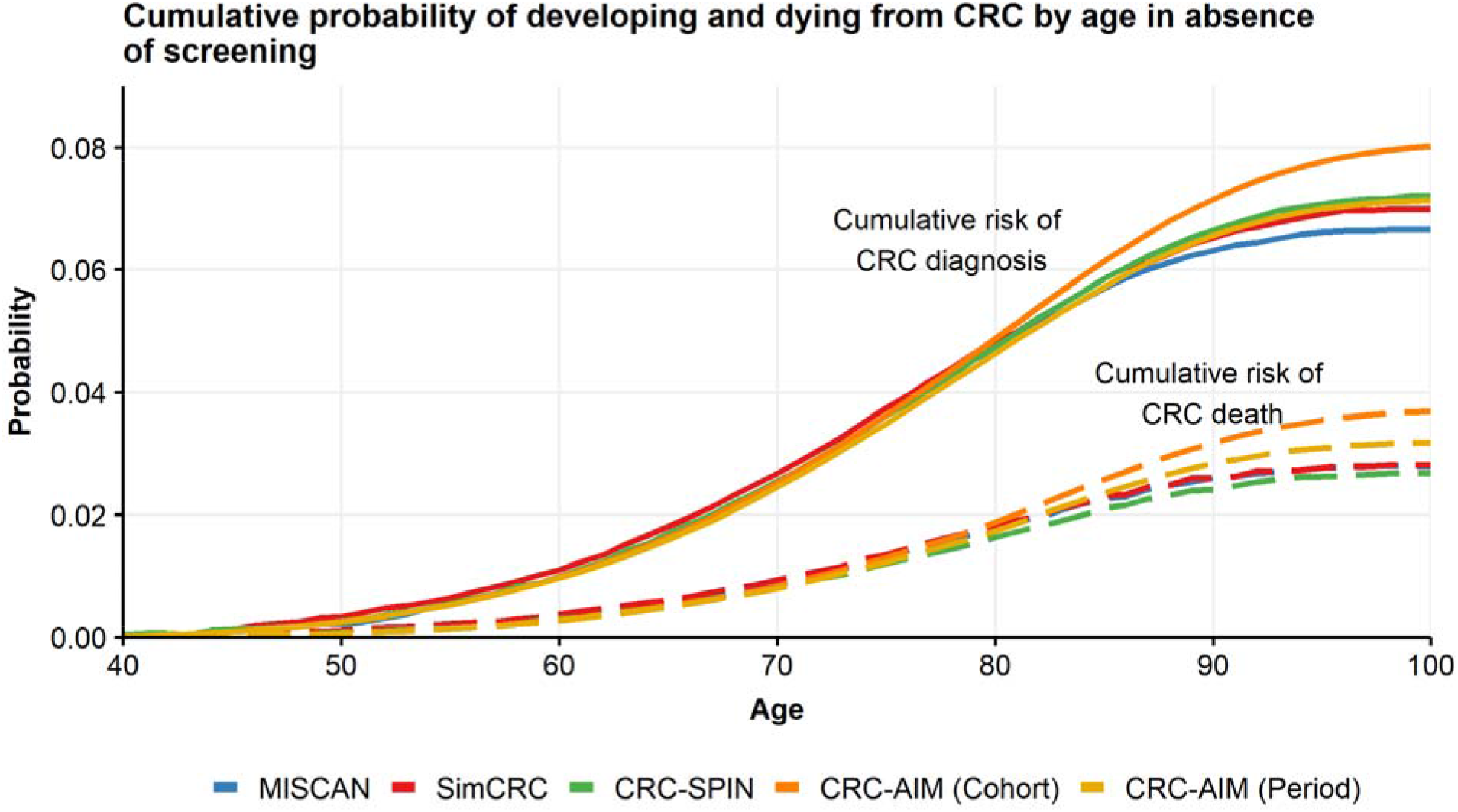
Cumulative probability of developing and dying from colorectal cancer (CRC) in the absence of screening by model. CRC-AIM results are displayed either using cohort life tables (“Cohort”), used standardly in CRC-AIM, or period life tables (“Period”), used in MISCAN, SimCRC, and CRC-SPIN, and for direct comparison in CRC-AIM. Data adapted from Zauber et al.^14^

Finally, we replicated the analysis from Kuntz et al^12^ where cumulative probability of developing CRC was considered purely as a function of the adenoma-carcinoma sequence and the competing risk of mortality is eliminated (ie, removing life tables from consideration). For the 20-year cumulative CRC incidence (at age 75) for individuals with underlying lesions and without competing mortality, CRC-AIM projected 13.5%, which is comparable to the CISNET models (13.5% for CRC-SPIN, 13.1% for SimCRC, and 8.6% for MISCAN) (**Figure 8**). Similar results were obtained for the risk ratios at age 75 for the group of individuals with lesions (at age 55) compared to the group without underlying lesions (at age 55)—CRC-AIM projected a risk ratio of 37.3, within the range of the CISNET models (75 for CRC-SPIN, 29 for SimCRC, and 7 for MISCAN) (**Figure 8**).

**Figure 8.**
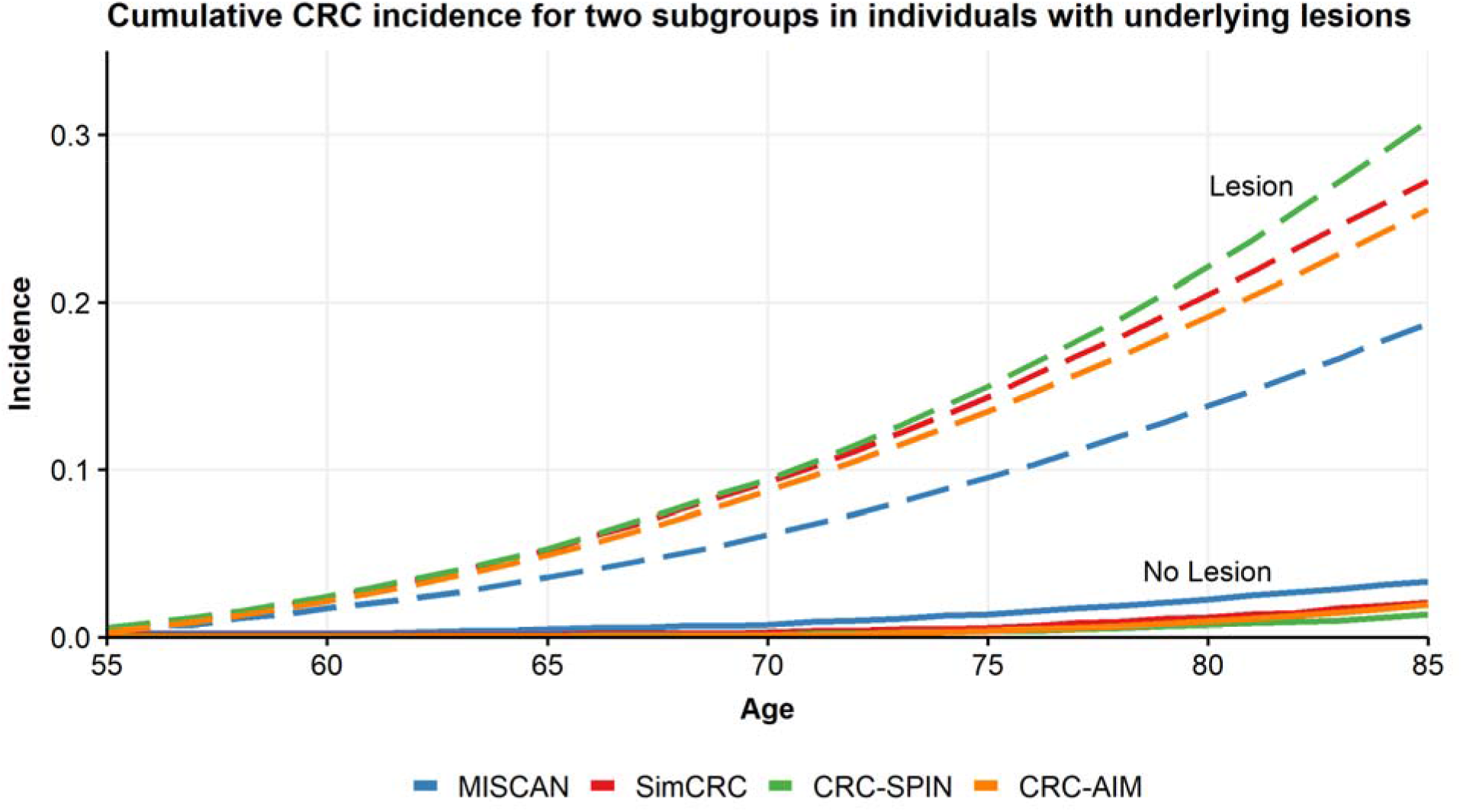
Cumulative CRC incidence for two subgroups in individuals with underlying lesions by model. The subgroups include individuals with or without an adenoma or preclinical cancer at age 55 (“Lesion” or “No Lesion”, respectively). Data from Kuntz et al.^12^

The screening overlay validation analysis assumed perfect adherence for CRC screening strategies. We generated screening overlay tables from CISNET publication (**Tables S36-43**) using CRC-AIM with period life tables. When no screening was conducted, the CRC-AIM model estimated that 71.1 out of 1000 individuals free of CRC at age 40 would be diagnosed with CRC during their lifetime and 31.7 would die of CRC (**Table S36**). When screening was conducted, all strategies provided clinical benefit in terms of LYG and reductions in CRC-related incidence and mortality (**Tables S36-43**). With those tables, we were able to compute the efficient frontiers (**Figure 9** and **Tables S44-45**).

**Figure 9.**
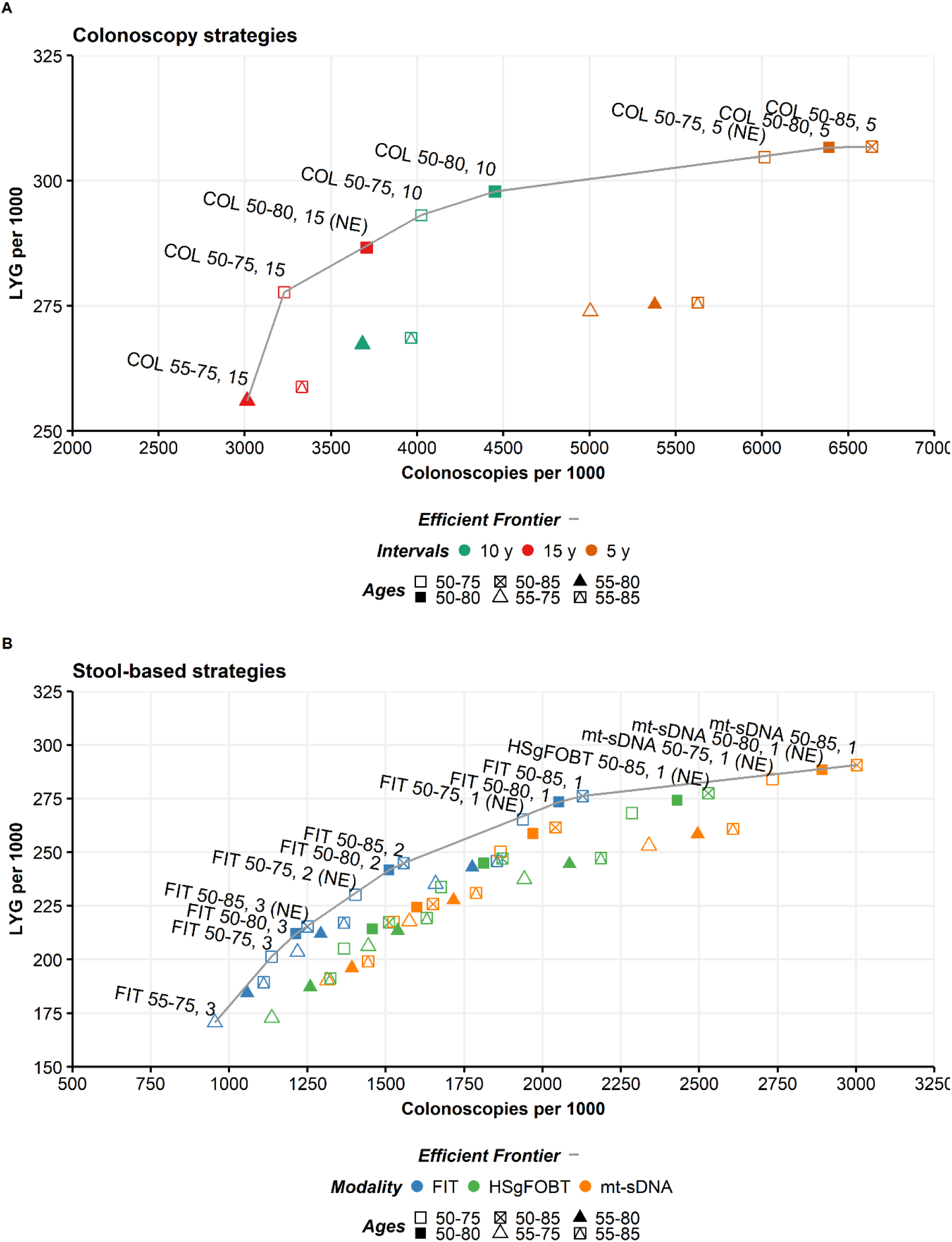
Efficient frontier of screening strategies for individuals aged 40 years using (A) colonoscopy and (B) stool-based tests. Models conformed to the CISNET assumption of perfect adherence. FIT, fecal immunochemical test; HSgFOBT, high sensitivity guaiac based fecal occult blood test; mt-sDNA, multi-target stool DNA. NE, near-efficient.

We conducted three analyses to demonstrate model cross-validation (1) quantitative method comparison, in which screen-related outcomes from all models were compared, using Passing-Bablok regression, in terms of systemic bias, proportional bias, and total bias at the low and high measurement range; (2) qualitative efficient frontier comparison, in which the efficient frontiers from all models were compared using concordance analysis; (3) medical decision making comparison, in which the final recommended screening strategies from all models were compared (see **Supplemental Material**).

Overall, there is substantial overlap across multiple natural history and screening overlay outputs between CRC-AIM and the CISNET models, especially CRC-SPIN.

## Discussion

We developed a robust natural history model of colorectal cancer—the Colorectal Cancer and Adenoma Incidence & Mortality (CRC-AIM) microsimulation model—to facilitate greater opportunities and efficiencies in collaborative modeling research. We compared the results of CRC-AIM with published results for the three CISNET CRC models: CRC-SPIN, SimCRC, and MISCAN.

The percentage of adenomas that had developed within 10 or 20 years prior to clinical CRC diagnosis is similar between CRC-AIM and the CISNET models (**Figure S1**, **Table 2**). This similarity suggests a comparable interaction between screening interval and CRC incidence reduction. If the adenoma transition rates are similar, then the screening benefits will be comparable because all models will have similar windows of opportunity for a hypothetical screen to detect and remove CRC-causing adenomas.^12^ However, if there is a difference in the proportion of adenomas that become cancerous within a particular timeframe, then the models could have different reductions in CRC incidence and mortality, depending on the screening modality.^12^ Because of this similarity, CRC-AIM demonstrated relative comparability with the CISNET models in terms of CRC incidence and mortality reduction and conclusions related to medical decision making.

Although the natural history outputs from CRC-AIM generally aligned with the CISNET models, there were some notable differences. CRC-AIM’s derived annual transition probabilities align closely with those of CRC-SPIN (**Table S3**), except for preclinical CRC transition probabilities, which is likely caused by specifying a different sojourn time (ST) for CRC-AIM (**Table S2**; see **Supplemental Material**). When the original CRC-SPIN parameter estimates are used for ST, the preclinical CRC transition probabilities are similar between CRC-AIM and CRC-SPIN (data not shown).

Minor differences in cumulative cancer mortality by age 100 (natural history) and life-years-gained (screening overlay) are likely due to a different method for calculating CRC survival between CRC-AIM (cause-specific survival) and the CISNET models (relative survival). The outcomes between relative survival and cause-specific survival are generally comparable; although relative survival is more commonly used with registry data, cause-specific survival benefits from enhanced flexibility and can better incorporate risk factors for particular populations.^24,25^ Because the CISNET models use identical CRC-based survival functions,^14,20,26,27^ the potential inter-model variability that would have occurred if each group had independently developed their own survival functions is unknown. Since CRC-AIM’s cause-specific survival functions overlap considerably with the CISNET models until age 85, after which only minor deviations are observed, we predict that CRC-AIM’s cumulative cancer mortality would lie within an inter-model variability.

In our analysis, we compared the impact of cohort versus period life tables on the sensitivity of natural history outputs in an unscreened population. CRC-AIM uses cohort life tables, which determine annual survival from past mortality rates and/or projected mortality rates. The CISNET models use period life tables, which describe what would happen to a hypothetical cohort if it experienced, throughout its entire life, the mortality conditions of a particular time period. In general, cohort tables are preferred over period life tables for predicting future mortality,^8,9^ and the latter may underestimate survival because they only represent a snapshot of the current mortality experience.^8,9^

We demonstrate that the choice of life table impacts CRC incidence and mortality, particularly after age 85. The differences in cumulative probability of developing CRC after age 85 in CRC-AIM compared to the CISNET models is primarily driven by using cohort life tables—older individuals are alive and at risk of developing CRC. Consequently, this impacts cumulative mortality, because more individuals will die of CRC if more individuals develop CRC. We found that choice of life table did not significantly impact other modeling outputs.

In general, by conforming to CISNET’s assumptions regarding their efficient frontier calculations, we reproduced their overall screening outputs (**Figure 9**). We also conducted multiple model cross-validation comparisons between CRC-AIM and the CISNET models. For comparisons based on quantitative outcomes, qualitative efficient frontiers, and overall strategy recommendations, we found that the differences in outcomes between CRC-AIM and the CISNET models were generally similar to the differences among the CISNET models themselves (see **Supplemental Materials**). Quantitative differences for CRC-AIM versus the CISNET models were observed with total surveillance colonoscopies, total colonoscopies, and life-years-gained, depending on the comparison that was performed. Regardless, qualitative decisions related to inform clinical practice in terms of screening modality using CRC-AIM are almost identical to those made by the CISNET modelsOne limitation of our analyses is that were limited to the descriptions of model parameters and assumptions found in the CISNET model publications, which we replicated as closely as possible. These models have likely been altered over time and some analytical details are not provided within the publications.

With CRC-AIM, we want to stimulate community engagement and enhance research into underexplored questions in population health by clearly articulating its assumptions and framework. To that end, we have deposited CRC-AIM’s formulas, parameters, and additional documentation on a publicly accessible repository (https://github.com/CRCAIM/CRC-AIM-Public). (This information is also included in Table S5 and Supplemental Material.) In addition, we will make CRC-AIM’s underlying Python code available to collaborators to facilitate information-sharing and emphasize visibility and transparency. We have included additional details regarding opportunities for collaboration on the CRC-AIM repository.

## Supporting information

Supplemental Material - Analysis

## Acknowledgments

Medical writing and editorial/data visualization support were provided by David K Edwards V, PhD, with assistance from Rebecca Swartz, PhD (both of Exact Sciences, Madison, WI) and Erin P Scott, PhD (Maple Health Group, LLC; funded by Exact Sciences).

We would like to thank A Burak Ozbay, PhD (Exact Sciences) and Marcus Parton (Exact Sciences) for helpful guidance and overall feedback; David Young, MS (Auburn University) and Gennadiy Gorelik, PhD (EmpiriQA) for technical advice; and Evan Musick (Exact Sciences) for assistance with data visualization.

## Permissions

Permissions will be obtained from the publishers as needed.

