## Supplemental Material - Text for "Description and Validation of the Colorectal Cancer and Adenoma Incidence & Mortality (CRC-AIM) Microsimulation Model"

**Introduction**

This supplemental section provides additional information about the development and validation of the Colorectal Cancer and Adenoma Incidence & Mortality (CRC-AIM) microsimulation model. First, we describe the predominant differences between CRC-AIM and CRC-SPIN—namely, the differences in overall approach and calibration. Next, we discuss the recalibration process for CRC-AIM, including presenting outputs from a prototype version of the model before the recalibration was conducted. We provide greater detail into the methodological details of CRC-AIM, especially for components for which sufficiently detailed formulas and parameter descriptions from CRC-SPIN were not published. For cause-specific survival, we include comprehensive supplemental tables and figures that describe the fitting and selection from among five different parametric linear regression models for colorectal cancers (CRCs) based on American Joint Committee on Cancer (AJCC) stage and location. We describe how we generated a multinomial logistic regression to determine CRC stage based on size and provide extensive descriptions of our corresponding query of the Surveillance, Epidemiology, and End Results (SEER) dataset so that our methodology can be replicated. We detail our process of determining the size distribution of clinically detectable CRC, again with descriptions of the corresponding SEER query. We include comprehensive supplemental tables describing the screening overlay outputs similar to those provided by CISNET in their technical report^1^ and the USPSTF outputs.^2^ Finally, we describe the results of three comparisons to demonstrate model cross-validation—a quantitative method comparison, qualitative efficiency frontier comparison, and medical decision making comparison—in which CRC-AIM demonstrated similar outputs based on CISNET modeling assumptions.

Furthermore, to emphasize transparency in our methodology and parsimony for subsequent publications, we created a publicly available repository for CRC-AIM’s formulas and parameters (<https://github.com/CRCAIM/CRC-AIM-Public>). The repository also will list all subsequent scientific presentations or publications related to CRC-AIM. We will link versions of the natural history model and/or other model components to each presentation and publication over time to maintain version control. We will update the repository periodically as various components of CRC-AIM are added or improved.

We are excited to collaborate with researchers and members of the modeling community to help address outstanding questions with CRC-AIM related to CRC screening and early detection. If you are interested in working with us, please email us at.

**Notable Differences between CRC-AIM and CRC-SPIN**

There are two fundamental differences between CRC-AIM and CRC-SPIN.

The first is in overall approach. CRC-SPIN is a continuous-time microsimulation model, in which the natural history processes that describe an individual’s adenoma-carcinoma sequence are generally expressed as continuous cumulative distribution functions (CDFs). As a result, CRC-SPIN generates events in an individual’s life that can be precisely expressed within an arbitrarily small unit of time.

In contrast, CRC-AIM features a cycle-based approach, in which event probabilities—namely adenoma generation and transition into preclinical cancer—are assigned at the beginning of a cycle and depend on the pre-defined cycle length. This approach is commonly used to develop microsimulation models (eg, by TreeAge Pro, a popular modeling software) and reflects more classic, canonical ways of structuring and considering microsimulations. A cycle-based approach can facilitate collaboration among clinicians and researchers who do not have a substantial background in programming. One consequence is that it does not allow for either the generation of multiple adenomas in the same year or an individual transitioning across “health states” in the same year (eg, an adenoma transitioning to both preclinical and clinical CRC). However, the probability of multiple health state transitions occurring within the same year has been quantified in a CRC-SPIN publication and was found to be exceedingly small.^3^

The second difference relates to microsimulation model calibration. Calibration is the process of deriving model parameters that allow the model to replicate expected or observed data, referred to as calibration targets.^3^ The CRC-SPIN model uses a Bayesian calibration method, which offers two advantages over non-Bayesian methods^3^: (1) posterior model parameters can be explicitly influenced by prior knowledge such as expert opinion or empirical data, which is incorporated as prior distributions; and (2) the uncertainty of modeled outcomes can be quantified as prediction intervals by sampling across the joint posterior distribution of model parameters.

Calibration may be implemented as an automated statistical procedure to update a model’s parameters and can be performed for various reasons, including incorporating new empirical evidence, modifying prior distribution(s) (for Bayesian models) and/or calibration targets, or updating poorly performing statistical distributions within the model (eg, for CRC-SPIN v2.x^4^). After calibration is performed, when a Bayesian microsimulation model is executed, parameters are sampled across the joint posterior distribution. This parameter sampling generates an estimate for each model parameter (23 for CRC-SPIN) and the natural history of individuals—often millions of individuals—are simulated. The parameters are then resampled and another set of individuals are simulated. This process is repeated for hundreds or thousands of samples of the joint posterior distribution. The average outcome across all draws is often reported as the point-prediction, and prediction intervals for the outcome can be estimated using percentile methods to indicate uncertainty in model predictions. The process is similar in concept to a probabilistic sensitivity analysis.

CRC-AIM does not implement a Bayesian calibration component, although we may incorporate a formal calibration component into future versions of the model. Bayesian calibration methods for complex microsimulations are computationally costly to perform and, once calibrated, Bayesian microsimulation models are computationally costly to execute. After we recognized that certain model parameters had to be recalibrated (see “CRC-AIM Recalibration”), we estimated a single set of these best-fitting parameters using a Gaussian process framework. This optimization framework can be suitable for computer simulations that yield little to no variation in outcome given the same set of inputs, as long as a sufficiently large simulation size is used. We selected particular natural history outcomes from CRC-SPIN as calibration targets to help ensure comparability between CRC-SPIN and CRC-AIM.

**CRC-AIM Recalibration**

To initially assess the performance of the model, we used the posterior means of the 23 calibrated parameters from CRC-SPIN that are described in Rutter et al.^5^ As detailed in the main manuscript, we compared the natural history outputs of CRC-AIM with publicly available outcomes from CRC-SPIN, which have been described in multiple publications over time. Through this process, we observed a high degree of comparability in almost every natural history output between both models. However, we want to describe some of the discrepancies we observed that required us to recalibrate CRC-AIM. Here, we describe these dissimilarities and the recalibration process in greater detail. For the complete list of parameters and their recalibrated estimates, see **Table S5**.

*Adenoma growth parameters*

We observed a discrepancy between CRC-AIM and CRC-SPIN in the distribution of the most-advanced adenoma size by age (**Figure S3**). Specifically, at age 40, CRC-AIM reported only 4% of adenomas were ≥10 mm, compared to 8% for CRC-SPIN. Because adenomas only begin to generate after age 20, we hypothesized that certain adenomas were growing too slowly after generation. We discovered colon-based adenomas were growing too slowly—for adenomas generated at ages less than 40, in our implementation of the model, it was improbable for them to become ≥10 mm by the age of 40.

However, the adenoma size distribution at age 65 was similar for CRC-AIM and CRC-SPIN, including for adenomas ≥10 mm (**Table S6**). Aside from differences in model implementation, there are at least two possible explanations for this discrepancy. One possibility is that after age 40, there is enough time for the adenomas to grow to match the size distribution from CRC-SPIN. Another possibility is that CRC-SPIN could have been recalibrated after this publication and the impact of a potential recalibration has not been fully described.

*Transition probabilities by location/sex*

We also observed that CRC-AIM was generating lower values for overall SEER cancer incidence compared to CRC-SPIN (**Figure S4A**), which also resulted in an underestimation of overall cumulative cancer risk (**Figure S4B**). (The recalibrated comparisons between CRC-AIM and the CISNET models can be found in **Figure 4C** and **Figure 7**, respectively, in the main manuscript.) Overall CRC incidence is a direct function of age-stratified CRC incidence by location and cancer site. In the prototype version of CRC-AIM, age-stratified cancer incidence was only slightly under-generated for colon cancers in males and females and rectal cancers in females—approximately 4-13% below expected results (**Table S7**). The primary reason for under-generated overall CRC incidence was rectal cancers in males, which were consistently 30% below expected results (**Table S7**).

We considered multiple potential explanations for this discrepancy. Our first consideration was sojourn time (ST), because CRC incidence is partly a function of ST: changing sojourn time will shift cancer incidence when all other model parameters are kept equal. However, the prototype version of CRC-AIM replicated the overall ST for CRC-SPIN (see **Table S2** for CRC-SPIN and **Table S8** for the CRC-AIM prototype), which was unsurprising because ST parameters for preclinical cancers are explicitly defined by formulas and therefore are tightly controlled. Additionally, ST is neither age- nor sex-specific in both CRC-SPIN^5^ and CRC-AIM, and while the rectal cancer incidence in males was greatly underestimated, the rectal cancer incidence in females was relatively comparable between models. Therefore, we concluded that ST did not likely contribute to this discrepancy.

Next, we considered other components that contribute to CRC incidence—namely, adenoma generation (both sex- and age-specific), adenoma growth, and adenoma transition parameters. Adenoma generation was comparable between CRC-SPIN and CRC-AIM (**Figure 4B** and **Table S4**). Although we had experienced discrepancies in the adenoma growth by location (see “Adenoma growth parameters”), adenoma growth is not sex-specific. In addition, the differences in CRC incidence for rectal cancer incidence in males occurred across all age groups, not just in individuals over 40 years old.

The only remaining parameters were those that enabled adenomas to transition to preclinical cancer—specifically, the adenoma size (γ_1_) and age of adenoma initiation (γ_2_) parameters. These parameters are highly sensitive, with minor variations having a large impact on cancer incidence. The minor yet consistent differences for age-stratified colon cancer incidence in males and females and rectal cancer incidence in females could simply have been due to the lack of precision in the published estimates of the gamma parameters estimates (reported only to the thousandths place) or due to differences in the way the programs are implemented. Using the published parameter estimates, we found that simple plots of the CDF for adenomas transitioning to preclinical cancer also indicated that rectal cancers in males would be underestimated. Those curves revealed that the rectal-male curves lie underneath the other location/sex curves across multiple ages of adenoma initiation (**Figure S5**), demonstrating that these parameters were contributing to the discrepancy in CRC incidence.

*Sojourn time*

We observed that the mean sojourn times (STs) of colon- and rectal-based cancers for CRC-SPIN (averaged to 1.9 years for 2010^3^ and 1.6 years for 2011^6^) was noticeably shorter than the published ST of MISCAN (3.0 years^6^) and SimCRC (4.0 years^6^) (**Table S2**). This discrepancy prompted a more thorough analysis into modeling ST, including estimates based on real-world evidence and model validation studies.

One study by Zheng and Rutter estimated sojourn time in a study of 42,079 patients who received fecal occult blood tests (FOBT).^7^ The estimated mean sojourn time (in years) for colon and rectal cancers, respectively, was determined to be 3.86 and 3.35 for 45- to 54-year-olds, 3.78 and 2.24 for 55- to 64-year-olds, and 2.70 and 2.10 for 65- to 74-year-olds,^7^ all values larger than the mean STs for CRC-SPIN. Another study evaluated the accuracy of the CISNET model outputs against outcomes from the United Kingdom Flexible Sigmoidoscopy Screening (UKFSS) trial.^8^ Notably, although CRC-SPIN did not accurately predict the number of screen-detected cancers, the two models with the longer sojourn times (MISCAN and SimCRC) did. Based on this observation, the authors concluded that the mean ST is probably between 1.6 and 4.0 years and that the actual value is on the higher end of that estimated range. Finally, Brenner et al used data from a German national screening database to estimate sex- and age-stratified mean sojourn times, which were determined to be between 4.5 and 5.8 years for the subgroups assessed.^9^

We determined that this evidence provided sufficient justification to extend the sojourn time for CRC-AIM beyond that of CRC-SPIN, whose STs for colon and rectal cancers are 1.9 and 2.7 years, respectively. Therefore, we conservatively extended the average sojourn by one year each (to 2.9 and 3.7 years, respectively) but maintained the standard deviation (µ_c_*τ_c_, µ_r_*τ_r_) of the distributions equivalent to CRC-SPIN (see **Tables S2** and **S5**).

We chose to be conservative in extending the sojourn time because preclinical cancer detection depends on both the interval length of a screening test and the ST length. If the ST length is shorter than a screening interval, then there is a possibility of no chance to detect a preclinical cancer since the development from preclinical to clinical CRC could fall entirely in the window between screening tests.^6^ However, if the ST is double that of a screening interval, then there are two chances for a screening test can detect the preclinical cancer. In this way, for cancers with a longer ST, a greater overall screening benefit could be conferred to patients who are screened with a test with a longer screening interval. We did not want to bias CRC-AIM to favor screening tests with longer intervals. We hypothesize that CRC-SPIN’s sojourn time will be updated through recalibration but the recalibrated parameters are unknown.

*Recalibration methodology*

After updating the model to include the evidence-based sojourn times as described above, we conducted a sequential recalibration of the discrepant parameters, first updating the adenoma growth parameters and then updating the transition to preclinical cancer parameters. In general, we used space-filling Latin hypercube sampling across the published posterior distribution estimates (Bayesian credible intervals) for the parameters.^3^ For adenoma growth parameters, we restricted combinations of β_1_/β_2_ to limit the probability of an adenoma reaching 10 mm within 10 years from 0.0001 to 0.25, similar to the CRC-SPIN v2.x recalibration.^4^

For the adenoma growth recalibration targets, we used a combination of the adenoma number and size distribution by age for CRC-SPIN (**Table 1**) and the adenoma size distribution for the most advanced adenoma for CRC-SPIN (**Figure 5**). For the transition probability recalibration targets, we used the 1975-1979 SEER and CRC-SPIN v1.0 observed cancer rates as lower and upper limit optimization targets.

Parameters were then optimized using a Kriging model approach. The resulting recalibrated parameters are reported in the main manuscript (see **Table S5** for a complete list of CRC-AIM’s parameters), and the outputs from recalibration are also reported.

**Additional Methodological Details**

*Adenoma generation*

CRC-AIM generates adenomas using the non-homogenous Poisson process from CRC-SPIN v1.0, specifically the instantaneous risk function, which has seven calibrated parameters (**Table S5**).^10^

Two parameters describe the per-person risk of developing an adenoma: a baseline log-risk parameter (*α_0_*) and a standard deviation (*σ_α_*) for baseline-log risk that are used in conjunction to sample a unique per-person risk (*α_0i_*), allowing heterogeneity in adenoma generation across individuals. One sex effect parameter (*α_1_*) alters the risk of developing an adenoma based on the sex of the individual. There are four age-group parameters (*α_20_, α_21_, α_22_, α_23_*) that alter adenoma risk based on age.^10^ The instantaneous risk function is defined as:

$$\psi_{i}\left( t \right)=exp\left( \alpha_{0i}+\alpha_{1}{sex}_{i}+\sum_{k=1}^{4} \delta\left( A_{\kappa}<{age}_{i}\left( t \right)\leq A_{\kappa+1} \right)\left\{ {age}_{i}\left( t \right)\alpha_{2\kappa}+\sum_{j=2}^{\kappa} A_{j}\left( \alpha_{2 j-1}-\alpha_{2j} \right) \right\} \right)$$

$$\alpha_{0i} \sim N\left( \alpha_{0,}\sigma_{\alpha} \right)$$

Notably, CRC-AIM does not use CRC-SPIN’s modification of this function for a continuous-time model^10^ because CRC-AIM uses a cycle-based approach.

*Adenoma growth*

CRC-AIM uses the adenoma growth function from CRC-SPIN v1.0, which uses two calibrated parameters that describe a CDF for a type II extreme value distribution defined as the time required for an adenoma to reach 10 mm in size. The CDF of time *t* to 10 mm is defined as:^10^

$$F\left( t \right)= exp\left( -\left( \frac{t}{\beta_{1}} \right)^{{-\beta}_{2}} \right)$$

where *β_1_* and *β_2_* are the location and scale parameters, respectively, and vary based on location of the adenoma (colon vs rectum).

The sampled time to reach 10 mm can be used to calculate the growth rate *λ* for the *j-*th adenoma in the *i-*th individual, which is defined as:^10^

$$\lambda_{ij}=\frac{-log\left( \frac{d_{\infty}-\text{10}}{d_{\infty}-d_{0}} \right)}{t_{10mm}}$$

The growth curve function describing the diameter (in mm) at time *t* is:^10^

$$d_{ij}\left( t \right)= d_{\infty} -(d_{\infty}-d_{0})e^{-\lambda_{ij}t}$$

where $d_{\infty}$ and $d_{0}$ are the maximum and minimum adenoma sizes, respectively.

The distribution of large (≥10 mm) adenomas generated using this method from CRC-SPIN appears to be larger than the distribution from SimCRC and MISCAN after age 40.^1^ For example, at age 60, 30% of the most advanced adenoma are ≥10 mm in CRC-SPIN compared to 18% and 19% for SimCRC and MISCAN, respectively.^1^ CRC-SPIN v2.x uses a different growth function, perhaps to correct for this observation.^11^ Because CRC-AIM uses the same adenoma growth methodology as CRC-SPIN v1.0, it reproduces the same large adenoma distribution as CRC-SPIN and replicates the discrepancy compared to SimCRC and MISCAN (**Figure 5**).

*Transition to clinically detectable CRC*

CRC-AIM uses a lognormal distribution similar to CRC-SPIN v1.0 to determine the sojourn time (ST), or the time required for a preclinical cancer to become clinically detectable.^10^ CRC-SPIN v1.0 models ST for the *i-*th individual’s *j-*th preclinical cancer using a lognormal distribution that is conditional on location (colon vs rectum).

If *t* represents ST, then log(*t*)~Normal(ξ_c_,ν_c_) for adenomas in the colon. This distribution is parameterized by CRC-SPIN using bracket notation in terms of mean and standard deviation: t~Log Normal [µ_c_,τ_c_μ_c_], where the mean of *t* is $\mu_{c}=\text{exp}\left( \xi_{c}+\frac{\text{1}}{2}\nu_{c}^{2} \right)$ and standard deviation of *t* is given by τ_c_μ_c_ where $\tau_{c}=\sqrt{exp\left( \nu_{c}^{2} -1 \right)}$. This yields:

$$\xi_{c}=\text{log}\left( \mu_{c} \right)-\frac{\text{1}}{2}\nu_{c}^{2}$$

$$\nu_{c}=\sqrt{\text{log}\left( \tau_{c}^{2}+1 \right)}$$

CRC-AIM samples a random ST as:

$${lognormal}_{rand}={exp}^{\xi_{c}+\nu_{c}z_{rand}}$$

where z_rand_ is a random Z value generated from the inverse of the standard normal CDF.

*CRC stage upon detection*

Similar to CRC-SPIN v1.0, CRC-AIM derived a multinomial logistic regression model to determine AJCC stage based on CRC size. This approach produces a different stage distribution compared to SimCRC and MISCAN (and compared to the empirical SEER 1975-1979 data) (**Figure 6**). SimCRC and MISCAN use a different approach, although their approach is not described in sufficient detail to replicate.^1^ CRC-SPIN v2.x appears to correct for this discrepancy in stage distribution,^11^ although the updated approach is also not well-described.

*CRC cause-specific survival*

Many of CRC-SPIN v1.0’s published natural history modeling outputs were generated prior to a 2013 CRC survival update.^12^ CRC-SPIN v1.0’s initial CRC survival model was based on relative survival using SEER survival data from 1975 to 1979 (prior to the diffusion of colorectal cancer screening).^3^ This data was imported into CAN*SURV to generate proportional hazards models stratified by location (colon and rectum) and American Joint Committee on Cancer (AJCC) CRC stage (stages I through IV), with sex and age as covariates.^10^

In 2013, CISNET updated the CRC survival methodology,^10,12^ and since that time, all three models (CRC-SPIN, SimCRC, and MISCAN) use the same survival method.^2^ The updated survival methodology is additive-hazards-based with time-varying covariables and takes into account improved survival for more recently diagnosed CRC relative to those diagnosed in 1975 due to improvements in therapy and surveillance methods. This method is used to determine CRC survival based on sex, stage at diagnosis, age at diagnosis, and year of diagnosis, with the optional inclusion of race for those models that can incorporate race as a risk factor (ie, SimCRC). This consistency reduced the between-model variability related to CRC survival, allowing investigators to directly compare differences in model results primarily due to natural history structure and assumptions, although it introduces the possibility of systemic bias if inherent issues exist within the survival approach. Ultimately, although the CISNET teams provide a general theoretical discussion into their methodology, an enumeration of the parameters themselves and a detailed description of how they are implemented in their models is not published.^12^

CRC-AIM implements cause-specific survival as a set of parametric regression equations that model survival probabilities, stratified by location and AJCC CRC stage, as a function of sex and age at diagnosis. To compare the survival outcomes of CRC-AIM to those of the CISNET models, we generated survival curves that mimicked the timeframes of SEER data used by CISNET both before and after their survival update in 2013. We generated parametric cause-specific survival curves using SEER data from 1975 to 1979 and compared natural history outcomes to publicly available values across multiple CISNET publications before the survival update. Additionally, we generated a similar set of curves using 2000-2003 SEER data and compared natural history outcomes to results described in the 2016 USPSTF modeling report, after the survival update. Both sets of comparisons are described in greater detail in the main manuscript.

Here, we include detailed descriptions of the SEER queries that generated data for the parametric linear regression models and comprehensive information about those models that were used to create the cause-specific survival curves (**Table S1**). Specifically, for each condition—using the SEER data timeframes before and after the survival update, subdivided by AJCC CRC stage and location—we include the survival curves themselves along with model selection details and fitted model diagnostics and parameter estimates. Here is a complete list of the figures and tables that correspond to each condition:

- 1975-1979 SEER data:
  - Stage I: colon (**Figure S6**, **Table S10**); rectum (**Figure S7**, **Table S11**)
  - Stage II: colon (**Figure S8**, **Table S12**); rectum (**Figure S9**, **Table S13**)
  - Stage III: colon (**Figure S10**, **Tables S14-15**); rectum (**Figure S11**, **Table S16**)
  - Stage IV: colon (**Figure S12**, **Table S17**); rectum (**Figure S13**, **Tables S18-19**)
- 2000-2003 SEER data:
  - Stage I: colon (**Figure S14**, **Table S20**); rectum (**Figure S15**, **Tables S21-22**)
  - Stage II: colon (**Figure S16**, **Table S23**); rectum (**Figure S17**, **Tables S24-25**)
  - Stage III: colon (**Figure S18**, **Tables S26-27**); rectum (**Figure S19**, **Tables S28-29**)
  - Stage IV: colon (**Figure S20**, **Tables S30-31**); rectum (**Figure S21**, **Tables S32-33**)

SEER queries

We queried the SEER database to extract data from the 1975-1979 and 2000-2003 timeframes using the criteria described below. The case listing file was saved as a comma-separated values (CSV) file and imported into JMP v13.0 (SAS Institute) for statistical analysis. Furthermore, we describe how the SEER data was modified in JMP prior to fitting the regression models (**Table S9**).

*SEER query for 1975-1979 survival*

Software

Surveillance Research Program, National Cancer Institute SEER*Stat software (www.seer.cancer.gov/seerstat) version 8.3.5. 10/05/2018

Data

Surveillance, Epidemiology, and End Results (SEER) Program ([www.seer.cancer.gov](http://www.seer.cancer.gov)) SEER*Stat Database: Incidence – SEER 9 Regs Research Data, Nov 2017 Sub (1973-2015) <Katrina/Rita Population Adjustment> - Linked To County Attributes – Total U.S., 1969-2016 Counties, National Cancer Institute, DCCPS, Surveillance Research Program, released April 2018, based on the November 2017 submission.

Statistic

Cause-Specific Survival

Definition of Cause of Death: Dead due to cancer using SEER cause-specific death classification

Missing/Unknown COD: Exclude From Analysis

Selection

Select Only: Malignant Behavior, Known Age

Exclude: All Death Certificate Only and Autopsy Only, Alive with No Survival Time

Exclusions to Match the Expected Survival Table: Age Values Not Found in Table, Invalid Year, Values Not Found for Other Variables in Table

Case Selection

{Site and Morphology.Site recode ICD-O-3/WHO 2008} = '	Colon and Rectum'

AND {Site and Morphology.Histologic Type ICD-O-3} = 8000-8001,8010,8020,8140,8210-8211,8220-8221,8260-8263,8480-8482,8490

AND {Race, Sex, Year Dx, Registry, County.Year of diagnosis} = '1975','1976','1977','1978','1979'

Multiple Primary Selection: First Primary Only (Sequence Number 0 or 1)

Parameters

Pre-calculated Duration: Survival Months (from complete dates)

Study Cutoff: Dec 2015

Censor When Attained Age Exceeds Expected Table Max

Display: Case Listing

Intervals: Number: 999, Months Per: 1

Table

Case Listing:

- SEER historic stage A
- 2-Digit NS EOD part 1 (1973-1982)
- AJCC 5^th^ Ed Schrag Code 1975-1979
- Age recode with <1 year olds
- Survival months
- Survival months flag
- Vital status recode (study cutoff used)
- Sex
- COD to site recode
- COD to site rec KM
- SEER cause-specific death classification
- SEER summary stage 1977 (1995-2000)
- SEER other cause of death classification
- Site Colon vs Rectum

*SEER query for 2000-2003 survival*

Software

Surveillance Research Program, National Cancer Institute SEER*Stat software (www.seer.cancer.gov/seerstat) version 8.3.5. 10/05/2018

Data

Surveillance, Epidemiology, and End Results (SEER) Program ([www.seer.cancer.gov](http://www.seer.cancer.gov)) SEER*Stat Database: Incidence – SEER 9 Regs Research Data, Nov 2017 Sub (1973-2015) <Katrina/Rita Population Adjustment> - Linked To County Attributes – Total U.S., 1969-2016 Counties, National Cancer Institute, DCCPS, Surveillance Research Program, released April 2018, based on the November 2017 submission.

Statistic

Cause-Specific Survival

Definition of Cause of Death: Dead due to cancer using SEER cause-specific death classification

Missing/Unknown COD: Exclude From Analysis

Selection

Select Only: Malignant Behavior, Known Age

Exclude: All Death Certificate Only and Autopsy Only, Alive with No Survival Time

Exclusions to Match the Expected Survival Table: Age Values Not Found in Table, Invalid Year, Values Not Found for Other Variables in Table

Case Selection

{Site and Morphology.Site recode ICD-O-3/WHO 2008} = '	Colon and Rectum'

AND {Site and Morphology.Histologic Type ICD-O-3} = 8000-8001,8010,8020,8140,8210-8211,8220-8221,8260-8263,8480-8482,8490

AND {Race, Sex, Year Dx, Registry, County.Year of diagnosis} = '2000','2001','2002','2003'

Multiple Primary Selection: First Primary Only (Sequence Number 0 or 1)

Parameters

Pre-calculated Duration: Survival Months (from complete dates)

Study Cutoff: Dec 2015

Censor When Attained Age Exceeds Expected Table Max

Display: Case Listing

Intervals: Number: 999, Months Per: 1

Table

Case Listing:

- AJCC 5^th^ Ed Schrag Code 1988-2003
- Age recode with <1 year olds
- Survival months
- Survival months flag
- Vital status recode (study cutoff used)
- Sex
- COD to site recode
- COD to site rec KM
- SEER cause-specific death classification
- SEER summary stage 1977 (1995-2000)
- SEER summary stage 2000 (2001-2003)
- SEER other cause of death classification
- Site Colon vs Rectum

AJCC staging for 1975-1979 SEER data

SEER registry data prior to 1988 uses historic staging criteria that categorize cancer as local, regional, and distant. However, the AJCC staging system uses tumor, lymph node, and metastasis information to stage cancer. Since the CISNET CRC models use AJCC stages I through IV, we needed to convert pre-1988 SEER registry data. We used code developed by Deborah Schrag, coauthor on the CISNET 2013 survival update,^12^ which essentially is a complicated user-defined variable that uses 12 standard SEER variables to recode stage according to the AJCC 5^th^ Edition AJCC Cancer Staging Manual.^13^ Although the AJCC staging manual staging is currently in its eighth edition,^14^ the changes since the fifth edition primarily involve subgrouping stages II through IV and providing prognosis-related details.^15^

Functional implementation of survival models in CRC-AIM

There are some technical issues related to survival assumptions that must be considered when implementing a screening overlay. To illustrate, consider an example of an individual with two synchronous preclinical CRCs, one stage I and the other stage III. According to the documentation of CRC-SPIN’s natural history model, the first cancer to become clinically detectable—whichever cancer’s sojourn time expires first—determines CRC-related survival.^10^ If the stage I cancer’s sojourn time expires before the stage III cancer, the stage I cancer would determine survival. However, if a cancer is detected through screening, CRC-SPIN assumes the detection of all synchronous cancers and that all existing cancer is treated.^10^ In this example, both the stage I and stage III cancers would be observed by an endoscopist. The CRC-SPIN authors do not explicitly state which cancer dictates the choice of survival function for this scenario, or when the survival function would be applied (ie, after the ST expiration for the stage I cancer or for the stage III cancer). If the stage III cancer dictates survival, the screened individual may likely have a shorter lifespan compared to natural history, which was dictated by the stage I survival function.

This issue also extends to the removal of adenomas through screening. Consider a scenario in which an individual has a large, fast-growing adenoma A1 and a preclinical cancer pCRC2, which arose from another adenoma A2. In the natural history model, assume that pCRC2 transitions to clinical detection CRC2 through expiration of sojourn time into a stage I CRC, after which the survival method would be applied. In a hypothetical screening scenario, the adenoma that led to the CRC (A2) may have been removed but A1 could have been missed. This adenoma could hypothetically transition to preclinical CRC (pCRC1) and then a stage IV clinically detected CRC (CRC1) within a year after the appearance of the non-existent stage I CRC (CRC2) in the “parallel universe” natural history model. This individual would likely die sooner in the screening scenario from the stage IV CRC than the parallel universe individual that had stage I CRC in the natural history arm.

Although these corner-case scenarios can be addressed in numerous ways, the specific approaches undertaken by CRC-SPIN is not publicly described.

In CRC-AIM, if a cancer is detected by screening, the survival method is determined by the CRC stage diagnosed upon detection. This approach potentially confers a survival benefit if an earlier-stage CRC is detected relative to the CRC stage present when the sojourn time expires in a parallel universe. To prevent bias resulting from automatically assigning a worse survival outcome for screen-detected CRC compared to natural history, the survival function will be applied only when the sojourn time would have otherwise expired for the screen-detected cancer. In other words, the CRC survival functions in CRC-AIM will not be implemented during the lead-time of the cancer. For CRC-AIM, upon clinical detection, the cancer stage and location are used to define the survival model. Based on the survival model, age at diagnosis and sex may be used to determine the cumulative probability of survival for each yearly interval after diagnosis. We restricted the number of intervals to 100 (the maximum age of an individual minus the minimum age one can be diagnosed with CRC, or 120 minus 20). A random uniform (0,1) distribution is used as an inverse CDF lookup to determine CRC-based survival years. If the random uniform distribution exceeds the maximum probability, then the maximum survival years are applied.

Linear regression modeling

We fit five separate parametric linear regression models for each SEER dataset (1975-1979 and 2000-2003), AJCC stage (stages I to IV), and location (colon versus rectum). These models were based on five different distributions to describe survival time—Weibull, lognormal, exponential, Fréchet, and loglogistic. Model effects were sex and age at CRC diagnosis and statistical significance of an effect was based on the Wald test. Age at diagnosis was subdivided into 20-49, 50-59, 60-69, 70-79, and ≥80 years, consistent with the categories used by CISNET.^12^ Right-censoring was performed as appropriate. Any subject with a time of death that was reliably recorded as 0 months was recoded to 0.5 months to prevent model-fitting issues. Model selection was based on smallest Akaike information criterion (AICc) value for the fitted distribution across the five separate models for a given combination of AJCC stage and location (**Table S1**).

CRC-AIM uses the CDF to describe the cumulative transition probability of CRC-specific death as a function of sex and age at CRC diagnosis. Regression-based coefficients are multiplied by an indicator function if sex or age at diagnosis criteria are met (sex indicator is 1 if met, -1 if not met; age at diagnosis indicator is 1 if met, 0 if not met) and are linearly combined with an intercept, represented by λ in Weibull models and µ in loglogistic, lognormal, and Fréchet models..

For 1975-1979 and 2000-2003 Stage I colon cancers, the Weibull model was selected. The probability of transition to CRC-specific death at or before time *t* is given by:

$\text{F(}\text{t; k}\text{, }\text{λ}\text{) = 1 }-\text{ }e^{{-(\frac{t}{\lambda})}^{k}}$ for t ≥ 0.

with *k* as the shape parameter, and λ as the scale parameter. Since *k* < 1 in each of the two Weibull models, the failure rate decreases over time, representing the curative effect one would expect in a proportion of diagnosed and treated early-stage cancer.

For 1975-1979 Stage IV rectal-based cancers and 2000-2003 Stage I rectal-based cancers, the loglogistic model was selected:

$\text{F(t; }\text{μ}\text{, }\text{σ}\text{) = }\Phi_{\text{logis}}\left[ \frac{\text{log}_{\text{(t)}}-\mu}{\sigma} \right]$ for t > 0

where

$$\Phi_{\text{logis}}\text{(z) = }\frac{\text{1}}{\text{1 + }e^{-z}}$$

For 1975-1979 Stage I, II, and III rectal-based cancers; Stage II colon-based cancers; and 2000-2003 Stage II, III, and IV colon- and rectal-based cancers, the lognormal model was selected:

$\text{F(t; μ, σ) = }\Phi_{\text{nor}}\left[ \frac{\text{log}_{\text{(t)}}-\mu}{\sigma} \right]$ for t > 0

where

$$\Phi_{\text{nor}}\text{(z) = }\int_{-\infty}^{z} \Phi_{\text{nor}}\left( w \right)dw$$

For 1975-1979 Stage III and IV colon cancer, the Fréchet model was selected:

$\text{F(t; μ, σ) = }exp\left[ -exp\left( -\frac{{log}_{(t)}-\mu}{\sigma} \right) \right]$for t > 0

**Additional CRC Incidence Comparison**

We want to report additional information not explicitly described in Kuntz et al^6^ to improve transparency and completeness in terms of numerical comparisons. The 20-year CRC incidence for individuals with no underlying lesions is not reported,^6^ although CRC-AIM overlaps with SimCRC and CRC-SPIN across the 20-year (CRC-AIM: 0.36%) and 30-year (CRC-AIM: 1.9%) follow-up period (**Figure 8**). In addition, the exact values for 30-year cumulative CRC incidence for the subgroup with underlying lesions is not reported^6^—CRC-AIM yielded 25.6% (**Figure 8**).

**Multinomial Logistic Regression for CRC Stage based on Size**

CRC-SPIN v1.0 uses a multinomial logistic regression model to predict CRC stage, based on the American Joint Committee on Cancer (AJCC) guidelines, that are conditional on CRC size.^10^ The model was developed using SEER data for colorectal cancer cases diagnosed between 1975-1979. However, the parameterization of CRC-SPIN v1.0 is not described in the literature.

Although the authors of the CRC-SPIN CISNET profile admit that the dependency between CRC stage and size at clinical detection is weak,^10^ linking both variables enables a direct method to confer a greater survival benefit for early (ie, preclinical) CRC detection based on size. Upon initiation of a preclinical cancer, CRC-SPIN v1.0 assigns an initial size (0.5 mm) and a size upon transitioning to clinically detectable cancer (ie, the cancer size when ST expires). CRC-SPIN uses an exponential growth function to define CRC size at any time during ST. If CRC is detected during ST (ie, while it is preclinical), it will be smaller and could be assigned a less advanced AJCC stage compared to the same CRC upon clinical detection based on the logistic regression formula.

One consequence of this method is that it generates a different distribution of AJCC staging in a natural history modeling scenario compared to the other CISNET models (MISCAN and SimCRC) and the empirically observed distribution from SEER 1975-1979. In the absence of screening, CRC-SPIN generates fewer stage IV cancers and more stage II and stage III cancers^1^ (**Figure 6**). In contrast, CRC-SPIN v2.x first simulates CRC stage at clinical detection, then the size at clinical detection, which is stratified by stage.^11^ The authors explain that this inverted approach allows for greater flexibility in specifying stage distribution, although their exact methodology is not clearly characterized.^11^

Here, we describe in detail our approach to develop a multinomial logistic regression model for CRC-AIM to predict AJCC stage conditional on CRC size.

CRC-AIM’s multinomial logistic regression formula is based on 1975-1979 SEER data, which uses the code developed by Deborah Schrag to convert pre-1988 SEER registry data to AJCC staging categories^13^ (see “AJCC staging for 1975-1979 SEER data” for more information.) The SEER query resulted in approximately 33,485 CRC cases with reported sizes from 1 mm to 97 mm (inclusive) and about 1,453 results (~4% of the data) coded as “≥98 mm,” which we recoded to 98 mm (**Figure S22A**). We noted a monotonically decreasing proportion of stage I CRCs and a monotonically increasing proportion of stage IV CRCs conditioned on size (up to 85 mm). There are low counts for smaller CRC sizes—only 113 (0.34%) CRCs in the 1-5 mm inclusive size category, and only 42 (0.13%) CRCs in the 1-3 mm inclusive size category—which indicates uncertainty in the stage distribution of smaller CRCs. The implication of the empirical size distribution for these smaller lesions is that even if a 1 mm CRC were detected (roughly the size of CRC initiation in the model), the cancer would only have a 60% chance of being stage I. (Notably, the discrepancy in small polyp sizes could reflect changes in endoscopic technology^16^ or variability in polyp size estimation,^17^ although these hypotheses are beyond the scope of this analysis.)

To better understand the staging of small lesions, we referred to two other date ranges within the SEER dataset: 1988-1992 and 2011-2015 (**Figure S22B-C**). Both ranges indicate that very small CRC lesions (1 mm to 5 mm) are associated with ~90% chance of being designated as stage I. We specified the following *a priori* conditions for the multinomial logistic regression model:

- allowing for 90% stage I CRC for very small lesions (1 mm)
- conforming to a monotonic function for stage I CRC, decreasing probability as size increases
- conforming to a monotonic function (≤85 mm) for stage IV CRC, increasing probability as size increases
- accurately extrapolating to CRCs ≥98 mm, since cancers are modeled up to 140 mm in size
- basing the model on the 1975-1979 SEER dataset

We parameterized the regression by fitting numerous multinomial logistic regression models to the SEER 1975-1979 data and considered various size transformations, including using a fractional polynomial approach. In all models, we treated size as a continuous factor and recoded a size of “≥98 mm” as 98 mm.

The final model that satisfies our pre-specified conditions uses fractional polynomials (-0.5, 1) for size, and the model is summarized in **Table S34**. Wald tests indicate both size terms in the model are statistically significant. Model RSquare is poor and the lack of fit is statistically significant. Area under the curve (AUC) is very poor for stage II (0.56) and stage III CRC (0.52), and only modestly informative for stage I (0.69) and stage IV (0.57) CRCs (**Figure S24**). Very similar outputs were obtained across other models, however, and as mentioned previously, the weak association between CRC size and stage was noted by the CRC-SPIN authors.^10^

The final multinomial logistic regression function is plotted as an overlay with the 1975-1979 SEER empirical data (**Figure S23**), with a vertical red line indicating the model-predicted stage distribution for CRC-AIM’s extrapolated CRC sizes (ie, 98 mm to 140 mm). The implementation is described in further detail in the **Methods** section of the manuscript.

**Size distribution of clinically detected CRC**

When a preclinical cancer initiates, the size of the cancer upon clinical detection (in the absence of screening—ie, the expiration of sojourn time) is determined. CRC-SPIN’s size at clinical detection is based on the overall SEER distribution of CRC size from 1975-1979,^10^ but the parameterization of this size is not explained. Here, we briefly describe the steps we took to derive this distribution for CRC-AIM.

We conducted a SEER query of the 1975-1979 registry data using the conditions described below:

*SEER query for 1975-1979 CRC size*

Software

Surveillance Research Program, National Cancer Institute SEER*Stat software ([www.seer.cancer.gov/seerstat](http://www.seer.cancer.gov/seerstat)) version 8.3.5. accessed 07/12/2018

Data

Surveillance, Epidemiology, and End Results (SEER) Program ([www.seer.cancer.gov](http://www.seer.cancer.gov)) SEER*Stat Database: Incidence – SEER 18 Regs Research Data + Hurricane Katrina Impacted Louisiana Cases, Nov 2017 Sub (1973-2015 varying) – Linked to County Attributes – Total U.S., 1969-2016 Counties, National Cancer Institute, DCCPS, Surveillance Research Program, released April 2018, based on the November 2017 submission.

Selection

Select Only: Malignant Behavior, Known Age, Cases in Research Database

{Site and Morphology Site recode ICD-O-3/WHO 2008}=’Colon and Rectum’

AND {Site and Morphology Histologic Type ICD-O-3}=8000-8001,8010,8020,8140,8210-8211,8220-8221,8260-8263,8480-8482,8490

AND {Race, Sex, Year Dx, Registry, County.Year of diagnosis}='1975','1976','1977','1978','1979'

Table

Expanded EOD(1) - CP53 (1973-1982)

Expanded EOD(2) - CP54 (1973-1982)

Expanded EOD(3) - CP55 (1973-1982)

Expanded EOD(4) - CP56 (1973-1982)

Expanded EOD(5) - CP57 (1973-1982)

Expanded EOD(6) - CP58 (1973-1982)

Expanded EOD(7) - CP59 (1973-1982)

Expanded EOD(8) – CP60 (1973-1982)

Expanded EOD(9) – CP61 (1973-1982)

Expanded EOD(10) – CP62 (1973-1982)

Expanded EOD(11) – CP63 (1973-1982)

Expanded EOD(12) – CP64 (1973-1982)

Expanded EOD(13) – CP65 (1973-1982)

SEER historic stage A

2-Digit NS EOD part 1 (1973-1982)

AJCC 5^th^ Ed Schrag Code 1975-1979

A total of 50,743 CRCs were queried. The SEER Extent of Disease (EOD) coding scheme records CRC sizing information in the unit of millimeter: a value of 0 to 9 is recorded in EOD(1) – CP53 (1973-1982) for the value in the tens place, and a value of 0 to 9 is recorded in EOD(2) – CP54 (1973-1982) for the value in the ones place. Although this theoretically allows for CRC sizes up to 99 mm, this is not how the information is represented. Instead, size is actually coded up to 97 mm, with tumors that are greater than or equal to 98 mm coded as “98”.

Additionally, the following special codes are used^18^:

- 00: No mass
- 0&: Microscopic focus or foci only
- -- Not stated

The following criteria were used to filter out results from further analysis: Unstaged CRC (7,692 records) and CRC size where size was recorded as --, 00, 0&, Blank(s)Blank(s) (14,912 records). Ultimately, a total of 17,258 results were excluded, resulting in 33,485 results (50,743 minus 17,258). Of those 33,485 results, 32,032 CRCs range from 1 mm to 97 mm and 1,453 CRCs are recorded as 98 mm, corresponding to the “≥98 mm” category (**Figure S25**). We treated the 1,453 “≥98 mm” records as missing observations and modeled these values, extrapolating a non-truncated right-tail of the distribution. Specifically, we parametrically modeled the CRC counts from 50 mm to 97 mm and extrapolated the counts modeling past 97 mm until the extrapolated total equaled ~1,453 observations (actual n = 1,484). The extrapolated counts are combined with the original counts and the entire distribution was fit to obtain the probability density function of CRC size.

We plotted the counts of discrete CRC size categories from the SEER registry data and observed that most CRCs in this subsample were rounded to the nearest centimeter (eg, 50 mm, 60 mm, 70 mm, etc.) (**Figure S26**). Another set of CRCs was rounded to the nearest half-centimeter (eg, 55 mm, 65 mm, 75 mm, etc.). Finally, a third set was rounded to the nearest millimeter. Notably, counts rounded to the nearest centimeter are biased because they are inclusive of CRCs rounded to the nearest half-centimeter (eg, a 52 mm CRC rounds to 50 mm) and those rounded to the nearest 1 mm (eg, a 50.4 mm CRC rounds to 50mm). Similarly, the counts on the half-centimeter (eg, 55 mm, 65 mm, etc.) are biased since they are inclusive of counts rounded to the nearest millimeter. These biases were ignored for this simple modeling exercise.

To perform the model extrapolation, we created three separate Poisson regression models for each rounding scenario (**Figure S27**). The Poisson regressions were applied to each missing size group associated with each rounding scenario past 97 mm. For example, the regression for the nearest centimeter rounding scenarios was applied to sizes of 100 mm, 110 mm, etc. Size was increased by millimeter increments until ~1,453 observations were obtained (**Table S35**). Finally, the 32,032 values coded from 1 mm to 97 mm were combined with the extrapolated 1,484 values from 98 mm to 140 mm (**Figure S28**). The probability density function (PDF) of the generalized log distribution is sampled to generate a CRC size at clinical diagnosis from 1 mm to 140 mm.

**Model Cross-Validation**

To compare the overall outcomes of CRC-AIM to those of the CISNET models, we conducted three validation analyses: (1) quantitative analysis; (2) qualitative analysis; and (3) medical decision making analysis. We performed analyses comparing the colonoscopy-related outcomes and the stool-based testing outcomes (FIT, HSgFOBT, and mt-sDNA) both by comparing the outputs from CRC-AIM against outputs from each of the CISNET models, and by similarly comparing each CISNET model against one another.

Although we are using three CISNET models—SimCRC, MISCAN, and CRC-SPIN—as primary comparators to validate the outcomes of CRC-AIM, none of these models are considered a “reference standard”. Therefore, we conducted comparator analyses in which one of the CISNET models functioned as a “comparator” and the model being compared as a “candidate method”.

Quantitative analyses

Common regression techniques such as ordinary least-squares would not be relevant for this analysis because there is measurement variability in X and Y variables. Recommended approaches include orthogonal (e.g. Deming regression) or non-parametric (e.g. Passing-Bablok) regression. We selected Passing-Bablok for our comparison.

For each metric reported in the CISNET supplemental tables related to screening outcomes—colonoscopy (COL), Surveillance COL, Symptom COL, Total COL, Complications, CRC Cases, CRC Deaths, life-year (LY) with CRC, life-year gain (LYG), CRC Incidence Reduction, CRC Mortality Reduction— we performed regressions of the candidate method (Y-axis) versus the comparator method. (For stool-based tests, we added comparisons for Follow-Up COL and number of stool tests)

For these quantitative regressions, the intercept is referred to as "systematic bias" and the slope is referred to as "proportional bias." If the slope is 1 and intercept is 0, there is no measurable bias. In other words, if the 95% confidence interval (CI) of the intercept contains 0, and the 95% CI of the slope contains 1, the hypothesis that no difference (ie, the proportional and systemic bias) exists between the candidate and comparator methods cannot be rejected.  If systematic and/or proportional bias exists, the biases between the models should not be "clinically meaningful."  We identify any instance in which CRC-AIM shows statistically relevant bias against a comparator model but we concluded that the difference is not meaningful (from a modeling-difference standpoint) if the absolute difference between CRC-AIM (candidate) and a given CISNET comparator is within the absolute difference 95% CI of any CISNET model (candidate) versus a given CISNET comparator.

We perform this bias evaluation for any screening outcomes metric slightly below the lowest observed value across all comparisons (e.g. LOW in Column B, worksheet “SummaryQuantCOL” in Supplemental Excel Files) and perform the bias evaluation slightly above the highest observed value across all comparisons (HIGH in column B, worksheet “SummaryQuantCOL” in Supplemental Excel Files), in order to capture bias across the full measurement range of outcomes. The final evaluation for quantitative comparisons is in Column P.

*Qualitative analyses*

We performed an efficient frontier analysis with colonoscopy and stool-based tests. Using the efficient and near-efficient strategies from CRC-AIM and those from the CISNET technical report (Zauber et al), we performed standard qualitative comparisons for strategies that are on and off the efficient frontier. CISNET considered near-efficient strategies as being on the efficient frontier. However, since "near-efficient" strategies are basically "equivocal" according to clinical best practice, we perform two qualitative comparisons, one in which near-efficient strategies are considered on the frontier and another in which near-efficient strategies are interpreted as "OFF" the frontier. For each comparison, we demonstrate Negative, Positive, and Overall Percent Agreement (NPA, PPA, OPA, respectively) (see **Supplemental Excel Files**).

*Medical decision making analysis*

Using the efficient frontier results from CRC-AIM (SUPP TABLES) and the CISNET models (Knudsen et al), we performed a set of medical decision making analyses using CRC-AIM and compared the results to the exact same decisions made using the CISNET models. Decisions made for screening start/stop age, benchmark colonoscopy, and optimal strategy were similar for CRC-AIM compared to the CISNET models (see **Supplemental Excel File**).

**Figure S1. Percent of adenomas that had developed within 10 or 20 years of clinical colorectal cancer diagnosis for SimCRC, CRC-SPIN, and CRC-AIM.** Adenomas within (A) 10 years and (B) 20 years are displayed by age. Data adapted from Kuntz et al.^6^

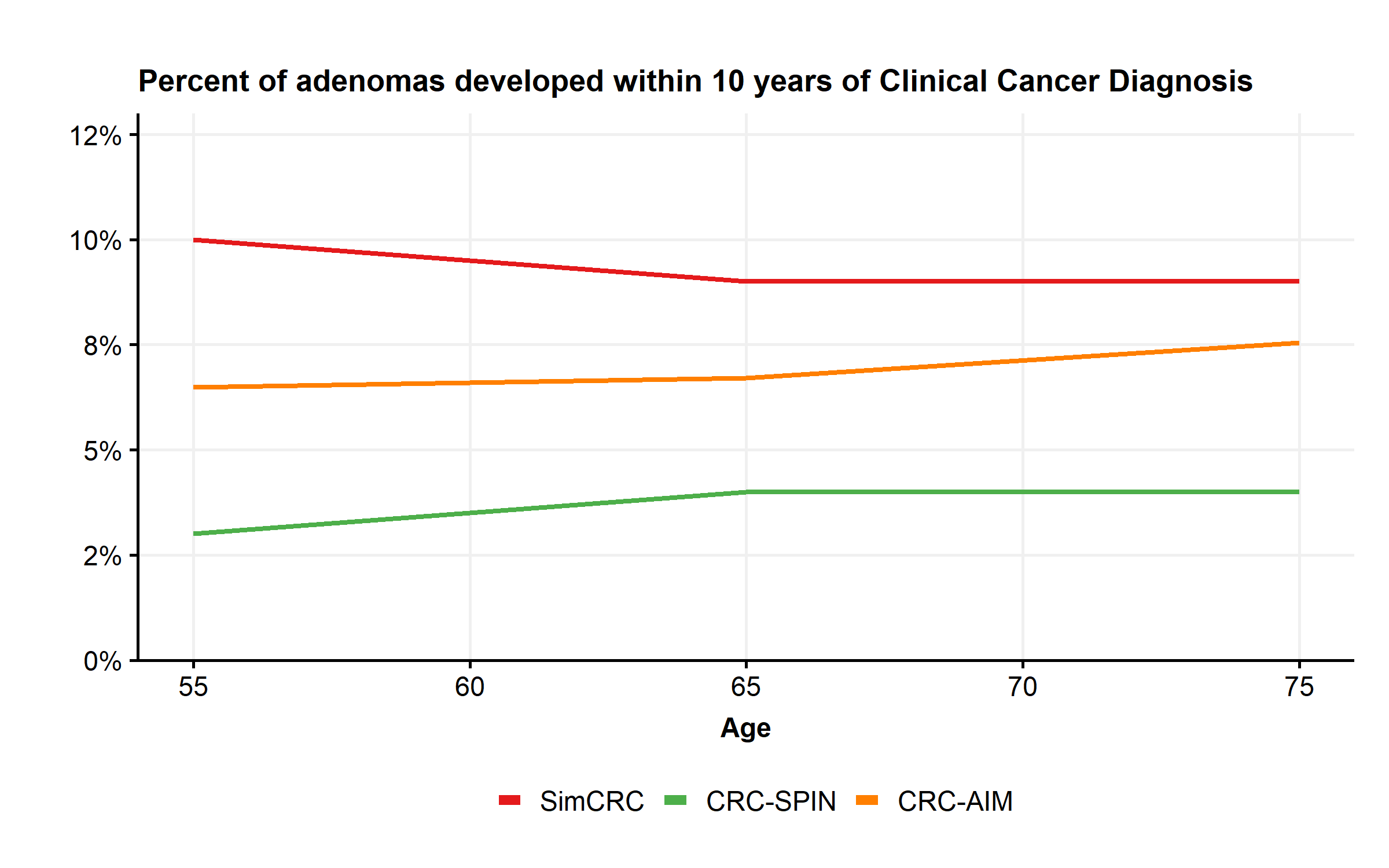

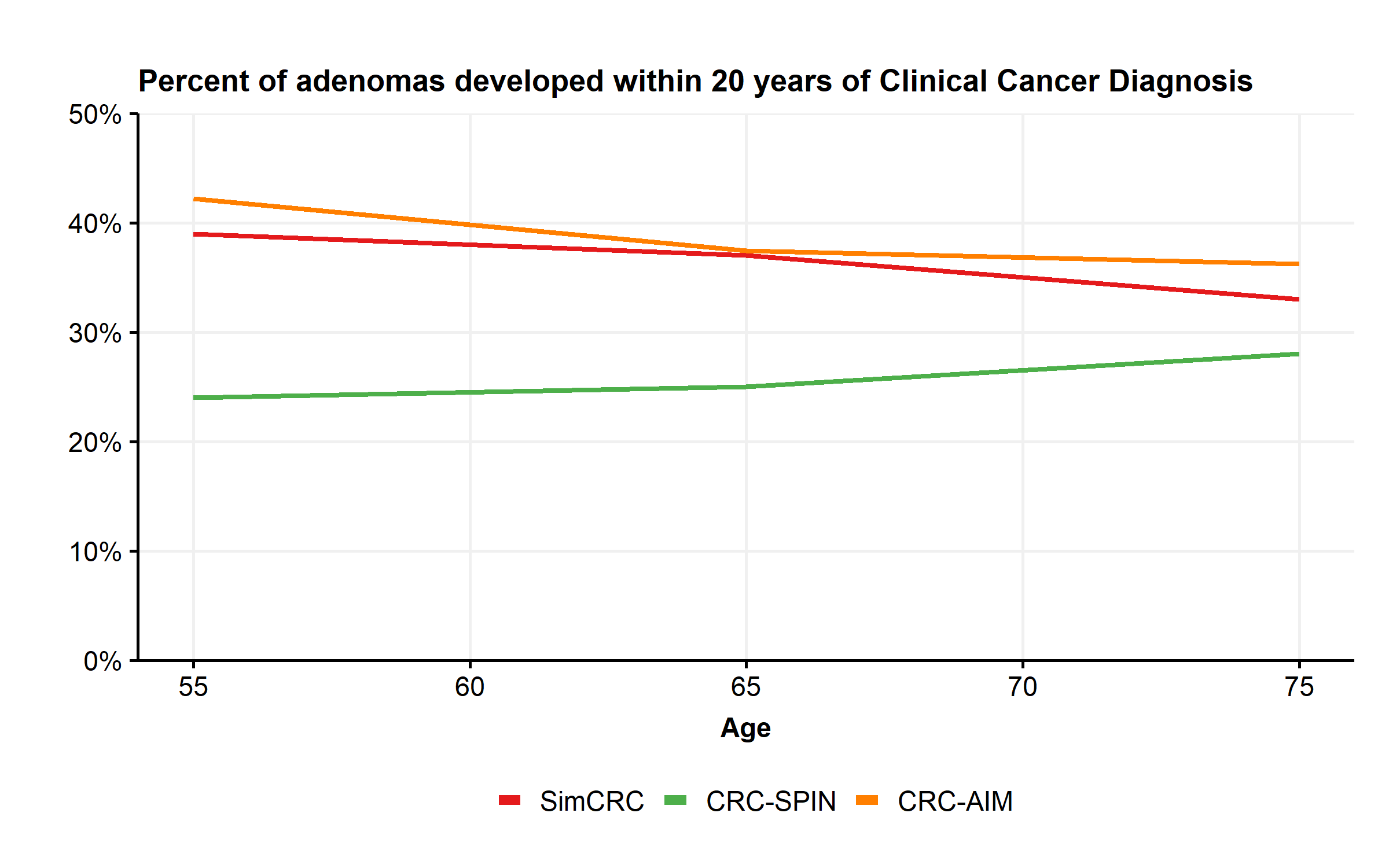

**Figure S2. Adenoma distribution by location among individuals aged 40 and older by model.** Data adapted from Zauber et al.^1^

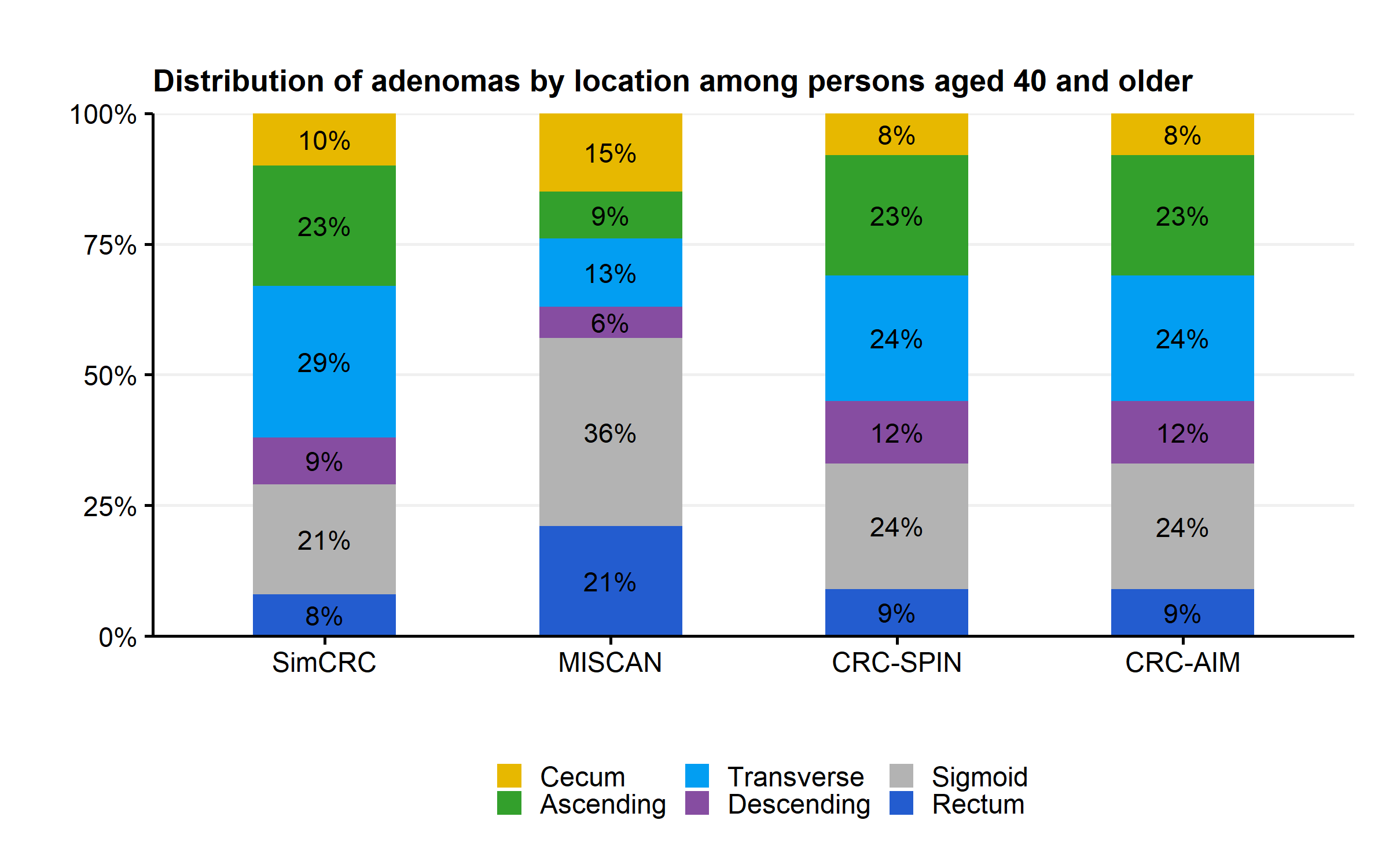

**Figure S3. Comparison of the distribution of the most-advanced adenoma size by age between CISNET models (SimCRC, MISCAN and CRC-SPIN) and a prototype version of CRC-AIM.** Size distribution is evaluated into 40, 60, and 80 years. The CRC-AIM prototype was prior to recalibration. Data adapted from Zauber et al.^1^

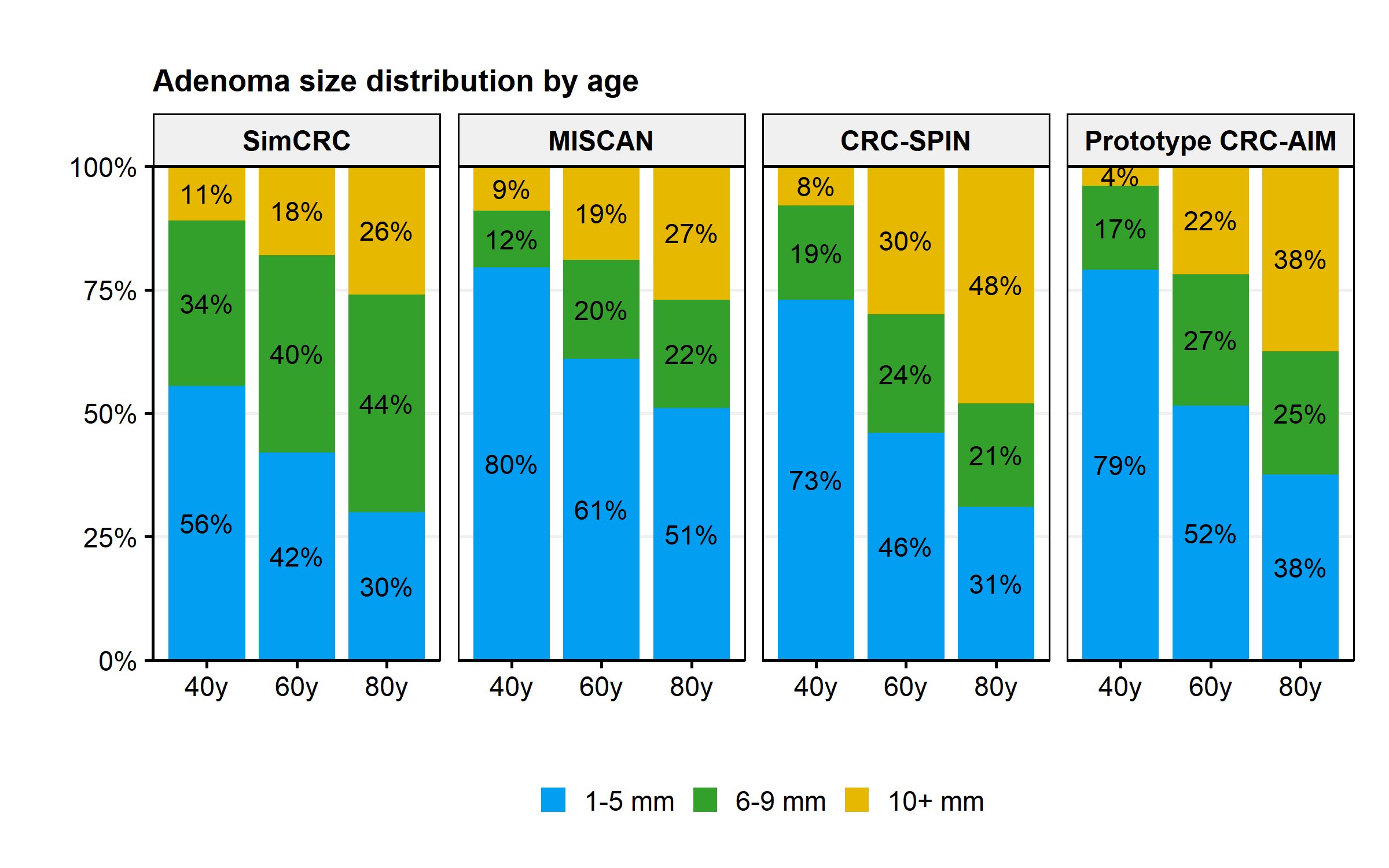

**Figure S4. Comparison of colorectal cancer (CRC) incidence and cumulative CRC risk between CISNET models (SimCRC, MISCAN and CRC-SPIN) and a prototype version of CRC-AIM.** (A) CRC incidence by age and model and (B) cumulative probability of developing CRC. The CRC-AIM prototype was prior to recalibration. Data adapted from Zauber et al.^1^

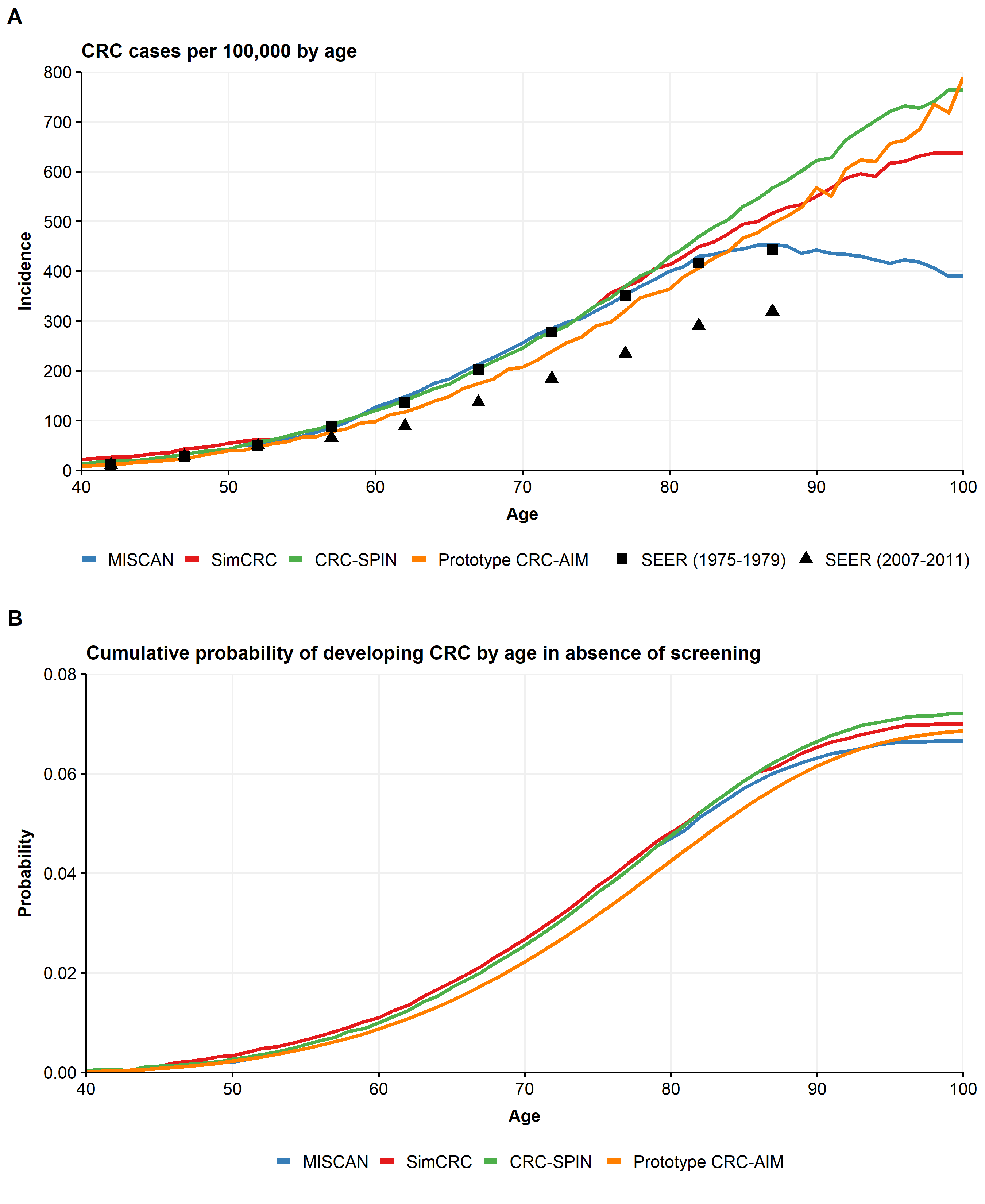

**Figure S5. Cumulative transition probability for an adenoma to become preclinical cancer as a function of adenoma size.** The transition probability is visualized at three different ages at adenoma initiation (25, 45, and 65 years), stratified by sex and adenoma location. The vertical red line indicates the maximum adenoma size in the model (50 mm).

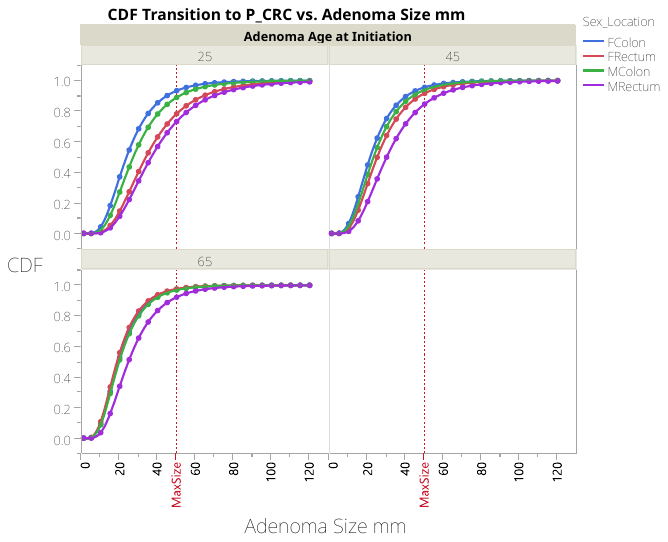

**Figure S6. Modeled cause-specific survival for 1975-79 stage I colorectal cancer (CRC) in the colon, stratified by age at diagnosis and sex.** CRC-specific survival is represented as percent survival by years since diagnosis. Note: The abnormal residual behavior at low probability is due to recoding survival month from 0 months to 0.5 months—the abnormal plot observations have survival month values of 0.5—and has little practical impact on survival, which is based on yearly increments.

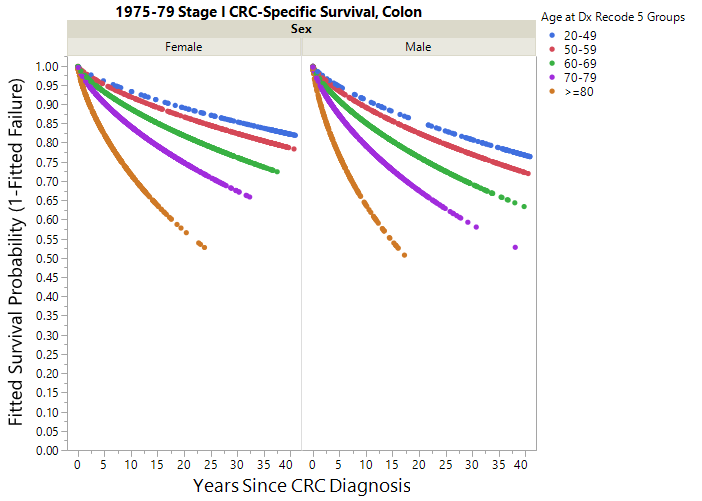

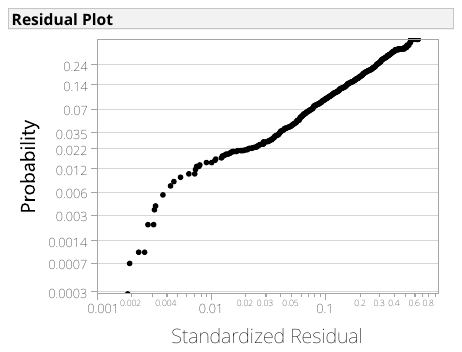
**Figure S7. Modeled cause-specific survival for 1975-79 stage I colorectal cancer (CRC) in the rectum, stratified by age at diagnosis and sex.** CRC-specific survival is represented as percent survival by years since diagnosis. Note: The abnormal residual behavior at low probability is due to recoding survival month from 0 months to 0.5 months—the abnormal plot observations have survival month values of 0.5—and has little practical impact on survival, which is based on yearly increments.

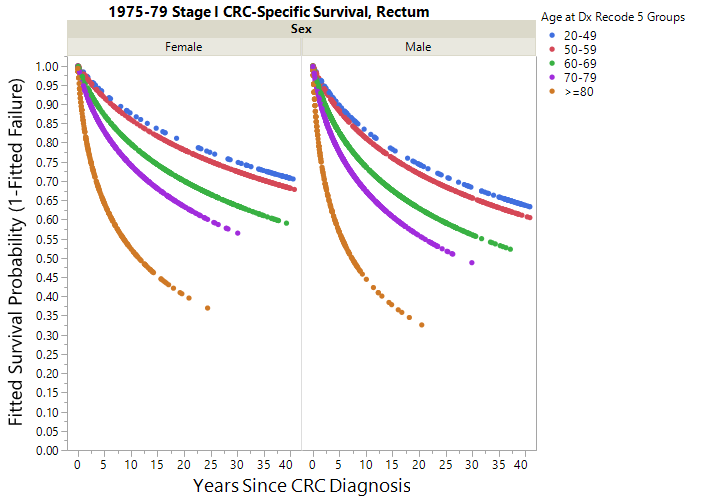

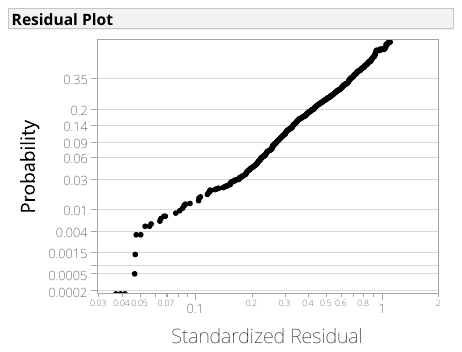

**Figure S8. Modeled cause-specific survival for 1975-79 stage II colorectal cancer (CRC) in the colon, stratified by age at diagnosis and sex.** CRC-specific survival is represented as percent survival by years since diagnosis. Note: The abnormal residual behavior at low probability is due to recoding survival month from 0 months to 0.5 months—the abnormal plot observations have survival month values of 0.5—and has little practical impact on survival, which is based on yearly increments.

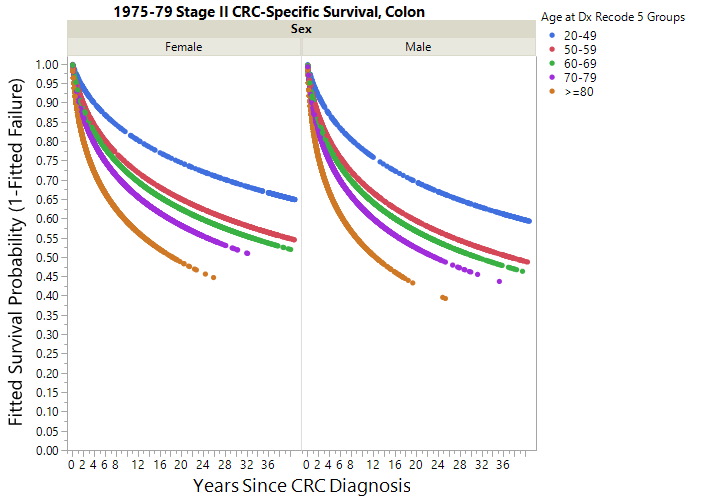

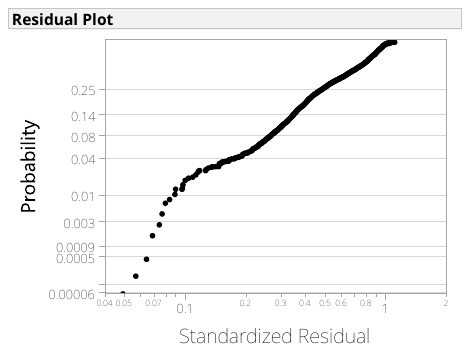

**Figure S9. Modeled cause-specific survival for 1975-79 stage II colorectal cancer (CRC) in the rectum, stratified by age at diagnosis and sex.** CRC-specific survival is represented as percent survival by years since diagnosis. Note: The abnormal residual behavior at low probability is due to recoding survival month from 0 months to 0.5 months—the abnormal plot observations have survival month values of 0.5—and has little practical impact on survival, which is based on yearly increments.

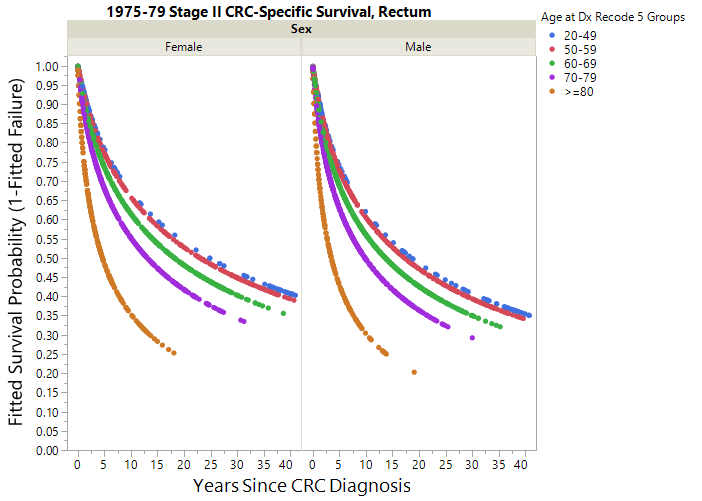

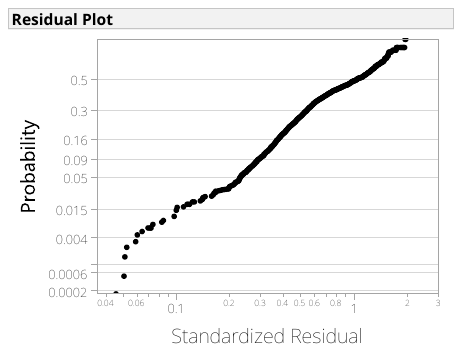
**Figure S10. Modeled cause-specific survival for 1975-79 stage III colorectal cancer (CRC) in the colon, stratified by age at diagnosis.** CRC-specific survival is represented as percent survival by years since diagnosis. Note: The abnormal residual behavior at low probability is due to recoding survival month from 0 months to 0.5 months—the abnormal plot observations have survival month values of 0.5—and has little practical impact on survival, which is based on yearly increments.

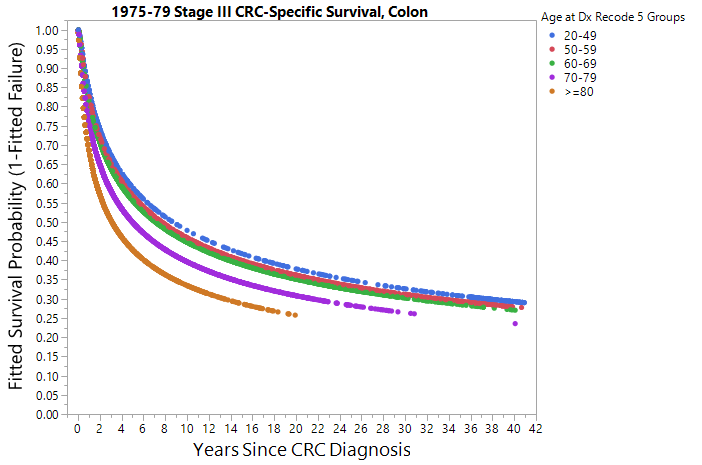

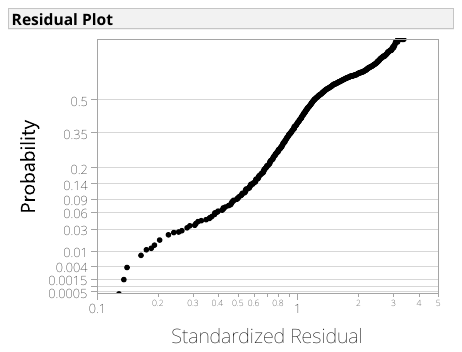
**Figure S11. Modeled cause-specific survival for 1975-79 stage III colorectal cancer (CRC) in the rectum, stratified by age at diagnosis and sex.** CRC-specific survival is represented as percent survival by years since diagnosis. Note: The abnormal residual behavior at low probability is due to recoding survival month from 0 months to 0.5 months—the abnormal plot observations have survival month values of 0.5—and has little practical impact on survival, which is based on yearly increments.
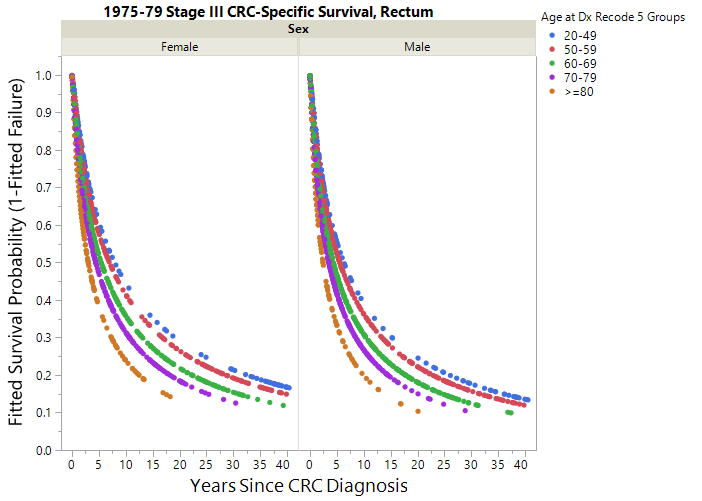

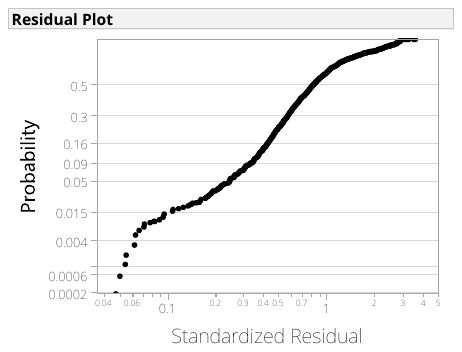
**Figure S12. Modeled cause-specific survival for 1975-79 stage IV colorectal cancer (CRC) in the colon, stratified by age at diagnosis and sex.** CRC-specific survival is represented as percent survival by years since diagnosis. Note: The abnormal residual behavior at low probability is due to recoding survival month from 0 months to 0.5 months—the abnormal plot observations have survival month values of 0.5—and has little practical impact on survival, which is based on yearly increments.

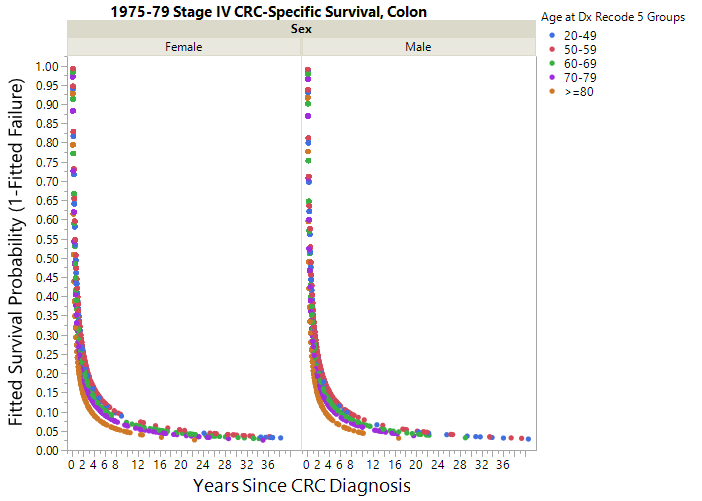

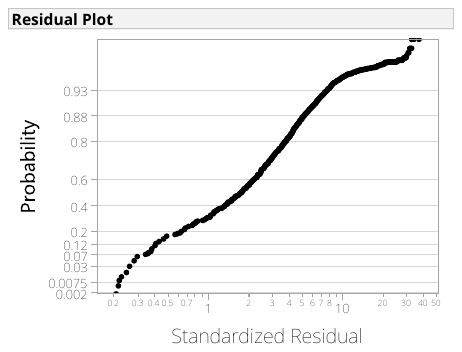
**Figure S13. Modeled cause-specific survival for 1975-79 stage IV colorectal cancer (CRC) in the rectum, stratified by age at diagnosis.** CRC-specific survival is represented as percent survival by years since diagnosis. Note: The abnormal residual behavior at low probability is due to recoding survival month from 0 months to 0.5 months—the abnormal plot observations have survival month values of 0.5—and has little practical impact on survival, which is based on yearly increments.

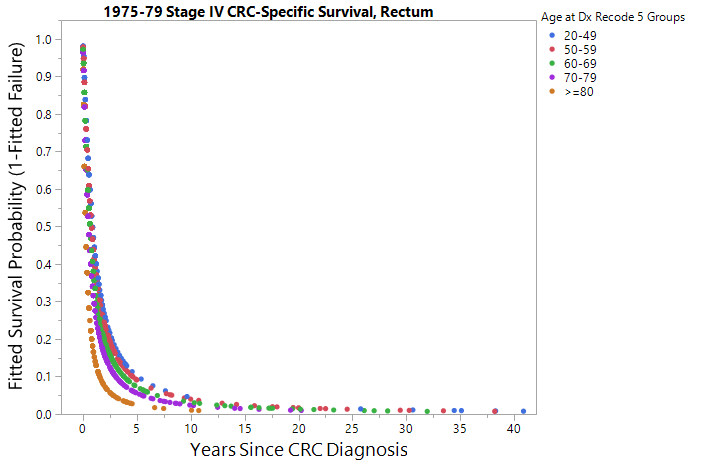

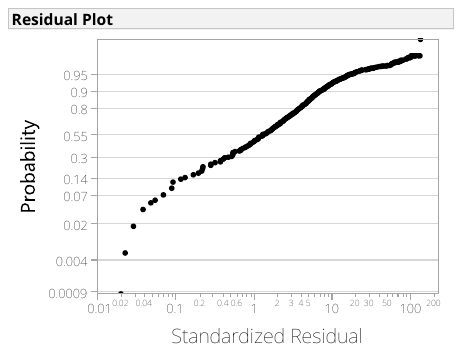

**Figure S14. Modeled cause-specific survival for 2000-03 stage I colorectal cancer (CRC) in the colon, stratified by age at diagnosis and sex.** CRC-specific survival is represented as percent survival by years since diagnosis. Note: The abnormal residual behavior at low probability is due to recoding survival month from 0 months to 0.5 months—the abnormal plot observations have survival month values of 0.5—and has little practical impact on survival, which is based on yearly increments.

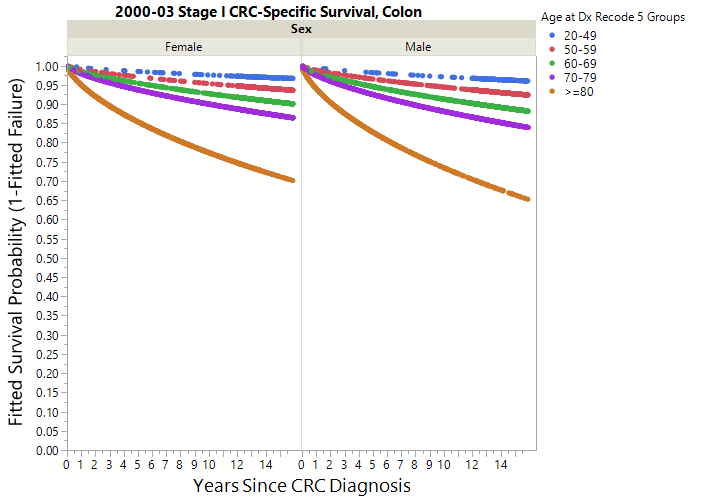

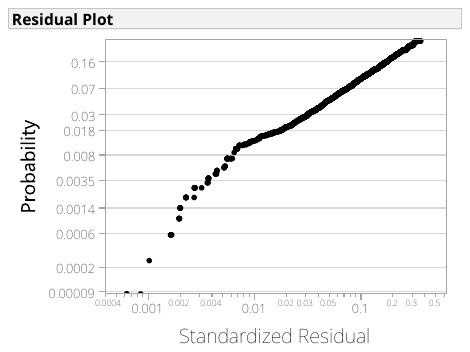
**Figure S15. Modeled cause-specific survival for 2000-03 stage I colorectal cancer (CRC) in the rectum, stratified by age at diagnosis.** CRC-specific survival is represented as percent survival by years since diagnosis. Note: The abnormal residual behavior at low probability is due to recoding survival month from 0 months to 0.5 months—the abnormal plot observations have survival month values of 0.5—and has little practical impact on survival, which is based on yearly increments.

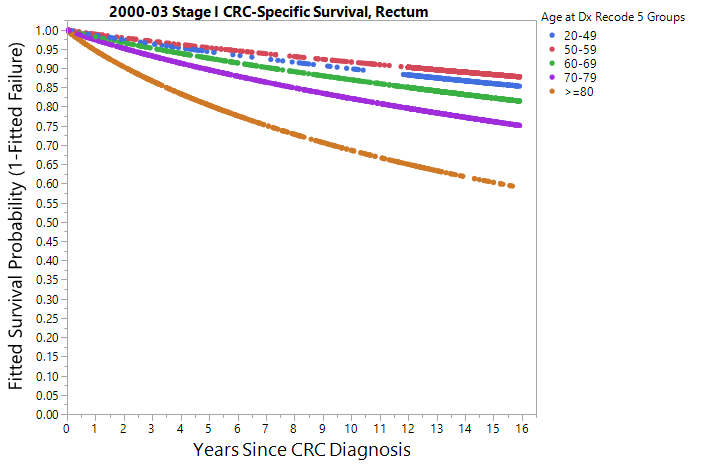

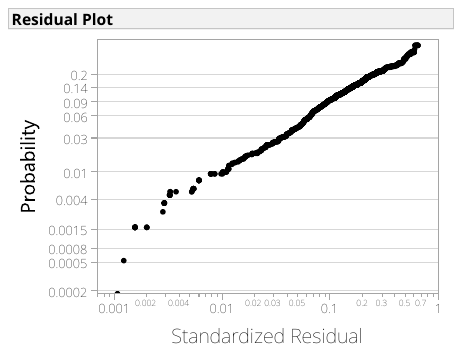
**Figure S16. Modeled cause-specific survival for 2000-03 stage II colorectal cancer (CRC) in the colon, stratified by age at diagnosis and sex.** CRC-specific survival is represented as percent survival by years since diagnosis. Note: The abnormal residual behavior at low probability is due to recoding survival month from 0 months to 0.5 months—the abnormal plot observations have survival month values of 0.5—and has little practical impact on survival, which is based on yearly increments.

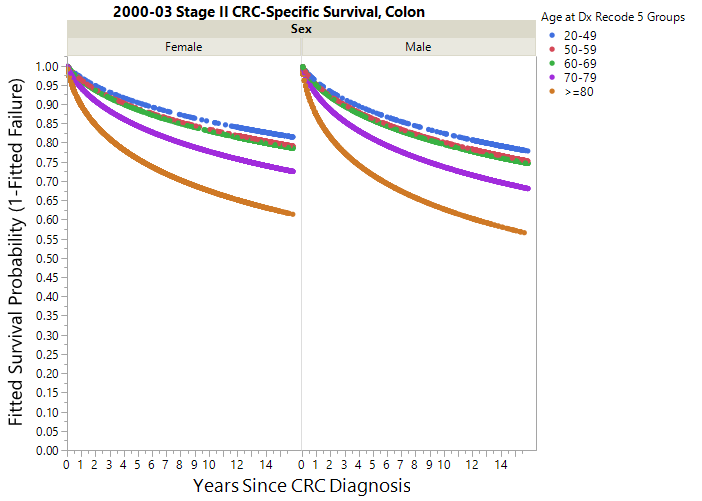

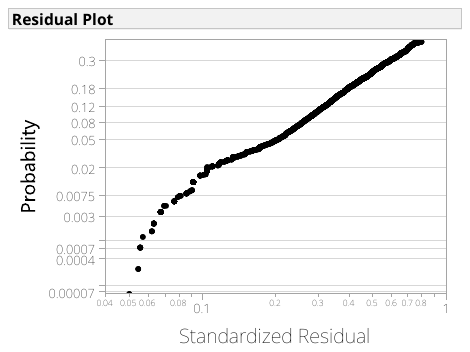

**Figure S17. Modeled cause-specific survival for 2000-03 stage II colorectal cancer (CRC) in the rectum, stratified by age at diagnosis.** CRC-specific survival is represented as percent survival by years since diagnosis. Note: The abnormal residual behavior at low probability is due to recoding survival month from 0 months to 0.5 months—the abnormal plot observations have survival month values of 0.5—and has little practical impact on survival, which is based on yearly increments.

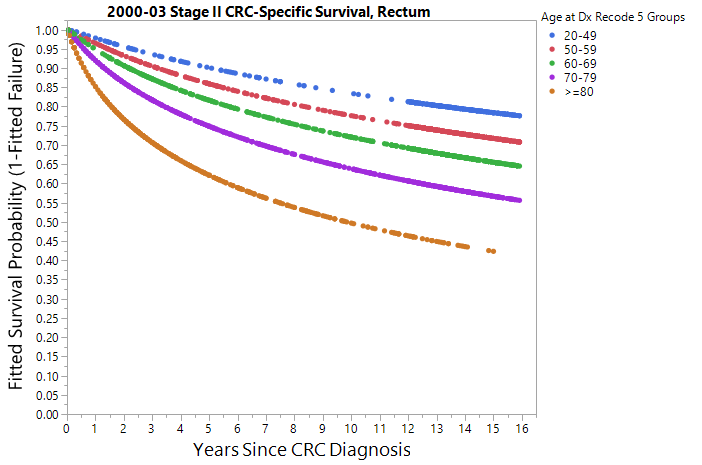

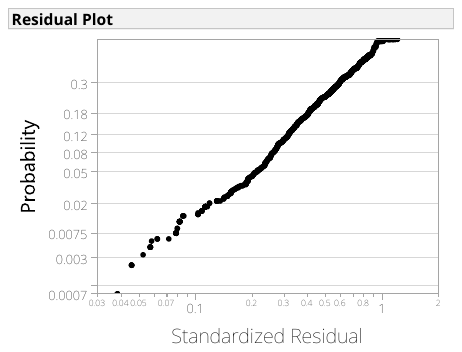

**Figure S18. Modeled cause-specific survival for 2000-03 stage III colorectal cancer (CRC) in the colon, stratified by age at diagnosis and sex.** CRC-specific survival is represented as percent survival by years since diagnosis. Note: The abnormal residual behavior at low probability is due to recoding survival month from 0 months to 0.5 months—the abnormal plot observations have survival month values of 0.5—and has little practical impact on survival, which is based on yearly increments.

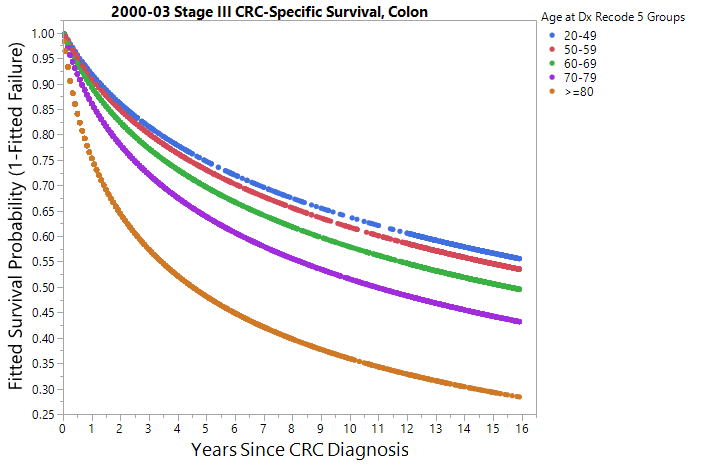

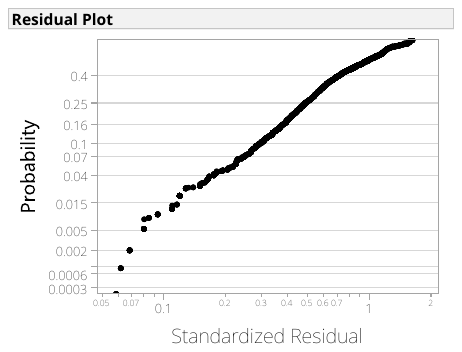

**Figure S19. Modeled cause-specific survival for 2000-03 stage III colorectal cancer (CRC) in the rectum, stratified by age at diagnosis.** CRC-specific survival is represented as percent survival by years since diagnosis. Note: The abnormal residual behavior at low probability is due to recoding survival month from 0 months to 0.5 months—the abnormal plot observations have survival month values of 0.5—and has little practical impact on survival, which is based on yearly increments.
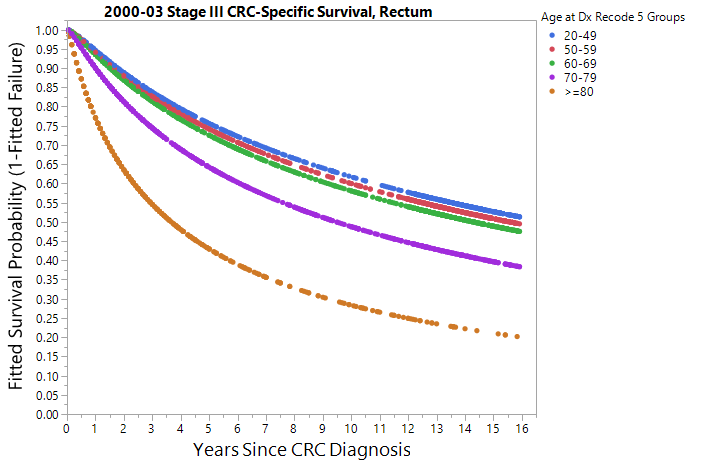

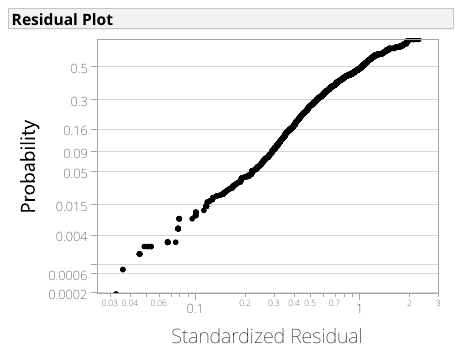

**Figure S20. Modeled cause-specific survival for 2000-03 stage IV colorectal cancer (CRC) in the colon, stratified by age at diagnosis.** CRC-specific survival is represented as percent survival by years since diagnosis. Note: The abnormal residual behavior at low probability is due to recoding survival month from 0 months to 0.5 months—the abnormal plot observations have survival month values of 0.5—and has little practical impact on survival, which is based on yearly increments.
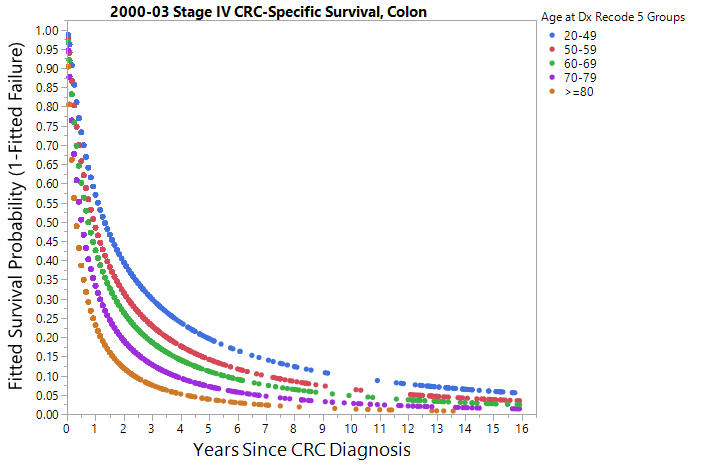

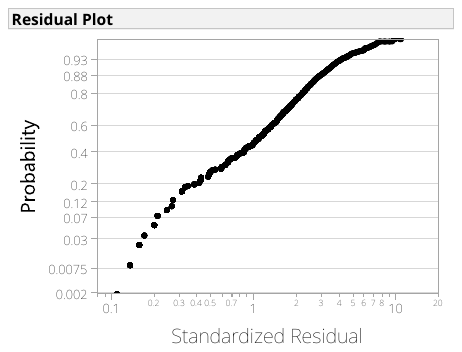

**Figure S21. Modeled cause-specific survival for 2000-03 stage IV colorectal cancer (CRC) in the rectum, stratified by age at diagnosis.** CRC-specific survival is represented as percent survival by years since diagnosis.
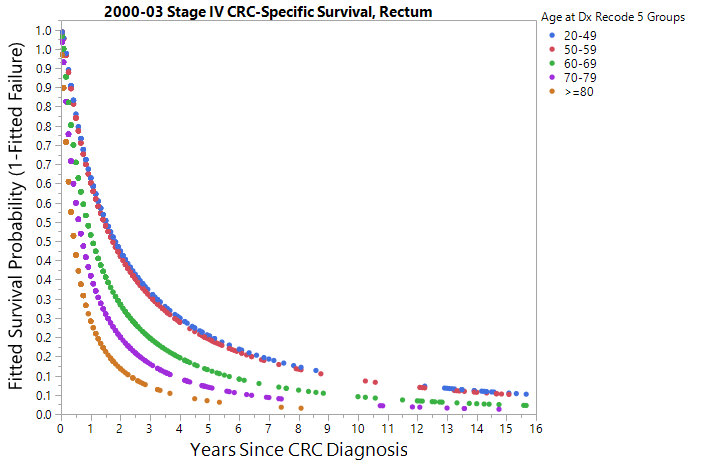

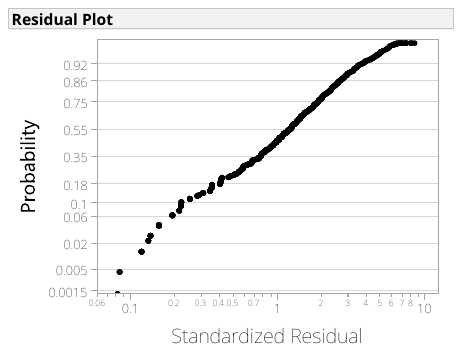

**Figure S22. Smoothed empirical distribution of American Joint Committee on Cancer (AJCC) stage vs size for different SEER datasets.** The plots were generated from SEER data for the following date collection ranges: (A) 1975-1979; (B) 1988-1992; and (C) 2011-2015. For (B) and (C), the AJCC stage percentages for CRC sizes from 98 mm to 998 mm were combined and coded as 98 mm (symbolized as an open diamond) to visually match (A).

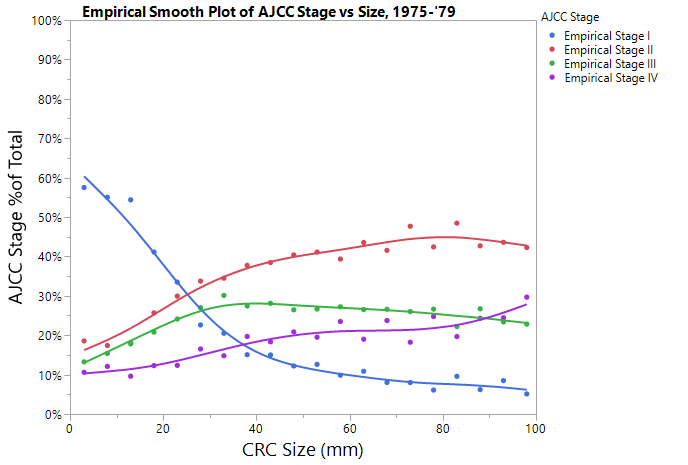

**(A)**

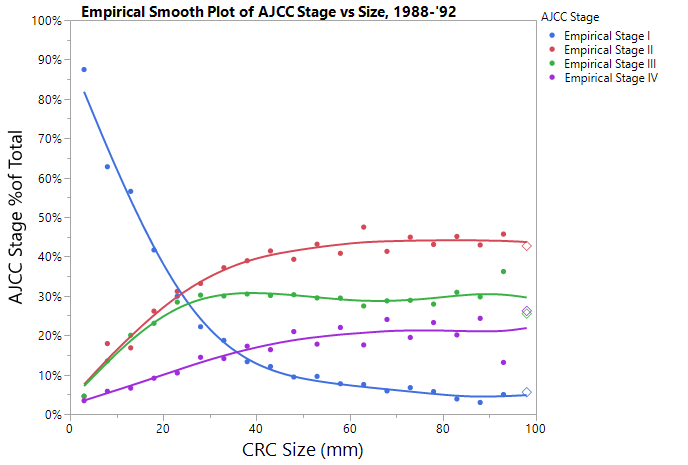

**(C)**

**(B)**

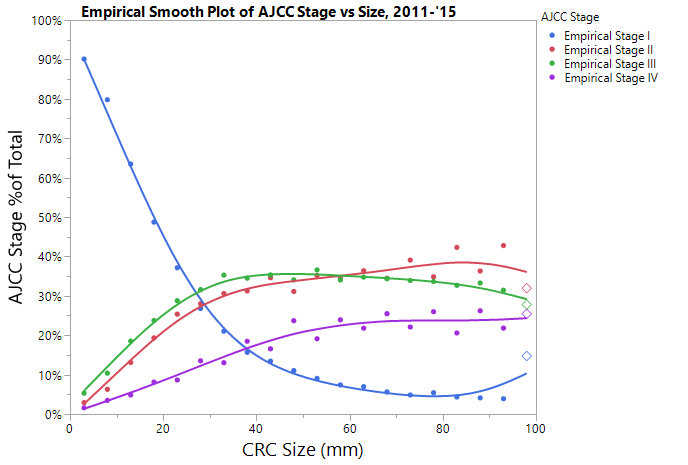

**Figure S23. Multinomial logistic regression function overlaying the 1975-1979 SEER data.** The vertical red line indicates the threshold for model-predicted stage distribution for CRC-AIM’s extrapolated CRC sizes.

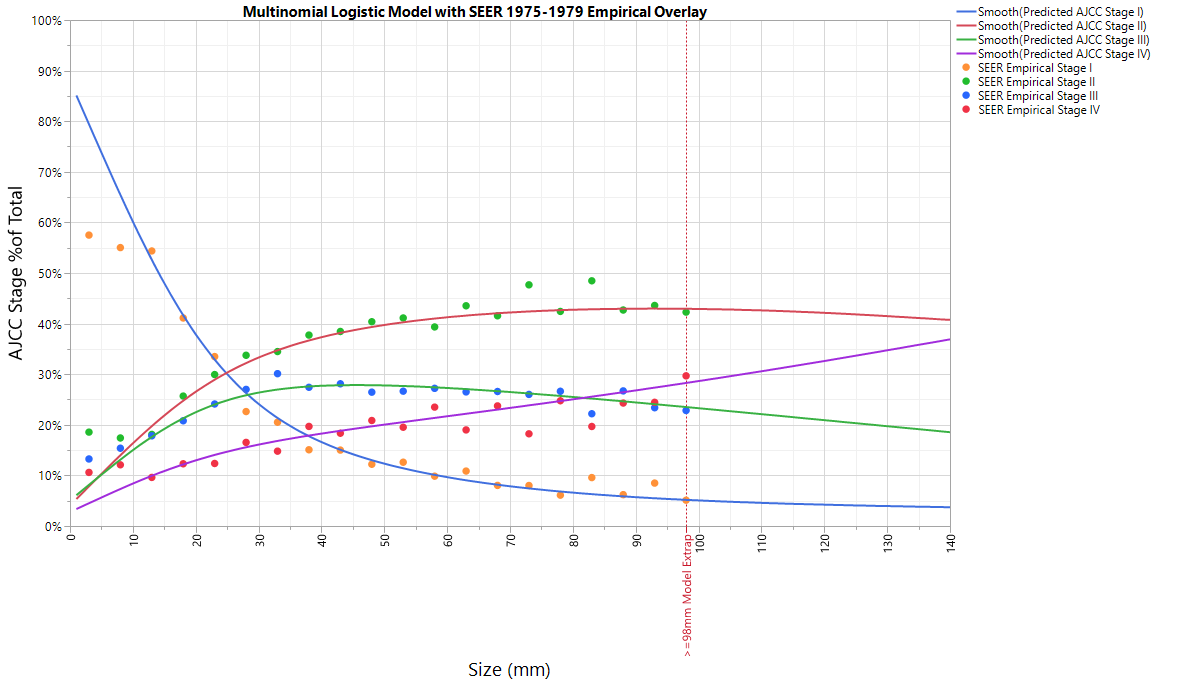

**Figure S24. Receiver-operating characteristic (ROC) analysis and associated area under the curve for the final CRC-AIM multinomial logistic regression model.**

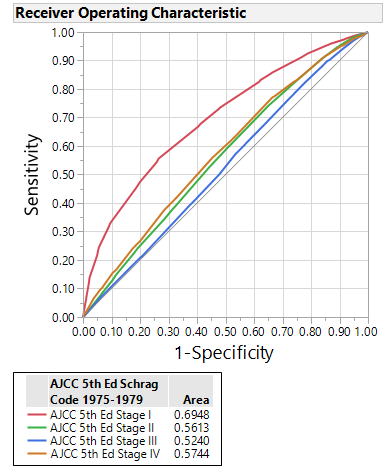

**Figure S25.** **Histogram of clinically diagnosed colorectal cancer (CRC) sizes from 1975-1979 SEER database.** Tumors within the “≥98 mm” category are represented by the patterned bar.

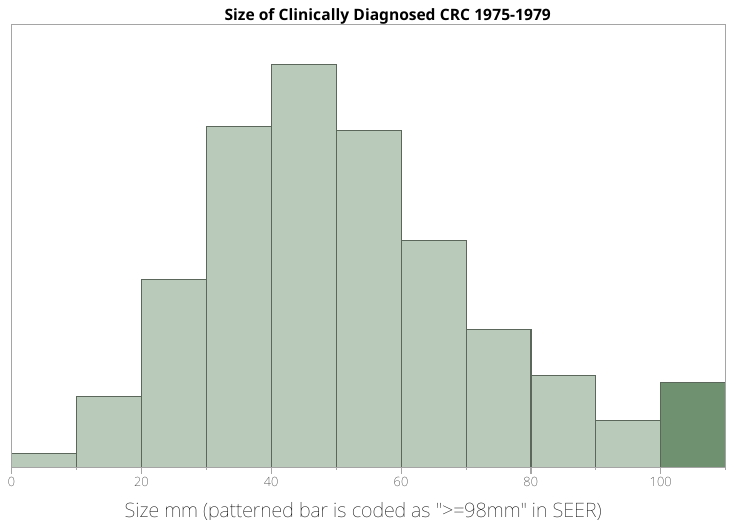

**Figure S26. Count of clinically diagnosed colorectal cancer (CRC) sizes from 1975-1979 SEER database.** Only CRCs between 50 mm and 97 mm are included.

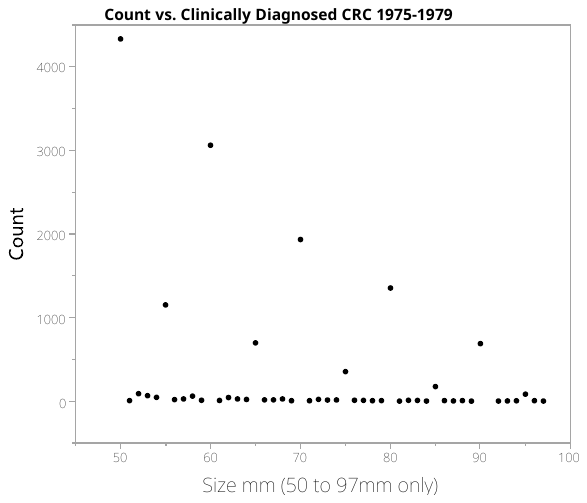

**Figure S27. Poisson regression models for colorectal cancer (CRC) size distribution rounding scenarios.** Models extrapolating sizes based on (A) rounding to the nearest centimeter (eg, 50 mm, 60 mm, 70 mm, etc.); (B) rounding to the nearest half-centimeter (eg, 55 mm, 65 mm, 75 mm, etc.), excluding scenario (A); and (C) rounding to the nearest millimeter, excluding scenarios (A) and (B).

**(A)**

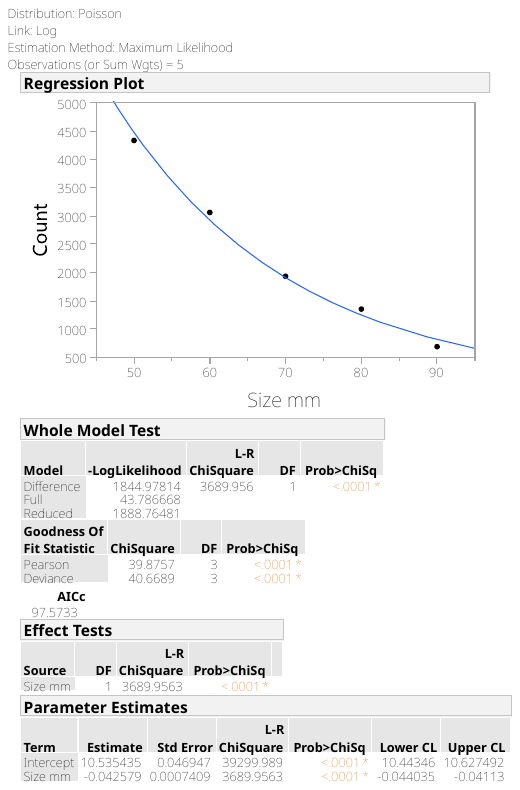

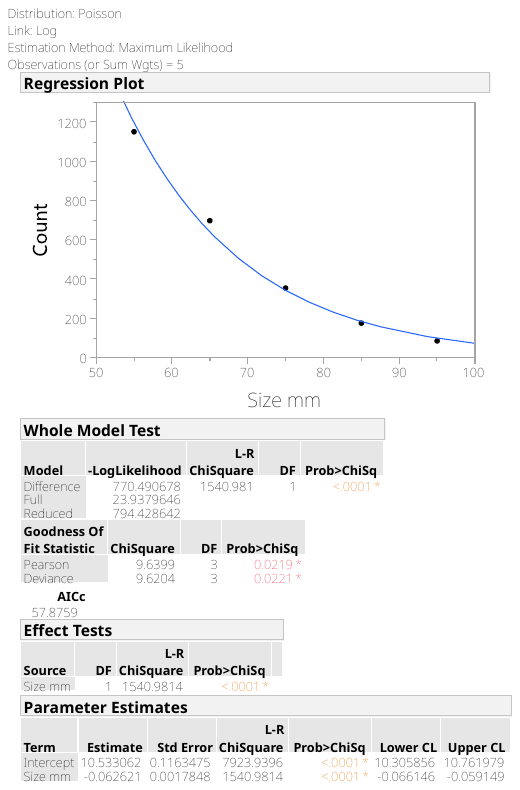

**(B)**

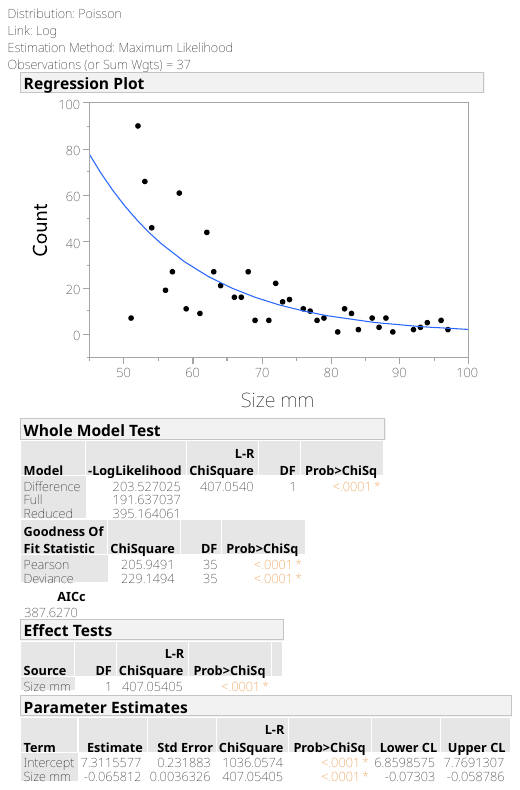

**(C)**

**Figure S28. Final model size distribution of colorectal cancer (CRC) at clinical detection for CRC-AIM.**

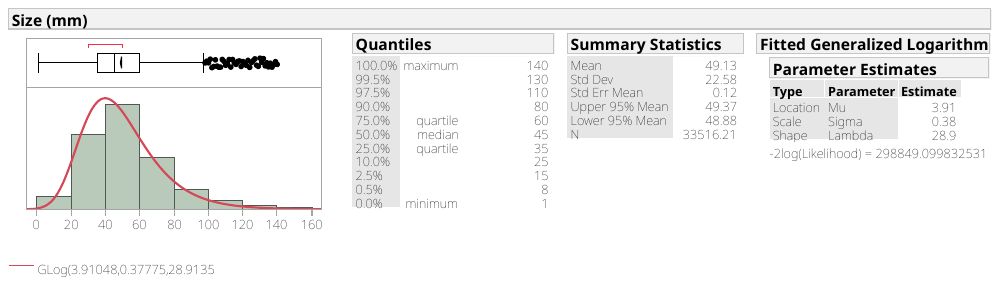

**Table S1. Colorectal cancer cause-specific survival models by location and stage.** Stage determined by the American Joint Committee on Cancer (AJCC) staging system.

| **Location, Stage** | **1975-1979** | | **2000-2003** | |
| --- | --- | --- | --- | --- |
|  | **Selected model** | **Significant effects** | **Selected model** | **Significant effects** |
| Colon, Stage I | Weibull | Sex, Age at Dx | Weibull | Sex, Age at Dx |
| Rectum, Stage I | Lognormal | Sex, Age at Dx | Loglogistic | Age at Dx |
| Colon, Stage II | Lognormal | Sex, Age at Dx | Lognormal | Sex, Age at Dx |
| Rectum, Stage II | Lognormal | Sex, Age at Dx | Lognormal | Age at Dx |
| Colon, Stage III | Fréchet | Age at Dx | Lognormal | Age at Dx |
| Rectum, Stage III | Lognormal | Sex, Age at Dx | Lognormal | Age at Dx |
| Colon, Stage IV | Fréchet | Sex, Age at Dx | Lognormal | Age at Dx |
| Rectum, Stage IV | Loglogistic | Age at Dx | Lognormal | Age at Dx |

**Table S2.** **Dwell time summary statistics by model.** Data from 2011 adapted from Kuntz et al;^6^ data from 2010 adapted from Rutter et al.^3^ IQR = interquartile range; n/a = not available

NOTE: MISCAN was recalibrated in 2014 and therefore these dwell time measures are no longer applicable. The dwell time summary statistics for the MISCAN’s recalibration were not described in their publication.^19^

| **Measure** | **Statistic** | **MISCAN (2011)** | **CRC-SPIN (2011)** | **SimCRC (2011)** | **CRC-SPIN (2010)** | **CRC-AIM** |
| --- | --- | --- | --- | --- | --- | --- |
| **Adenoma dwell time, y** | Mean | 7.6 | 24.2 | 21.2 | 25.4 | 23.5 |
|  | Median | 6.0 | 23.0 | 19.0 | n/a | 22.0 |
|  | IQR | 2 - 11 | 16 - 31 | 12 - 29 | n/a | 14.0 - 31.0 |
| **Sojourn time (preclinical cancer dwell time), y** | Mean | 3.0 | 1.6 | 4.0 | 1.9 | 2.9 |
|  | Median | 2.0 | 2.0 | 3.0 | n/a | 2.6 |
|  | IQR | 1 - 4 | 1 - 2 | 2 - 5 | n/a | 1.8 - 3.6 |
| **Overall dwell time (full adenoma-carcinoma sequence), y** | Mean | 10.6 | 25.8 | 25.2 | 27.2 | 26.4 |
|  | Median | 9.0 | 24.0 | 23.0 | n/a | 24.5 |
|  | IQR | 5 - 14 | 17 - 33 | 15-33 | n/a | 16.9 - 34.1 |

**Table S3.** **Derived transition probabilities for a simulated cohort of 60-year-olds.** Data adapted from Rutter et al.^3^

NOTE: CRC-SPIN results were generated from 1,000 draws across joint posterior probability distributions for all model parameters. For each draw, 30,000,000 individuals were simulated. Additionally, individuals were assigned their most advanced disease state at each time point. Individuals who died during the one-year period are included in their most recent state. CI = credible interval

| **Most Advanced Lesion State** | | **CRC-SPIN** | **CRC-AIM** |
| --- | --- | --- | --- |
| **At time 0**  **(60 years old)** | **One year later**  **(61 years old)** | **Transition probability**  **estimate (95% CI)** | **Transition probability**  **estimate** |
| **No adenomas** | No adenomas | 0.975 (0.969-0.979) | 0.9740 |
|  | ≥1 small adenoma | 0.013 (0.009-0.019) | 0.0128 |
|  | ≥1 medium adenoma | <10^-8^ (--) | 0.0000 |
|  | ≥1 large adenoma | <10^-8^ (--) | 0.0000 |
|  | Preclinical CRC | <10^-8^ (--) | 0.0000 |
|  | Clinically detected CRC | 0 (--) | 0.0000 |
|  | CRC death | 0 (--) | 0.0000 |
|  | Non-CRC death | 0.012 (0.012-0.013) | 0.0132 |
| **≥1 small adenoma** | ≥1 small adenoma | 0.925 (0.909-0.942) | 0.9219 |
|  | ≥1 medium adenoma | 0.061 (0.044-0.078) | 0.0640 |
|  | ≥1 large adenoma | <10^-8^ (--) | 0.0000 |
|  | Preclinical CRC | 0.00014 (0.00008-0.00021) | 0.0002 |
|  | Clinically detected CRC | 0.000002 (0.00000-0.00004) | 0.0000 |
|  | CRC death | <10^-8^ (--) | 0.0000 |
|  | Non-CRC death | 0.013 (0.013-0.013) | 0.0139 |
| **≥1 medium adenoma** | ≥1 medium adenoma | 0.932 (0.922-0.940) | 0.9245 |
|  | ≥1 large adenoma | 0.051 (0.043-0.061) | 0.0579 |
|  | Preclinical CRC | 0.003 (0.002-0.004) | 0.0035 |
|  | Clinically detected CRC | 0.0004 (0.0000-0.0010) | 0.0001 |
|  | CRC death | 0.00002 (0.00000-0.00006) | 0.0000 |
|  | Non-CRC death | 0.013 (0.013-0.013) | 0.0140 |
| **≥1 large adenoma** | ≥1 large adenoma | 0.970 (0.966-0.973) | 0.9709 |
|  | Preclinical CRC | 0.015 (0.011-0.019) | 0.0143 |
|  | Clinically detected CRC | 0.002 (0.000-0.004) | 0.0004 |
|  | CRC death | 0.00008 (0.00000-0.00024) | 0.0000 |
|  | Non-CRC death | 0.013 (0.013-0.014) | 0.0144 |
| **Preclinical CRC** | Preclinical CRC | 0.566 (0.351-0.719) | 0.6339 |
|  | Clinically detected CRC | 0.381 (0.244-0.575) | 0.3078 |
|  | CRC death | 0.039 (0.024-0.060) | 0.0451 |
|  | Non-CRC death | 0.013 (0.012-0.014) | 0.0132 |
| **Clinically detected CRC** | Clinically detected CRC | 0.924 (0.921-0.927) | 0.9314 |
|  | CRC death | 0.063 (0.060-0.067) | 0.0553 |
|  | Non-CRC death | 0.012 (0.012-0.013) | 0.0133 |

**Table S4.** **Adenoma prevalence by age.** From Figure 4B.

| **Age** | **CRC-AIM Adenoma Prevalence** |
| --- | --- |
| 40 | 10.1% |
| 50 | 16.9% |
| 60 | 24.6% |
| 70 | 32.8% |
| 80 | 40.8% |
| 90 | 47.8% |

**Table S5.** **Summary of CRC-AIM components and parameter estimates.** Source data from Rutter et al.^5^

| **Component** | **Symbol** | **Estimate** | **Source** |
| --- | --- | --- | --- |
| **Adenoma risk** | | | |
| Baseline log-risk | α_0_ | -6.6 | Rutter 2009 |
| Main sex effect | α_1_ | -0.24 | Rutter 2009 |
| Standard deviation baseline log-risk | σ_α_ | 1.1 | Rutter 2009 |
| Age effect, age ε [20,50) | α_20_ | 0.037 | Rutter 2009 |
| Age effect, age ε [50,60) | α_21_ | 0.031 | Rutter 2009 |
| Age effect, age ε [60,70) | α_22_ | 0.029 | Rutter 2009 |
| Age effect, age ≥ 70 | α_23_ | 0.030 | Rutter 2009 |
| **Adenoma growth: Time to 10 mm** | | | |
| Colon: Location | β_1c_ | 24.3 | Recalibration of Rutter 2009 |
| Colon: Scale | β_2c_ | 1.8472066 | Recalibration of Rutter 2009 |
| Rectum: Location | β_1r_ | 11.086583 | Recalibration of Rutter 2009 |
| Rectum: Scale | β_2r_ | 2.4485203 | Recalibration of Rutter 2009 |
| **Transition to cancer** | | | |
| Men, Colon, size | γ_1_*_cm_* | 0.0400000 | Recalibration of Rutter 2009 |
| Men, Colon, age at initiation | γ_2_*_cm_* | 0.0089232 | Recalibration of Rutter 2009 |
| Men, Rectum, size | γ_1_*_rm_* | 0.0472322 | Recalibration of Rutter 2009 |
| Men, Rectum, age at initiation | γ_2_*_rm_* | 0.0173598 | Recalibration of Rutter 2009 |
| Women, Colon, size | γ_1_*_cf_* | 0.0444762 | Recalibration of Rutter 2009 |
| Women, Colon, age at initiation | γ_2_*_cf_* | 0.0089362 | Recalibration of Rutter 2009 |
| Women, Rectum, size | γ_1_*_rf_* | 0.0470747 | Recalibration of Rutter 2009 |
| Women, Rectum, age at initiation | γ_2_*_rf_* | 0.0161731 | Recalibration of Rutter 2009 |
| Shape parameter | $\gamma_{3}$ | 0.5 | Rutter 2009 |
| **Mean sojourn time** | | | |
| Colon | μ_c_ | 2.9 | Uncalibrated; shifted value from Rutter 2009 (1.9) |
|  | τ_c_ | 0.5241379 | Set so standard deviation of distribution matched Rutter 2009 (1.52) |
| Rectum | μ_r_ | 3.7 | Uncalibrated; shifted value from Rutter 2009 (2.7) |
|  | τ_r_ | 0.612973 | Set so standard deviation of distribution matched Rutter 2009 (2.268) |
| **Clinically detected CRC size** | | | |
| Location | μ | 3.91048 | Derived from SEER |
| Scale | σ | 0.37775 | Derived from SEER |
| Shape | λ | 28.9135 | Derived from SEER |
| **Multinomial logistic regression CRC stage \| size** | | | |
| Stage I | α_1_ | 4.45118078 | Derived from SEER |
|  | β**_1_** | -0.8845893 | Derived from SEER |
|  | γ_1_ | 0.02665437 | Derived from SEER |
| Stage II | α_2_ | 0.29839421 | Derived from SEER |
|  | β_2_ | 0.16069904 | Derived from SEER |
|  | γ_2_ | -0.015023 | Derived from SEER |
| Stage III | α_3_ | 0.41562304 | Derived from SEER |
|  | β**_3_** | 0.10670585 | Derived from SEER |
|  | γ_3_ | -0.0169 | Derived from SEER |
| **CRC cause-specific survival, 1975-1979** | | | |
| Stage I Colon (1975-1979) | | | |
|  | k_Ic_ | 0.7549 | Derived from SEER |
| Intercept | Intercept_Ic_ | 4.7875 | Derived from SEER |
| Age effect, age ε [20,50) | A_20-49_Ic_ | 0.8587 | Derived from SEER |
| Age effect, age ε [50,60) | A_50-59_Ic_ | 0.5867 | Derived from SEER |
| Age effect, age ε [60,70) | A_60-69_Ic_ | 0.1352 | Derived from SEER |
| Age effect, age ε [70,80) | A_70-79_Ic_ | -0.3548 | Derived from SEER |
| Age effect, age ≥ 80 | A_80-∞_Ic_ | -1.2257 | Derived from SEER |
| Main sex effect | SexF_Ic_ | 0.2002 | Derived from SEER |
| Stage I Rectum (1975-1979) | | | |
|  | σ_Ir_ | 2.2738 | Derived from SEER |
| Intercept | Intercept_Ir_ | 3.7957 | Derived from SEER |
| Age effect, age ε [20,50) | A_20-49_Ir_ | 0.9111 | Derived from SEER |
| Age effect, age ε [50,60) | A_50-59_Ir_ | 0.7413 | Derived from SEER |
| Age effect, age ε [60,70) | A_60-69_Ir_ | 0.1719 | Derived from SEER |
| Age effect, age ε [70,80) | A_70-79_Ir_ | -0.2448 | Derived from SEER |
| Age effect, age ≥ 80 | A_80-∞_Ir_ | -1.5794 | Derived from SEER |
| Main sex effect | SexF_Ir_ | 0.2227 | Derived from SEER |
| Stage II Colon (1975-1979) | | | |
|  | σ_IIc_ | 2.6110 | Derived from SEER |
| Intercept | Intercept_IIc_ | 3.6001 | Derived from SEER |
| Age effect, age ε [20,50) | A_20-49_IIc_ | 0.9136 | Derived from SEER |
| Age effect, age ε [50,60) | A_50-59_IIc_ | 0.2072 | Derived from SEER |
| Age effect, age ε [60,70) | A_60-69_IIc_ | 0.0266 | Derived from SEER |
| Age effect, age ε [70,80) | A_70-79_IIc_ | -0.2606 | Derived from SEER |
| Age effect, age ≥ 80 | A_80-∞_IIc_ | -0.8869 | Derived from SEER |
| Main sex effect | SexF_IIc_ | 0.1928 | Derived from SEER |
| Stage II Rectum (1975-1979) | | | |
|  | σ_IIr_ | 2.0426 | Derived from SEER |
| Intercept | Intercept_IIr_ | 2.5271 | Derived from SEER |
| Age effect, age ε [20,50) | A_20-49_IIr_ | 0.5389 | Derived from SEER |
| Age effect, age ε [50,60) | A_50-59_IIr_ | 0.4658 | Derived from SEER |
| Age effect, age ε [60,70) | A_60-69_IIr_ | 0.2293 | Derived from SEER |
| Age effect, age ε [70,80) | A_70-79_IIr_ | -0.0995 | Derived from SEER |
| Age effect, age ≥ 80 | A_80-∞_IIr_ | -1.1344 | Derived from SEER |
| Main sex effect | SexF_IIr_ | 0.1430 | Derived from SEER |
| Stage III Colon (1975-1979) | | | |
|  | σ_IIIc_ | 2.20035 | Derived from SEER |
| Intercept | Intercept_IIIc_ | 0.97290 | Derived from SEER |
| Age effect, age ε [20,50) | A_20-49_IIIc_ | 0.38287 | Derived from SEER |
| Age effect, age ε [50,60) | A_50-59_IIIc_ | 0.25911 | Derived from SEER |
| Age effect, age ε [60,70) | A_60-69_IIIc_ | 0.17891 | Derived from SEER |
| Age effect, age ε [70,80) | A_70-79_IIIc_ | -0.17465 | Derived from SEER |
| Age effect, age ≥ 80 | A_80-∞_IIIc_ | -0.64624 | Derived from SEER |
| Stage III Rectum (1975-1979) | | | |
|  | σ_IIIr_ | 1.6754 | Derived from SEER |
| Intercept | Intercept_IIIr_ | 1.5492 | Derived from SEER |
| Age effect, age ε [20,50) | A_20-49_IIIr_ | 0.4097 | Derived from SEER |
| Age effect, age ε [50,60) | A_50-59_IIIr_ | 0.2800 | Derived from SEER |
| Age effect, age ε [60,70) | A_60-69_IIIr_ | 0.0311 | Derived from SEER |
| Age effect, age ε [70,80) | A_70-79_IIIr_ | -0.1718 | Derived from SEER |
| Age effect, age ≥ 80 | A_80-∞_IIIr_ | -0.5490 | Derived from SEER |
| Main sex effect | SexF_IIIr_ | 0.1138 | Derived from SEER |
| Stage IV Colon (1975-1979) | | | |
|  | σ_IVc_ | 1.37264 | Derived from SEER |
| Intercept | Intercept_IVc_ | -1.36191 | Derived from SEER |
| Age effect, age ε [20,50) | A_20-49_IVc_ | 0.25951 | Derived from SEER |
| Age effect, age ε [50,60) | A_50-59_IVc_ | 0.31202 | Derived from SEER |
| Age effect, age ε [60,70) | A_60-69_IVc_ | 0.06888 | Derived from SEER |
| Age effect, age ε [70,80) | A_70-79_IVc_ | -0.11106 | Derived from SEER |
| Age effect, age ≥ 80 | A_80-∞_IVc_ | -0.52935 | Derived from SEER |
| Main sex effect | SexF_IVc_ | 0.03688 | Derived from SEER |
| Stage IV Rectum (1975-1979) | | | |
|  | σ_IVr_ | 0.7855 | Derived from SEER |
| Intercept | Intercept_IVr_ | -0.5094 | Derived from SEER |
| Age effect, age ε [20,50) | A_20-49_IVr_ | 0.4154 | Derived from SEER |
| Age effect, age ε [50,60) | A_50-59_IVr_ | 0.3168 | Derived from SEER |
| Age effect, age ε [60,70) | A_60-69_IVr_ | 0.1257 | Derived from SEER |
| Age effect, age ε [70,80) | A_70-79_IVr_ | -0.0969 | Derived from SEER |
| Age effect, age ≥ 80 | A_80-∞_IVr_ | -0.7610 | Derived from SEER |
| **CRC cause-specific survival, 2000-2003** | | | |
| Stage I Colon (2000-2003) | | | |
|  | k_Ic_ | 0.699099 | Derived from SEER |
| Intercept | Intercept_Ic_ | 5.8797 | Derived from SEER |
| Age effect, age ε [20,50) | A_20-49_Ic_ | 1.6097 | Derived from SEER |
| Age effect, age ε [50,60) | A_50-59_Ic_ | 0.6499 | Derived from SEER |
| Age effect, age ε [60,70) | A_60-69_Ic_ | -0.0115 | Derived from SEER |
| Age effect, age ε [70,80) | A_70-79_Ic_ | -0.4863 | Derived from SEER |
| Age effect, age ≥ 80 | A_80-∞_Ic_ | -1.7619 | Derived from SEER |
| Main sex effect | SexF_Ic_ | 0.1321 | Derived from SEER |
| Stage I Rectum (2000- 2003) | | | |
|  | σ_Ir_ | 1.1079 | Derived from SEER |
| Intercept | Intercept_Ir_ | 4.2475 | Derived from SEER |
| Age effect, age ε [20,50) | A_20-49_Ir_ | 0.4703 | Derived from SEER |
| Age effect, age ε [50,60) | A_50-59_Ir_ | 0.7033 | Derived from SEER |
| Age effect, age ε [60,70) | A_60-69_Ir_ | 0.1593 | Derived from SEER |
| Age effect, age ε [70,80) | A_70-79_Ir_ | -0.2561 | Derived from SEER |
| Age effect, age ≥ 80 | A_80-∞_Ir_ | -1.0768 | Derived from SEER |
| Stage II Colon (2000-2003) | | | |
|  | σ_IIc_ | 2.8316 | Derived from SEER |
| Intercept | Intercept_IIc_ | 4.5013 | Derived from SEER |
| Age effect, age ε [20,50) | A_20-49_IIc_ | 0.6193 | Derived from SEER |
| Age effect, age ε [50,60) | A_50-59_IIc_ | 0.3778 | Derived from SEER |
| Age effect, age ε [60,70) | A_60-69_IIc_ | 0.3243 | Derived from SEER |
| Age effect, age ε [70,80) | A_70-79_IIc_ | -0.2226 | Derived from SEER |
| Age effect, age ≥ 80 | A_80-∞_IIc_ | -1.0987 | Derived from SEER |
| Main sex effect | SexF_IIc_ | 0.1825 | Derived from SEER |
| Stage II Rectum (2000-2003) | | | |
|  | σ_IIr_ | 2.1874 | Derived from SEER |
| Intercept | Intercept_IIr_ | 3.4680 | Derived from SEER |
| Age effect, age ε [20,50) | A_20-49_IIr_ | 0.9608 | Derived from SEER |
| Age effect, age ε [50,60) | A_50-59_IIr_ | 0.4976 | Derived from SEER |
| Age effect, age ε [60,70) | A_60-69_IIr_ | 0.1146 | Derived from SEER |
| Age effect, age ε [70,80) | A_70-79_IIr_ | -0.3914 | Derived from SEER |
| Age effect, age ≥ 80 | A_80-∞_IIr_ | -1.1817 | Derived from SEER |
| Stage III Colon (2000-2003) | | | |
|  | σ_IIIc_ | 2.2040 | Derived from SEER |
| Intercept | Intercept_IIIc_ | 2.5361 | Derived from SEER |
| Age effect, age ε [20,50) | A_20-49_IIIc_ | 0.5416 | Derived from SEER |
| Age effect, age ε [50,60) | A_50-59_IIIc_ | 0.4263 | Derived from SEER |
| Age effect, age ε [60,70) | A_60-69_IIIc_ | 0.2067 | Derived from SEER |
| Age effect, age ε [70,80) | A_70-79_IIIc_ | -0.1463 | Derived from SEER |
| Age effect, age ≥ 80 | A_80-∞_IIIc_ | -1.0283 | Derived from SEER |
| Stage III Rectum (2000-2003) | | | |
|  | σ_IIIr_ | 1.7519 | Derived from SEER |
| Intercept | Intercept_IIIr_ | 2.3558 | Derived from SEER |
| Age effect, age ε [20,50) | A_20-49_IIIr_ | 0.4700 | Derived from SEER |
| Age effect, age ε [50,60) | A_50-59_IIIr_ | 0.3899 | Derived from SEER |
| Age effect, age ε [60,70) | A_60-69_IIIr_ | 0.3039 | Derived from SEER |
| Age effect, age ε [70,80) | A_70-79_IIIr_ | -0.1072 | Derived from SEER |
| Age effect, age ≥ 80 | A_80-∞_IIIr_ | -1.0567 | Derived from SEER |
| Stage IV Colon (2000-2003) | | | |
|  | σ_IVc_ | 1.5583 | Derived from SEER |
| Intercept | Intercept_IVc_ | -0.3766 | Derived from SEER |
| Age effect, age ε [20,50) | A_20-49_IVc_ | 0.6531 | Derived from SEER |
| Age effect, age ε [50,60) | A_50-59_IVc_ | 0.3190 | Derived from SEER |
| Age effect, age ε [60,70) | A_60-69_IVc_ | 0.0861 | Derived from SEER |
| Age effect, age ε [70,80) | A_70-79_IVc_ | -0.2930 | Derived from SEER |
| Age effect, age ≥ 80 | A_80-∞_IVc_ | -0.7653 | Derived from SEER |
| Stage IV Rectum (2000-2003) | | | |
|  | σ_IVr_ | 1.4349 | Derived from SEER |
| Intercept | Intercept_IVr_ | -0.1707 | Derived from SEER |
| Age effect, age ε [20,50) | A_20-49_IVr_ | 0.5883 | Derived from SEER |
| Age effect, age ε [50,60) | A_50-59_IVr_ | 0.5408 | Derived from SEER |
| Age effect, age ε [60,70) | A_60-69_IVr_ | 0.0489 | Derived from SEER |
| Age effect, age ε [70,80) | A_70-79_IVr_ | -0.3437 | Derived from SEER |
| Age effect, age ≥ 80 | A_80-∞_IVr_ | -0.8343 | Derived from SEER |

**Table S6. Detailed adenoma statistics at age 65 for CISNET models (MISCAN, SimCRC, and CRC-SPIN) and a prototype version of CRC-AIM.** The CRC-AIM prototype was prior to recalibration. Data adapted from Knudsen et al.^20^

*The table from Knudsen et al^20^ labels this category as “1-10 mm”.

| **Outcome** | | **MISCAN** | **SimCRC** | **CRC-SPIN** | **Prototype CRC-AIM** |
| --- | --- | --- | --- | --- | --- |
| Adenoma prevalence, age 65 | | 39.80% | 37.20% | 30.70% | not calculated |
| **Number of adenomas per 1000 by site and size at age 65 y** | | | | | |
| Proximal Colon | 1-5 mm | 121.2 | 171.7 | 190.2 | 200.2 |
|  | 6-9 mm | 69.9 | 186.2 | 67.8 | 75.1 |
|  | ≥10 mm* | 61.8 | 23.9 | 40.8 | 43.6 |
| Distal Colon | 1-5 mm | 134.4 | 124.2 | 124.5 | 131.0 |
|  | 6-9 mm | 77.4 | 18.2 | 44.4 | 49.4 |
|  | ≥10 mm | 68.4 | 41.6 | 26.7 | 28.5 |
| Rectum | 1-5 mm | 133.5 | 8.7 | 14.1 | 15.5 |
|  | 6-9 mm | 76.8 | 16.0 | 9.1 | 9.4 |
|  | ≥10 mm | 68.1 | 15.8 | 20.2 | 22.6 |

**Table S7. Comparison to estimates of clinical cancers per 100,000 among SEER 1975-1979 estimates published in Rutter et al, CRC-SPIN, and a prototype version of CRC-AIM.** The CRC-AIM prototype was prior to recalibration. Data adapted from Rutter et al.^5^

| **Location, Sex** | **Age (years)** | **SEER 1975-1979, per Rutter 2009** | **CRC-SPIN (Rutter 2009)** | **Prototype**  **CRC-AIM** | **Difference between CRC-SPIN and Prototype CRC-AIM (%)** |
| --- | --- | --- | --- | --- | --- |
| Colon, Female | 20-49 | 4.8 | 4.4 | not calculated | NA |
|  | 50-59 | 43.3 | 45.5 | 41.19 | -9.48% |
|  | 60-69 | 100.7 | 99.3 | 94.63 | -4.70% |
|  | 70-84 | 216.7 | 207.3 | 198.75 | -4.12% |
| Rectum, Female | 20-49 | 1.87 | 2.0 | not calculated | NA |
|  | 50-59 | 20.4 | 18.6 | 18.45 | -0.80% |
|  | 60-69 | 42.5 | 39.7 | 41.44 | 4.38% |
|  | 70-84 | 73.9 | 82.1 | 83.58 | 1.80% |
| Colon, Male | 20-49 | 4.51 | 4.2 | not calculated | NA |
|  | 50-59 | 45.9 | 50.6 | 43.85 | -13.35% |
|  | 60-69 | 121.4 | 120.0 | 111.88 | -6.76% |
|  | 70-84 | 268.4 | 263.4 | 257.15 | -2.37% |
| Rectum, Male | 20-49 | 2.3 | 3.2 | not calculated | NA |
|  | 50-59 | 30.0 | 29.8 | 21.23 | -28.76% |
|  | 60-69 | 71.4 | 63.2 | 42.97 | -32.00% |
|  | 70-84 | 128.0 | 123.8 | 84.83 | -31.48% |

**Table S8. Dwell time summary statistics for a prototype version of CRC-AIM.** The CRC-AIM prototype was prior to recalibration. See **Table S2** for dwell time summary statistics for other models, including CISNET models and recalibrated CRC-AIM. IQR = interquartile range

| **Measure** | **Statistic** | **Prototype**  **CRC-AIM** |
| --- | --- | --- |
| **Adenoma dwell time, y** | Mean | 26.7 |
|  | Median | 25 |
|  | IQR | 17-35 |
| **Sojourn time (preclinical cancer dwell time), y** | Mean | 2.1 |
|  | Median | 1.6 |
|  | IQR | 1.0-2.7 |
| **Overall dwell time (full adenoma-carcinoma sequence), y** | Mean | 28.9 |
|  | Median | 27.2 |
|  | IQR | 18.9-37.3 |

**Table S9. Coding details for derived variables used for the SEER survival analysis.** Survival analysis was performed using survival years (where a month value of 0 was recoded as 0.5) versus age and sex factors, stratified by site and American Joint Committee on Cancer (AJCC) stage. Survival models were selected based on smallest Akaike information criterion (AICc). Analysis performed using JMP v13.0 (SAS Institute).

| **New Column Name** | **Description** | **Code** |
| --- | --- | --- |
| “Age at Dx Recode 5 Groups” | Re-classifies SEER*STAT “Age recode with <1 year olds” (age at Diagnosis, 17 groups) to 5 groups | Match( :Name( "Age recode with <1 year olds" ),  "01-04 years", "<20",  "10-14 years", "<20",  "15-19 years", "<20",  "20-24 years", "20-49",  "25-29 years", "20-49",  "30-34 years", "20-49",  "35-39 years", "20-49",  "40-44 years", "20-49",  "45-49 years", "20-49",  "50-54 years", "50-59",  "55-59 years", "50-59",  "60-64 years", "60-69",  "65-69 years", "60-69",  "70-74 years", "70-79",  "75-79 years", "70-79",  "80-84 years", ">=80",  "85+ years", ">=80",  Empty()  ) |
| “Site: C vs R” | Recodes SEER*STAT “Site Colon vs Rectum” to either Colon or Rectum | Match( :Name( "Site recode ICD-O-3/WHO 2008" ),  "Appendix", "Colon",  "Ascending Colon","Colon",  "Cecum", "Colon",  "Descending Colon", "Colon",  "Hepatic Flexure", "Colon",  "Large Intestine, NOS", "Colon",  "Rectosigmoid Junction","Rectum",  "Rectum", "Rectum",  "Sigmoid Colon", "Colon",  "Splenic Flexure", "Colon",  "Transverse Colon", "Colon",  Empty()  ) |
| “Exclude Based on AJCC” | Filter column to exclude subjects where AJCC is Unstaged | Match( :Name( "AJCC 5th Ed Schrag Code" ),  "AJCC 5th Ed Stage I", "NO",  "AJCC 5th Ed Stage II", "NO",  "AJCC 5th Ed Stage III", "NO",  "AJCC 5th Ed Stage IV", "NO",  "AJCC 5th Ed Unstaged", "YES",  Empty()  ) |
| “Exclude Age too Young” | Filter column to exclude subjects who are younger than the youngest CISNET Age to be at risk of colorectal cancer (<Age 20) | Match( :Age at Dx Recode 5 Groups,  "<20", "YES",  ">=80", "NO",  "20-49", "NO",  "50-59", "NO",  "60-69", "NO",  "70-79", "NO",  Empty()  ) |
| “ExcludeIs1.Survival.Months” | Filter column to exclude from analysis where survival values could not reliably be calculated and there could be 0 days of follow-up. Uses SEER*STAT survival months flag column.  If survival month dates were incomplete and there could be 0 days of follow-up, then data was excluded.  We included data where complete dates were available. | Match( :Survival months flag,  "Complete dates are available and there are 0 days of survival", "0",  "Complete dates are available and there are more than 0 days of survival", "0",  "Incomplete dates are available and there cannot be zero days of follow-up", "0",  "Incomplete dates are available and there could be zero days of follow-up", "1",  Empty()  ) |
| “Surv.Mo.0.as.05” | When sufficient documentation existed for a recorded 0-month survival, the 0-month survival was recoded as 0.5 month survival (so it is not censored in survival analysis).  Note: this can alter the type of parametric regression that is ultimately selected. | If( :ExcludeIs1.Survival.Months == "0" & :Survival months == 0, 0.5, If( :ExcludeIs1.Survival.Months == "0" & :Survival months != 0, :Survival months, Empty())) |
| “Surv.Year.MonthRecoded” | Calculates survival years using the “Surv.Mo.0.as.05” column | :Surv.Mo.0.as0.5 / 12 |
| “CensorEquals1” | If the individual is recorded as still be alive at the end of the follow-up or has died of other causes, then the individual is considered as a Censor candidate (pending not being excluded due to other factors listed above). | Match( :Name( "SEER cause-specific death classification" ),  "Alive or dead of other cause", 1,  "Dead (attributable to this cancer dx)", 0,  Empty()  ) |

**Table S10. Model selection details and fitted model diagnostics and parameter estimates for cause-specific survival for 1975-79 stage I colorectal cancer (CRC) in the colon.** The highlighted distribution was selected because it resulted in the smallest Akaike information criterion (AICc) and the parameter estimates correspond to that distribution. Parameter δ = 1/κ. The Wald confidence interval was used.

| **1975-79 Stage I CRC, Colon: Model Comparison** | | | | |
| --- | --- | --- | --- | --- |
| **Distribution** | **AICc** |  | | |
| Weibull | 6825.0 |  |  |  |
| Lognormal | 6864.6 |  |  |  |
| Exponential | 6895.8 |  |  |  |
| Frechet | 6936.9 |  |  |  |
| Loglogistic | 6829.9 |  |  |  |
| Observation Used | 3555 |  | | |
| Uncensored Values | 667 |  |  |  |
| Right Censored Values | 2888 |  |  |  |
| **Whole Model Test** | | | | |
| **ChiSquare** | **DF** | **Prob>Chisq** |  | |
| 134.9306 | 5 | <.0001 |  |  |
| **Parameter Estimates** | | | | |
| **Term** | **Estimate** | **Std Error** | **Lower 95%** | **Upper 95%** |
| Intercept | 4.787517 | 0.092399 | 4.606419 | 4.968616 |
| Sex[Female] | 0.858742 | 0.169591 | 0.526351 | 1.191134 |
| Age at Dx Recode 5 Groups[20-49] | 0.586693 | 0.117597 | 0.356207 | 0.817178 |
| Age at Dx Recode 5 Groups[50-59] | 0.135153 | 0.095296 | -0.051624 | 0.321931 |
| Age at Dx Recode 5 Groups[60-69] | -0.354845 | 0.093406 | -0.537918 | -0.171773 |
| Age at Dx Recode 5 Groups[70-79] | 0.200183 | 0.051827 | 0.098603 | 0.301763 |
| δ | 1.324690 | 0.046260 | 1.234021 | 1.415358 |
| **Wald Tests** | | | | |
| **Source** | **Nparm** | **DF** | **Wald ChiSquare** | **Prob>ChiSq** |
| Sex | 1 | 1 | 14.9188 | 0.0001 |
| Age at Dx Recode 5 Groups | 4 | 4 | 145.6113 | <.0001 |

**Table S11. Model selection details and fitted model diagnostics and parameter estimates for cause-specific survival for 1975-79 stage I colorectal cancer (CRC) in the rectum.** The highlighted distribution was selected because it resulted in the smallest Akaike information criterion (AICc) and the parameter estimates correspond to that distribution. The Wald confidence interval was used.

| **1975-79 Stage I CRC, Rectum: Model Comparison** | | | | |
| --- | --- | --- | --- | --- |
| **Distribution** | **AICc** |  | | |
| Weibull | 7812.9 |  |  |  |
| Lognormal | 7787.6 |  |  |  |
| Exponential | 7933.7 |  |  |  |
| Frechet | 7849.7 |  |  |  |
| Loglogistic | 7792.9 |  |  |  |
| Observation Used | 2951 |  | | |
| Uncensored Values | 870 |  |  |  |
| Right Censored Values | 2081 |  |  |  |
| **Whole Model Test** | | | | |
| **ChiSquare** | **DF** | **Prob>Chisq** |  | |
| 196.0059 | 5 | <.0001 |  |  |
| **Parameter Estimates** | | | | |
| **Term** | **Estimate** | **Std Error** | **Lower 95%** | **Upper 95%** |
| Intercept | 3.795728 | 0.078583 | 3.641709 | 3.949748 |
| Sex[Female] | 0.222715 | 0.053987 | 0.116902 | 0.328528 |
| Age at Dx Recode 5 Groups[20-49] | 0.911051 | 0.171131 | 0.575640 | 1.246462 |
| Age at Dx Recode 5 Groups[50-59] | 0.741344 | 0.111520 | 0.522769 | 0.959920 |
| Age at Dx Recode 5 Groups[60-69] | 0.171881 | 0.095995 | -0.016266 | 0.360028 |
| Age at Dx Recode 5 Groups[70-79] | -0.244835 | 0.101088 | -0.442965 | -0.046706 |
| δ | 2.273770 | 0.060525 | 2.155143 | 2.392396 |
| **Wald Tests** | | | | |
| **Source** | **Nparm** | **DF** | **Wald ChiSquare** | **Prob>ChiSq** |
| Sex | 1 | 1 | 17.0182 | <.0001 |
| Age at Dx Recode 5 Groups | 4 | 4 | 206.1739 | <.0001 |

**Table S12. Model selection details and fitted model diagnostics and parameter estimates for cause-specific survival for 1975-79 stage II colorectal cancer (CRC) in the colon.** The highlighted distribution was selected because it resulted in the smallest Akaike information criterion (AICc) and the parameter estimates correspond to that distribution. The Wald confidence interval was used.

| **1975-79 Stage II CRC, Colon: Model Comparison** | | | | |
| --- | --- | --- | --- | --- |
| **Distribution** | **AICc** |  | | |
| Weibull | 26345.42 |  |  |  |
| Lognormal | 26170.66 |  |  |  |
| Exponential | 27542.08 |  |  |  |
| Frechet | 26271.67 |  |  |  |
| Loglogistic | 26247.34 |  |  |  |
| Observation Used | 9215 |  | | |
| Uncensored Values | 3172 |  |  |  |
| Right Censored Values | 6043 |  |  |  |
| **Whole Model Test** | | | | |
| **ChiSquare** | **DF** | **Prob>Chisq** |  | |
| 210.33 | 5 | <.0001 |  |  |
| **Parameter Estimates** | | | | |
| **Term** | **Estimate** | **Std Error** | **Lower 95%** | **Upper 95%** |
| Intercept | 3.600079 | 0.046776 | 3.508399 | 3.691758 |
| Sex[Female] | 0.192775 | 0.033208 | 0.127689 | 0.257860 |
| Age at Dx Recode 5 Groups[20-49] | 0.913622 | 0.108666 | 0.700641 | 1.126604 |
| Age at Dx Recode 5 Groups[50-59] | 0.207249 | 0.078138 | 0.054102 | 0.360396 |
| Age at Dx Recode 5 Groups[60-69] | 0.026601 | 0.062580 | -0.096054 | 0.149255 |
| Age at Dx Recode 5 Groups[70-79] | -0.260595 | 0.059840 | -0.377880 | -0.143311 |
| δ | 2.610969 | 0.036533 | 2.539365 | 2.682572 |
| **Wald Tests** | | | | |
| **Source** | **Nparm** | **DF** | **Wald ChiSquare** | **Prob>ChiSq** |
| Sex | 1 | 1 | 33.6997 | <.0001 |
| Age at Dx Recode 5 Groups | 4 | 4 | 194.7110 | <.0001 |

**Table S13. Model selection details and fitted model diagnostics and parameter estimates for cause-specific survival for 1975-79 stage II colorectal cancer (CRC) in the rectum.** The highlighted distribution was selected because it resulted in the smallest Akaike information criterion (AICc) and the parameter estimates correspond to that distribution. The Wald confidence interval was used.

| **1975-79 Stage II CRC, Rectum: Model Comparison** | | | | |
| --- | --- | --- | --- | --- |
| **Distribution** | **AICc** |  | | |
| Weibull | 10796.0836 |  |  |  |
| Lognormal | 10639.6158 |  |  |  |
| Exponential | 11120.6735 |  |  |  |
| Frechet | 10682.5047 |  |  |  |
| Loglogistic | 10686.3145 |  |  |  |
| Observation Used | 3016 |  | | |
| Uncensored Values | 1432 |  |  |  |
| Right Censored Values | 1584 |  |  |  |
| **Whole Model Test** | | | | |
| **ChiSquare** | **DF** | **Prob>Chisq** |  | |
| 141.3549 | 5 | <.0001 |  |  |
| **Parameter Estimates** | | | | |
| **Term** | **Estimate** | **Std Error** | **Lower 95%** | **Upper 95%** |
| Intercept | 2.527056 | 0.054762 | 2.419724 | 2.634389 |
| Sex[Female] | 0.143037 | 0.042876 | 0.059002 | 0.227073 |
| Age at Dx Recode 5 Groups[20-49] | 0.538863 | 0.144432 | 0.255782 | 0.821944 |
| Age at Dx Recode 5 Groups[50-59] | 0.465769 | 0.089596 | 0.290165 | 0.641373 |
| Age at Dx Recode 5 Groups[60-69] | 0.229271 | 0.076962 | 0.078427 | 0.380115 |
| Age at Dx Recode 5 Groups[70-79] | -0.099496 | 0.079690 | -0.255685 | 0.056693 |
| δ | 2.042596 | 0.041566 | 1.961128 | 2.124063 |
| **Wald Tests** | | | | |
| **Source** | **Nparm** | **DF** | **Wald ChiSquare** | **Prob>ChiSq** |
| Sex | 1 | 1 | 11.1294 | 0.0008 |
| Age at Dx Recode 5 Groups | 4 | 4 | 145.4905 | <.0001 |

**Table S14. Model selection details and fitted model diagnostics and parameter estimates for cause-specific survival for 1975-79 stage III colorectal cancer (CRC) in the colon.** The highlighted distribution was selected because it resulted in the smallest Akaike information criterion (AICc) and the parameter estimates correspond to that distribution. The Wald confidence interval was used.

| **1975-79 Stage III CRC, Colon: Model Comparison** | | | | |
| --- | --- | --- | --- | --- |
| **Distribution** | **AICc** |  | | |
| Weibull | 23746.7629 |  |  |  |
| Lognormal | 23031.5681 |  |  |  |
| Exponential | 25518.8877 |  |  |  |
| Frechet | 22867.6911 |  |  |  |
| Loglogistic | 23169.7505 |  |  |  |
| Observation Used | 6205 |  | | |
| Uncensored Values | 3667 |  |  |  |
| Right Censored Values | 2538 |  |  |  |
| **Whole Model Test** | | | | |
| **ChiSquare** | **DF** | **Prob>Chisq** |  | |
| 139.7007 | 4 | <.0001 |  |  |
| **Parameter Estimates** | | | | |
| **Term** | **Estimate** | **Std Error** | **Lower 95%** | **Upper 95%** |
| Intercept | 0.972896 | 0.034369 | 0.905534 | 1.040258 |
| Age at Dx Recode 5 Groups[20-49] | 0.382867 | 0.083996 | 0.218238 | 0.547496 |
| Age at Dx Recode 5 Groups[50-59] | 0.259110 | 0.060790 | 0.139964 | 0.378256 |
| Age at Dx Recode 5 Groups[60-69] | 0.178915 | 0.052158 | 0.076687 | 0.281143 |
| Age at Dx Recode 5 Groups[70-79] | -0.174654 | 0.051553 | -0.275695 | -0.073612 |
| σ | 2.200354 | 0.026582 | 2.148254 | 2.252454 |
| **Wald Tests** | | | | |
| **Source** | **Nparm** | **DF** | **Wald ChiSquare** | **Prob>ChiSq** |
| Age at Dx Recode 5 Groups | 4 | 4 | 139.0287 | <.0001 |

**Table S15. Preliminary model parameter estimates for the cause-specific survival for 1975-79 stage III colorectal cancer (CRC) in the colon.** Parameter estimates correspond to the highlighted distribution in **Table S14**, demonstrating that sex is not a significant covariate. The Wald confidence interval was used.

| **1975-79 Stage III CRC, Colon: Preliminary Model Comparison** | | | | |
| --- | --- | --- | --- | --- |
| **Parameter Estimates** | | | | |
| **Term** | **Estimate** | **Std Error** | **Lower 95%** | **Upper 95%** |
| Intercept | 0.973078 | 0.034460 | 0.905538 | 1.040618 |
| Sex[Female] | -0.002081 | 0.028599 | -0.058135 | 0.053972 |
| Age at Dx Recode 5 Groups[20-49] | 0.382792 | 0.084002 | 0.218150 | 0.547433 |
| Age at Dx Recode 5 Groups[50-59] | 0.258967 | 0.060821 | 0.139760 | 0.378175 |
| Age at Dx Recode 5 Groups[60-69] | 0.178682 | 0.052256 | 0.076263 | 0.281102 |
| Age at Dx Recode 5 Groups[70-79] | -0.174549 | 0.051573 | -0.275630 | -0.073468 |
| σ | 2.200347 | 0.026582 | 2.148247 | 2.252448 |
| **Wald Tests** | | | | |
| **Source** | **Nparm** | **DF** | **Wald ChiSquare** | **Prob>ChiSq** |
| Sex | 1 | 1 | 0.0053 | 0.9420 |
| Age at Dx Recode 5 Groups | 4 | 4 | 137.6711 | <.0001 |

**Table S16. Model selection details and fitted model diagnostics and parameter estimates for cause-specific survival for 1975-79 stage III colorectal cancer (CRC) in the rectum.** The highlighted distribution was selected because it resulted in the smallest Akaike information criterion (AICc) and the parameter estimates correspond to that distribution. The Wald confidence interval was used.

| **1975-79 Stage III CRC, Rectum: Model Comparison** | | | | |
| --- | --- | --- | --- | --- |
| **Distribution** | **AICc** |  | | |
| Weibull | 11003.1542 |  |  |  |
| Lognormal | 10616.6864 |  |  |  |
| Exponential | 11431.3049 |  |  |  |
| Frechet | 10616.7219 |  |  |  |
| Loglogistic | 10621.7841 |  |  |  |
| Observation Used | 2640 |  | | |
| Uncensored Values | 1764 |  |  |  |
| Right Censored Values | 876 |  |  |  |
| **Whole Model Test** | | | | |
| **ChiSquare** | **DF** | **Prob>Chisq** |  | |
| 64.0900 | 5 | <.0001 |  |  |
| **Parameter Estimates** | | | | |
| **Term** | **Estimate** | **Std Error** | **Lower 95%** | **Upper 95%** |
| Intercept | 1.549157 | 0.041109 | 1.468586 | 1.629729 |
| Sex[Female] | 0.113776 | 0.035208 | 0.044770 | 0.182783 |
| Age at Dx Recode 5 Groups[20-49] | 0.409712 | 0.101801 | 0.210185 | 0.609238 |
| Age at Dx Recode 5 Groups[50-59] | 0.280006 | 0.068825 | 0.145111 | 0.414901 |
| Age at Dx Recode 5 Groups[60-69] | 0.031083 | 0.061877 | -0.090193 | 0.152359 |
| Age at Dx Recode 5 Groups[70-79] | -0.171752 | 0.066091 | -0.301289 | -0.042216 |
| σ | 1.675403 | 0.029953 | 1.616696 | 1.734110 |
| **Wald Tests** | | | | |
| **Source** | **Nparm** | **DF** | **Wald ChiSquare** | **Prob>ChiSq** |
| Sex | 1 | 1 | 10.4428 | 0.0012 |
| Age at Dx Recode 5 Groups | 4 | 4 | 57.8493 | <.0001 |

**Table S17. Model selection details and fitted model diagnostics and parameter estimates for cause-specific survival for 1975-79 stage IV colorectal cancer (CRC) in the colon.** The highlighted distribution was selected because it resulted in the smallest Akaike information criterion (AICc) and the parameter estimates correspond to that distribution. DNC = did not converge. The Wald confidence interval was used.

| **1975-79 Stage IV CRC, Colon: Model Comparison** | | | | |
| --- | --- | --- | --- | --- |
| **Distribution** | **AICc** |  | | |
| Weibull | 14411.1196 |  |  |  |
| Lognormal | 12728.1446 |  |  |  |
| Exponential | DNC |  |  |  |
| Frechet | 12705.7556 |  |  |  |
| Loglogistic | 12712.6132 |  |  |  |
| Observation Used | 6853 |  | | |
| Uncensored Values | 6209 |  |  |  |
| Right Censored Values | 644 |  |  |  |
| **Whole Model Test** | | | | |
| **ChiSquare** | **DF** | **Prob>Chisq** |  | |
| 268.2056 | 5 | <.0001 |  |  |
| **Parameter Estimates** | | | | |
| **Term** | **Estimate** | **Std Error** | **Lower 95%** | **Upper 95%** |
| Intercept | -1.361909 | 0.019978 | -1.401066 | -1.322753 |
| Sex[Female] | 0.036882 | 0.016846 | 0.003865 | 0.069900 |
| Age at Dx Recode 5 Groups[20-49] | 0.259513 | 0.052021 | 0.157554 | 0.361473 |
| Age at Dx Recode 5 Groups[50-59] | 0.312021 | 0.036765 | 0.239963 | 0.384078 |
| Age at Dx Recode 5 Groups[60-69] | 0.068881 | 0.030781 | 0.008550 | 0.129211 |
| Age at Dx Recode 5 Groups[70-79] | -0.111060 | 0.030466 | -0.170773 | -0.051347 |
| σ | 1.372644 | 0.013103 | 1.346962 | 1.398326 |
| **Wald Tests** | | | | |
| **Source** | **Nparm** | **DF** | **Wald ChiSquare** | **Prob>ChiSq** |
| Sex | 1 | 1 | 4.7934 | 0.0286 |
| Age at Dx Recode 5 Groups | 4 | 4 | 264.7558 | <.0001 |

**Table S18. Model selection details and fitted model diagnostics and parameter estimates for cause-specific survival for 1975-79 stage IV colorectal cancer (CRC) in the rectum.** The highlighted distribution was selected because it resulted in the smallest Akaike information criterion (AICc) and the parameter estimates correspond to that distribution. DNC = did not converge. The Wald confidence interval was used.

| **1975-79 Stage IV CRC, Rectum: Model Comparison** | | | | |
| --- | --- | --- | --- | --- |
| **Distribution** | **AICc** |  | | |
| Weibull | 5536.4062 |  |  |  |
| Lognormal | 5070.5617 |  |  |  |
| Exponential | DNC |  |  |  |
| Frechet | 5254.9737 |  |  |  |
| Loglogistic | 5065.0192 |  |  |  |
| Observation Used | 2408 |  | | |
| Uncensored Values | 2212 |  |  |  |
| Right Censored Values | 196 |  |  |  |
| **Whole Model Test** | | | | |
| **ChiSquare** | **DF** | **Prob>Chisq** |  | |
| 143.4434 | 4 | <.0001 |  |  |
| **Parameter Estimates** | | | | |
| **Term** | **Estimate** | **Std Error** | **Lower 95%** | **Upper 95%** |
| Intercept | -0.509357 | 0.031974 | -0.572025 | -0.446688 |
| Age at Dx Recode 5 Groups[20-49] | 0.415409 | 0.084990 | 0.248831 | 0.581987 |
| Age at Dx Recode 5 Groups[50-59] | 0.316781 | 0.056604 | 0.205838 | 0.427723 |
| Age at Dx Recode 5 Groups[60-69] | 0.125683 | 0.050379 | 0.026942 | 0.224424 |
| Age at Dx Recode 5 Groups[70-79] | -0.096889 | 0.053525 | -0.201797 | 0.008018 |
| σ | 0.785458 | 0.013837 | 0.758339 | 0.812577 |
| **Wald Tests** | | | | |
| **Source** | **Nparm** | **DF** | **Wald ChiSquare** | **Prob>ChiSq** |
| Age at Dx Recode 5 Groups | 4 | 4 | 149.8985 | <.0001 |

**Table S19. Preliminary model parameter estimates for the cause-specific survival for 1975-79 stage IV colorectal cancer (CRC) in the rectum.** Parameter estimates correspond to the highlighted distribution in **Table S18**, demonstrating that sex is not a significant covariate. The Wald confidence interval was used.

| **1975-79 Stage IV CRC, Rectum: Preliminary Model Comparison** | | | | |
| --- | --- | --- | --- | --- |
| **Parameter Estimates** | | | | |
| **Term** | **Estimate** | **Std Error** | **Lower 95%** | **Upper 95%** |
| Intercept | -0.511735 | 0.032056 | -0.574563 | -0.448906 |
| Sex[Female] | -0.027858 | 0.028592 | -0.083898 | 0.028181 |
| Age at Dx Recode 5 Groups[20-49] | 0.413433 | 0.084969 | 0.246896 | 0.579970 |
| Age at Dx Recode 5 Groups[50-59] | 0.312648 | 0.056746 | 0.201428 | 0.423869 |
| Age at Dx Recode 5 Groups[60-69] | 0.122855 | 0.050451 | 0.023973 | 0.221737 |
| Age at Dx Recode 5 Groups[70-79] | -0.095562 | 0.053518 | -0.200456 | 0.009332 |
| σ | 0.785255 | 0.013834 | 0.758140 | 0.812370 |
| **Wald Tests** | | | | |
| **Source** | **Nparm** | **DF** | **Wald ChiSquare** | **Prob>ChiSq** |
| Sex | 1 | 1 | 0.9493 | 0.3299 |
| Age at Dx Recode 5 Groups | 4 | 4 | 144.2795 | <.0001 |

**Table S20. Model selection details and fitted model diagnostics and parameter estimates for cause-specific survival for 2000-03 stage I colorectal cancer (CRC) in the colon.** The highlighted distribution was selected because it resulted in the smallest Akaike information criterion (AICc) and the parameter estimates correspond to that distribution. Parameter δ = 1/κ. The Wald confidence interval was used.

| **2000-03 Stage I CRC, Colon: Model Comparison** | | | | |
| --- | --- | --- | --- | --- |
| **Distribution** | **AICc** |  | | |
| Weibull | 6712.5586 |  |  |  |
| Lognormal | 6732.2328 |  |  |  |
| Exponential | 6820.2702 |  |  |  |
| Frechet | 6778.3830 |  |  |  |
| Loglogistic | 6714.0577 |  |  |  |
| Observation Used | 6054 |  | | |
| Uncensored Values | 645 |  |  |  |
| Right Censored Values | 5409 |  |  |  |
| **Whole Model Test** | | | | |
| **ChiSquare** | **DF** | **Prob>Chisq** |  | |
| 241.2994 | 5 | <.0001 |  |  |
| **Parameter Estimates** | | | | |
| **Term** | **Estimate** | **Std Error** | **Lower 95%** | **Upper 95%** |
| Intercept | 5.879682 | 0.163628 | 5.558976 | 6.200387 |
| Sex[Female] | 0.132082 | 0.057253 | 0.019869 | 0.244295 |
| Age at Dx Recode 5 Groups[20-49] | 1.609704 | 0.352986 | 0.917864 | 2.301544 |
| Age at Dx Recode 5 Groups[50-59] | 0.649916 | 0.180608 | 0.295931 | 1.003902 |
| Age at Dx Recode 5 Groups[60-69] | -0.011486 | 0.141261 | -0.288353 | 0.265381 |
| Age at Dx Recode 5 Groups[70-79] | -0.486255 | 0.128291 | -0.737700 | -0.234809 |
| δ | 1.430413 | 0.052380 | 1.327749 | 1.533077 |
| **Wald Tests** | | | | |
| **Source** | **Nparm** | **DF** | **Wald ChiSquare** | **Prob>ChiSq** |
| Sex | 1 | 1 | 5.3222 | 0.0211 |
| Age at Dx Recode 5 Groups | 4 | 4 | 204.3100 | <.0001 |

**Table S21. Model selection details and fitted model diagnostics and parameter estimates for cause-specific survival for 2000-03 stage I colorectal cancer (CRC) in the rectum.** The highlighted distribution was selected because it resulted in the smallest Akaike information criterion (AICc) and the parameter estimates correspond to that distribution. The Wald confidence interval was used.

| **2000-03 Stage I CRC, Rectum: Model Comparison** | | | | |
| --- | --- | --- | --- | --- |
| **Distribution** | **AICc** |  | | |
| Weibull | 4427.7791 |  |  |  |
| Lognormal | 4431.6023 |  |  |  |
| Exponential | 4439.5690 |  |  |  |
| Frechet | 4462.4581 |  |  |  |
| Loglogistic | 4425.4890 |  |  |  |
| Observation Used | 2827 |  | | |
| Uncensored Values | 439 |  |  |  |
| Right Censored Values | 2388 |  |  |  |
| **Whole Model Test** | | | | |
| **ChiSquare** | **DF** | **Prob>Chisq** |  | |
| 241.2994 | 5 | <.0001 |  |  |
| **Parameter Estimates** | | | | |
| **Term** | **Estimate** | **Std Error** | **Lower 95%** | **Upper 95%** |
| Intercept | 4.247524 | 0.106285 | 4.039210 | 4.455839 |
| Age at Dx Recode 5 Groups[20-49] | 0.470342 | 0.163954 | 0.148998 | 0.791685 |
| Age at Dx Recode 5 Groups[50-59] | 0.703278 | 0.131565 | 0.445416 | 0.961140 |
| Age at Dx Recode 5 Groups[60-69] | 0.159296 | 0.110116 | -0.056527 | 0.375118 |
| Age at Dx Recode 5 Groups[70-79] | -0.256080 | 0.104809 | -0.461501 | -0.050659 |
| δ | 1.107932 | 0.048396 | 1.013078 | 1.202787 |
| **Wald Tests** | | | | |
| **Source** | **Nparm** | **DF** | **Wald ChiSquare** | **Prob>ChiSq** |
| Age at Dx Recode 5 Groups | 4 | 4 | 96.8821 | <.0001 |

**Table S22. Preliminary model parameter estimates for the cause-specific survival for 2000-03 stage I colorectal cancer (CRC) in the rectum.** Parameter estimates correspond to the highlighted distribution in **Table S20**, demonstrating that sex is not a significant covariate. The Wald confidence interval was used.

| **2000-03 Stage I CRC, Rectum: Preliminary Model Comparison** | | | | |
| --- | --- | --- | --- | --- |
| **Parameter Estimates** | | | | |
| **Term** | **Estimate** | **Std Error** | **Lower 95%** | **Upper 95%** |
| Intercept | 4.258682 | 0.107017 | 4.048934 | 4.468431 |
| Sex[Female] | 0.087242 | 0.059925 | -0.030209 | 0.204693 |
| Age at Dx Recode 5 Groups[20-49] | 0.476808 | 0.163836 | 0.155696 | 0.797920 |
| Age at Dx Recode 5 Groups[50-59] | 0.713406 | 0.131665 | 0.455347 | 0.971464 |
| Age at Dx Recode 5 Groups[60-69] | 0.171302 | 0.110311 | -0.044904 | 0.387507 |
| Age at Dx Recode 5 Groups[70-79] | -0.260059 | 0.104704 | -0.465276 | -0.054843 |
| σ | 1.106348 | 0.048312 | 1.011658 | 1.201038 |
| **Wald Tests** | | | | |
| **Source** | **Nparm** | **DF** | **Wald ChiSquare** | **Prob>ChiSq** |
| Sex | 1 | 1 | 2.1195 | 0.1454 |
| Age at Dx Recode 5 Groups | 4 | 4 | 98.7242 | <.0001 |

**Table S23. Model selection details and fitted model diagnostics and parameter estimates for cause-specific survival for 2000-03 stage II colorectal cancer (CRC) in the colon.** The highlighted distribution was selected because it resulted in the smallest Akaike information criterion (AICc) and the parameter estimates correspond to that distribution. The Wald confidence interval was used.

| **2000-03 Stage II CRC, Colon: Model Comparison** | | | | |
| --- | --- | --- | --- | --- |
| **Distribution** | **AICc** |  | | |
| Weibull | 16392.3346 |  |  |  |
| Lognormal | 16356.1742 |  |  |  |
| Exponential | 16909.8138 |  |  |  |
| Frechet | 16423.1679 |  |  |  |
| Loglogistic | 16372.8197 |  |  |  |
| Observation Used | 8258 |  | | |
| Uncensored Values | 1866 |  |  |  |
| Right Censored Values | 6392 |  |  |  |
| **Whole Model Test** | | | | |
| **ChiSquare** | **DF** | **Prob>Chisq** |  | |
| 204.1411 | 5 | <.0001 |  |  |
| **Parameter Estimates** | | | | |
| **Term** | **Estimate** | **Std Error** | **Lower 95%** | **Upper 95%** |
| Intercept | 4.501293 | 0.072495 | 4.359205 | 4.643381 |
| Sex[Female] | 0.182476 | 0.042847 | 0.098496 | 0.266455 |
| Age at Dx Recode 5 Groups[20-49] | 0.619266 | 0.136899 | 0.350949 | 0.887583 |
| Age at Dx Recode 5 Groups[50-59] | 0.377763 | 0.106751 | 0.168534 | 0.586992 |
| Age at Dx Recode 5 Groups[60-69] | 0.324346 | 0.089388 | 0.149149 | 0.499542 |
| Age at Dx Recode 5 Groups[70-79] | -0.222634 | 0.076338 | -0.372255 | -0.073013 |
| δ | 2.831562 | 0.052886 | 2.727907 | 2.935217 |
| **Wald Tests** | | | | |
| **Source** | **Nparm** | **DF** | **Wald ChiSquare** | **Prob>ChiSq** |
| Sex | 1 | 1 | 18.1369 | <.0001 |
| Age at Dx Recode 5 Groups | 4 | 4 | 206.2314 | <.0001 |

**Table S24. Model selection details and fitted model diagnostics and parameter estimates for cause-specific survival for 2000-03 stage II colorectal cancer (CRC) in the rectum.** The highlighted distribution was selected because it resulted in the smallest Akaike information criterion (AICc) and the parameter estimates correspond to that distribution. The Wald confidence interval was used.

| **2000-03 Stage II CRC, Rectum: Model Comparison** | | | | |
| --- | --- | --- | --- | --- |
| **Distribution** | **AICc** |  | | |
| Weibull | 5542.9467 |  |  |  |
| Lognormal | 5520.6179 |  |  |  |
| Exponential | 5607.6837 |  |  |  |
| Frechet | 5566.6658 |  |  |  |
| Loglogistic | 5524.4337 |  |  |  |
| Observation Used | 2183 |  | | |
| Uncensored Values | 663 |  |  |  |
| Right Censored Values | 1520 |  |  |  |
| **Whole Model Test** | | | | |
| **ChiSquare** | **DF** | **Prob>Chisq** |  | |
| 117.9298 | 4 | <.0001 |  |  |
| **Parameter Estimates** | | | | |
| **Term** | **Estimate** | **Std Error** | **Lower 95%** | **Upper 95%** |
| Intercept | 3.467962 | 0.084494 | 3.302357 | 3.633568 |
| Age at Dx Recode 5 Groups[20-49] | 0.960835 | 0.164340 | 0.638735 | 1.282935 |
| Age at Dx Recode 5 Groups[50-59] | 0.497604 | 0.122020 | 0.258449 | 0.736758 |
| Age at Dx Recode 5 Groups[60-69] | 0.114634 | 0.111704 | -0.104301 | 0.333570 |
| Age at Dx Recode 5 Groups[70-79] | -0.391419 | 0.105896 | -0.598971 | -0.183867 |
| δ | 2.187381 | 0.067674 | 2.054742 | 2.320019 |
| **Wald Tests** | | | | |
| **Source** | **Nparm** | **DF** | **Wald ChiSquare** | **Prob>ChiSq** |
| Age at Dx Recode 5 Groups | 4 | 4 | 119.1489 | <.0001 |

**Table S25. Preliminary model parameter estimates for the cause-specific survival for 2000-03 stage II colorectal cancer (CRC) in the rectum.** Parameter estimates correspond to the highlighted distribution in **Table S24**, demonstrating that sex is not a significant covariate. The Wald confidence interval was used.

| **2000-03 Stage II CRC, Rectum: Preliminary Model Comparison** | | | | |
| --- | --- | --- | --- | --- |
| **Parameter Estimates** | | | | |
| **Term** | **Estimate** | **Std Error** | **Lower 95%** | **Upper 95%** |
| Intercept | 3.467825 | 0.084915 | 3.301394 | 3.634256 |
| Sex[Female] | -0.000969 | 0.059721 | -0.118020 | 0.116081 |
| Age at Dx Recode 5 Groups[20-49] | 0.960789 | 0.164363 | 0.638643 | 1.282934 |
| Age at Dx Recode 5 Groups[50-59] | 0.497535 | 0.122094 | 0.258236 | 0.736834 |
| Age at Dx Recode 5 Groups[60-69] | 0.114536 | 0.111868 | -0.104721 | 0.333793 |
| Age at Dx Recode 5 Groups[70-79] | -0.391428 | 0.105897 | -0.598983 | -0.183874 |
| σ | 2.187376 | 0.067674 | 2.054737 | 2.320016 |
| **Wald Tests** | | | | |
| **Source** | **Nparm** | **DF** | **Wald ChiSquare** | **Prob>ChiSq** |
| Sex | 1 | 1 | 0.0003 | 0.9871 |
| Age at Dx Recode 5 Groups | 4 | 4 | 118.0013 | <.0001 |

**Table S26. Model selection details and fitted model diagnostics and parameter estimates for cause-specific survival for 2000-03 stage III colorectal cancer (CRC) in the colon.** The highlighted distribution was selected because it resulted in the smallest Akaike information criterion (AICc) and the parameter estimates correspond to that distribution. The Wald confidence interval was used.

| **2000-03 Stage III CRC, Colon: Model Comparison** | | | | |
| --- | --- | --- | --- | --- |
| **Distribution** | **AICc** |  | | |
| Weibull | 23078.6953 |  |  |  |
| Lognormal | 22728.0115 |  |  |  |
| Exponential | 23978.1744 |  |  |  |
| Frechet | 22747.3516 |  |  |  |
| Loglogistic | 22845.1864 |  |  |  |
| Observation Used | 7326 |  | | |
| Uncensored Values | 3284 |  |  |  |
| Right Censored Values | 4042 |  |  |  |
| **Whole Model Test** | | | | |
| **ChiSquare** | **DF** | **Prob>Chisq** |  | |
| 344.5836 | 4 | <.0001 |  |  |
| **Parameter Estimates** | | | | |
| **Term** | **Estimate** | **Std Error** | **Lower 95%** | **Upper 95%** |
| Intercept | 2.536067 | 0.036069 | 2.465373 | 2.606762 |
| Age at Dx Recode 5 Groups[20-49] | 0.541576 | 0.081362 | 0.382109 | 0.701043 |
| Age at Dx Recode 5 Groups[50-59] | 0.426254 | 0.066332 | 0.296246 | 0.556262 |
| Age at Dx Recode 5 Groups[60-69] | 0.206712 | 0.058163 | 0.092714 | 0.320710 |
| Age at Dx Recode 5 Groups[70-79] | -0.146281 | 0.053108 | -0.250372 | -0.042191 |
| σ | 2.204045 | 0.030060 | 2.145128 | 2.262963 |
| **Wald Tests** | | | | |
| **Source** | **Nparm** | **DF** | **Wald ChiSquare** | **Prob>ChiSq** |
| Age at Dx Recode 5 Groups | 4 | 4 | 359.3299 | <.0001 |

**Table S27. Preliminary model parameter estimates for the cause-specific survival for 2000-03 stage III colorectal cancer (CRC) in the colon.** Parameter estimates correspond to the highlighted distribution in **Table S26**, demonstrating that sex is not a significant covariate. The Wald confidence interval was used.

| **2000-03 Stage III CRC, Colon: Preliminary Model Comparison** | | | | |
| --- | --- | --- | --- | --- |
| **Parameter Estimates** | | | | |
| **Term** | **Estimate** | **Std Error** | **Lower 95%** | **Upper 95%** |
| Intercept | 2.534884 | 0.036061 | 2.464206 | 2.605561 |
| Sex[Female] | 0.057506 | 0.029917 | -0.001131 | 0.116142 |
| Age at Dx Recode 5 Groups[20-49] | 0.547783 | 0.081409 | 0.388224 | 0.707343 |
| Age at Dx Recode 5 Groups[50-59] | 0.434101 | 0.066450 | 0.303862 | 0.564341 |
| Age at Dx Recode 5 Groups[60-69] | 0.213685 | 0.058264 | 0.099489 | 0.327881 |
| Age at Dx Recode 5 Groups[70-79] | -0.149754 | 0.053122 | -0.253871 | -0.045638 |
| σ | 2.203402 | 0.030051 | 2.144504 | 2.262300 |
| **Wald Tests** | | | | |
| **Source** | **Nparm** | **DF** | **Wald ChiSquare** | **Prob>ChiSq** |
| Sex | 1 | 1 | 3.6947 | 0.0546 |
| Age at Dx Recode 5 Groups | 4 | 4 | 361.1771 | <.0001 |

**Table S28. Model selection details and fitted model diagnostics and parameter estimates for cause-specific survival for 2000-03 stage III colorectal cancer (CRC) in the rectum.** The highlighted distribution was selected because it resulted in the smallest Akaike information criterion (AICc) and the parameter estimates correspond to that distribution. The Wald confidence interval was used.

| **2000-03 Stage III CRC, Rectum: Model Comparison** | | | | |
| --- | --- | --- | --- | --- |
| **Distribution** | **AICc** |  | | |
| Weibull | 9203.2670 |  |  |  |
| Lognormal | 9061.5251 |  |  |  |
| Exponential | 9279.4864 |  |  |  |
| Frechet | 9086.8547 |  |  |  |
| Loglogistic | 9103.2215 |  |  |  |
| Observation Used | 2685 |  | | |
| Uncensored Values | 1248 |  |  |  |
| Right Censored Values | 1437 |  |  |  |
| **Whole Model Test** | | | | |
| **ChiSquare** | **DF** | **Prob>Chisq** |  | |
| 164.9559 | 4 | <.0001 |  |  |
| **Parameter Estimates** | | | | |
| **Term** | **Estimate** | **Std Error** | **Lower 95%** | **Upper 95%** |
| Intercept | 2.355839 | 0.043877 | 2.269841 | 2.441836 |
| Age at Dx Recode 5 Groups[20-49] | 0.470034 | 0.083975 | 0.305445 | 0.634622 |
| Age at Dx Recode 5 Groups[50-59] | 0.389915 | 0.073731 | 0.245406 | 0.534424 |
| Age at Dx Recode 5 Groups[60-69] | 0.303914 | 0.073070 | 0.160700 | 0.447128 |
| Age at Dx Recode 5 Groups[70-79] | -0.107169 | 0.072685 | -0.249628 | 0.035290 |
| σ | 1.751901 | 0.038754 | 1.675944 | 1.827857 |
| **Wald Tests** | | | | |
| **Source** | **Nparm** | **DF** | **Wald ChiSquare** | **Prob>ChiSq** |
| Age at Dx Recode 5 Groups | 4 | 4 | 174.3351 | <.0001 |

**Table S29. Preliminary model parameter estimates for the cause-specific survival for 2000-03 stage III colorectal cancer (CRC) in the rectum.** Parameter estimates correspond to the highlighted distribution in **Table S28**, demonstrating that sex is not a significant covariate. The Wald confidence interval was used.

| **2000-03 Stage III CRC, Rectum: Preliminary Model Comparison** | | | | |
| --- | --- | --- | --- | --- |
| **Parameter Estimates** | | | | |
| **Term** | **Estimate** | **Std Error** | **Lower 95%** | **Upper 95%** |
| Intercept | 2.360388 | 0.044059 | 2.274034 | 2.446742 |
| Sex[Female] | 0.048828 | 0.038920 | -0.027454 | 0.125110 |
| Age at Dx Recode 5 Groups[20-49] | 0.472463 | 0.083950 | 0.307924 | 0.637002 |
| Age at Dx Recode 5 Groups[50-59] | 0.395694 | 0.073836 | 0.250978 | 0.540410 |
| Age at Dx Recode 5 Groups[60-69] | 0.309335 | 0.073151 | 0.165962 | 0.452708 |
| Age at Dx Recode 5 Groups[70-79] | -0.108186 | 0.072639 | -0.250555 | 0.034183 |
| σ | 1.750808 | 0.038728 | 1.674903 | 1.826713 |
| **Wald Tests** | | | | |
| **Source** | **Nparm** | **DF** | **Wald ChiSquare** | **Prob>ChiSq** |
| Sex | 1 | 1 | 1.5740 | 0.2096 |
| Age at Dx Recode 5 Groups | 4 | 4 | 175.9034 | <.0001 |

**Table S30. Model selection details and fitted model diagnostics and parameter estimates for cause-specific survival for 2000-03 stage IV colorectal cancer (CRC) in the colon.** The highlighted distribution was selected because it resulted in the smallest Akaike information criterion (AICc) and the parameter estimates correspond to that distribution. The Wald confidence interval was used.

| **2000-03 Stage IV CRC, Colon: Model Comparison** | | | | |
| --- | --- | --- | --- | --- |
| **Distribution** | **AICc** |  | | |
| Weibull | 12638.3198 |  |  |  |
| Lognormal | 11934.9884 |  |  |  |
| Exponential | 14039.0074 |  |  |  |
| Frechet | 12219.7368 |  |  |  |
| Loglogistic | 12048.4684 |  |  |  |
| Observation Used | 5175 |  | | |
| Uncensored Values | 4668 |  |  |  |
| Right Censored Values | 507 |  |  |  |
| **Whole Model Test** | | | | |
| **ChiSquare** | **DF** | **Prob>Chisq** |  | |
| 403.4377 | 4 | <.0001 |  |  |
| **Parameter Estimates** | | | | |
| **Term** | **Estimate** | **Std Error** | **Lower 95%** | **Upper 95%** |
| Intercept | -0.376556 | 0.022984 | -0.421604 | -0.331508 |
| Age at Dx Recode 5 Groups[20-49] | 0.653115 | 0.055073 | 0.545174 | 0.761055 |
| Age at Dx Recode 5 Groups[50-59] | 0.319011 | 0.046280 | 0.228303 | 0.409719 |
| Age at Dx Recode 5 Groups[60-69] | 0.086081 | 0.042726 | 0.002340 | 0.169823 |
| Age at Dx Recode 5 Groups[70-79] | -0.292951 | 0.039731 | -0.370821 | -0.215080 |
| σ | 1.558274 | 0.016318 | 1.526292 | 1.590256 |
| **Wald Tests** | | | | |
| **Source** | **Nparm** | **DF** | **Wald ChiSquare** | **Prob>ChiSq** |
| Age at Dx Recode 5 Groups | 4 | 4 | 422.7788 | <.0001 |

**Table S31. Preliminary model parameter estimates for the cause-specific survival for 2000-03 stage IV colorectal cancer (CRC) in the colon.** Parameter estimates correspond to the highlighted distribution in **Table S30**, demonstrating that sex is not a significant covariate. The Wald confidence interval was used.

| **2000-03 Stage IV CRC, Colon: Preliminary Model Comparison** | | | | |
| --- | --- | --- | --- | --- |
| **Parameter Estimates** | | | | |
| **Term** | **Estimate** | **Std Error** | **Lower 95%** | **Upper 95%** |
| Intercept | -0.376659 | 0.022984 | -0.421708 | -0.331610 |
| Sex[Female] | 0.010891 | 0.022191 | -0.032603 | 0.054385 |
| Age at Dx Recode 5 Groups[20-49] | 0.653505 | 0.055077 | 0.545556 | 0.761454 |
| Age at Dx Recode 5 Groups[50-59] | 0.320078 | 0.046330 | 0.229272 | 0.410883 |
| Age at Dx Recode 5 Groups[60-69] | 0.087155 | 0.042781 | 0.003305 | 0.171004 |
| Age at Dx Recode 5 Groups[70-79] | -0.292947 | 0.039730 | -0.370815 | -0.215079 |
| σ | 1.558236 | 0.016317 | 1.526255 | 1.590218 |
| **Wald Tests** | | | | |
| **Source** | **Nparm** | **DF** | **Wald ChiSquare** | **Prob>ChiSq** |
| Sex | 1 | 1 | 0.2409 | 0.6236 |
| Age at Dx Recode 5 Groups | 4 | 4 | 420.4701 | <.0001 |

**Table S32. Model selection details and fitted model diagnostics and parameter estimates for cause-specific survival for 2000-03 stage IV colorectal cancer (CRC) in the rectum.** The highlighted distribution was selected because it resulted in the smallest Akaike information criterion (AICc) and the parameter estimates correspond to that distribution. The Wald confidence interval was used.

| **2000-03 Stage IV CRC, Rectum: Model Comparison** | | | | |
| --- | --- | --- | --- | --- |
| **Distribution** | **AICc** |  | | |
| Weibull | 4760.9838 |  |  |  |
| Lognormal | 4604.2003 |  |  |  |
| Exponential | 4958.6266 |  |  |  |
| Frechet | 4801.3275 |  |  |  |
| Loglogistic | 4614.0333 |  |  |  |
| Observation Used | 1680 |  | | |
| Uncensored Values | 1519 |  |  |  |
| Right Censored Values | 161 |  |  |  |
| **Whole Model Test** | | | | |
| **ChiSquare** | **DF** | **Prob>Chisq** |  | |
| 188.1231 | 4 | <.0001 |  |  |
| **Parameter Estimates** | | | | |
| **Term** | **Estimate** | **Std Error** | **Lower 95%** | **Upper 95%** |
| Intercept | -0.170694 | 0.036309 | -0.241859 | -0.099529 |
| Age at Dx Recode 5 Groups[20-49] | 0.588259 | 0.079296 | 0.432842 | 0.743676 |
| Age at Dx Recode 5 Groups[50-59] | 0.540838 | 0.068976 | 0.405646 | 0.676029 |
| Age at Dx Recode 5 Groups[60-69] | 0.048875 | 0.067135 | -0.082708 | 0.180458 |
| Age at Dx Recode 5 Groups[70-79] | -0.343684 | 0.067486 | -0.475954 | -0.211414 |
| σ | 1.434866 | 0.026359 | 1.383203 | 1.486530 |
| **Wald Tests** | | | | |
| **Source** | **Nparm** | **DF** | **Wald ChiSquare** | **Prob>ChiSq** |
| Age at Dx Recode 5 Groups | 4 | 4 | 200.4892 | <.0001 |

**Table S33. Preliminary model parameter estimates for the cause-specific survival for 2000-03 stage IV colorectal cancer (CRC) in the rectum.** Parameter estimates correspond to the highlighted distribution in **Table S32**, demonstrating that sex is not a significant covariate. The Wald confidence interval was used.

| **2000-03 Stage IV CRC, Rectum: Preliminary Model Comparison** | | | | |
| --- | --- | --- | --- | --- |
| **Parameter Estimates** | | | | |
| **Term** | **Estimate** | **Std Error** | **Lower 95%** | **Upper 95%** |
| Intercept | -0.163711 | 0.036939 | -0.236111 | -0.091311 |
| Sex[Female] | 0.037664 | 0.036799 | -0.034460 | 0.109789 |
| Age at Dx Recode 5 Groups[20-49] | 0.590337 | 0.079294 | 0.434923 | 0.745750 |
| Age at Dx Recode 5 Groups[50-59] | 0.545970 | 0.069134 | 0.410469 | 0.681470 |
| Age at Dx Recode 5 Groups[60-69] | 0.053490 | 0.067261 | -0.078340 | 0.185320 |
| Age at Dx Recode 5 Groups[70-79] | -0.343140 | 0.067464 | -0.475366 | -0.210914 |
| σ | 1.434349 | 0.026350 | 1.382704 | 1.485994 |
| **Wald Tests** | | | | |
| **Source** | **Nparm** | **DF** | **Wald ChiSquare** | **Prob>ChiSq** |
| Sex | 1 | 1 | 1.0476 | 0.3061 |
| Age at Dx Recode 5 Groups | 4 | 4 | 200.9993 | <.0001 |

**Table S34. Diagnostics, coefficients and effect tests for final multinomial logistic regression model of colorectal cancer (CRC) stage conditioned on CRC size.** Probabilities that achieved statistical significance (α ≤ 0.05) are in bold.

| **Whole Model Test** | | | | |
| --- | --- | --- | --- | --- |
| **Model** | **-LogLikelihood** | **DF** | **ChiSquare** | **Prob>ChiSq** |
| Difference | 1347.124 | 6 | 2694.248 | **<.0001** |
| Full | 43493.676 |  | | |
| Reduced | 44840.800 |  |  |  |
| RSquare (U) | 0.0300 |  | | |
| AICc | 87005.4 |  |  |  |
| BIC | 87081.1 |  |  |  |
| Observations (or Sum Wgts) | 33485 |  |  |  |
| **Lack of Fit** | | | | |
| **Source** | **DF** | **-LogLikelihood** | **ChiSquare** | **Prob>ChiSq** |
| Lack of Fit | 282 | 245.133 | 490.2656 | **<.0001** |
| Saturated | 288 | 43248.543 |  | |
| Fitted | 6 | 43493.676 |  |  |
| **Parameter Estimates** | | | | |
| **Term** | **Estimate** | **Std Error** | **ChiSquare** | **Prob>ChiSq** |
| Intercept ${[\alpha}_{1}]$ | 4.45118078 | 0.2818035 | 249.49 | **<.0001** |
| Sqrt(Size (mm)) ${[\beta}_{1}$] | -0.8845893 | 0.0854211 | 107.24 | **<.0001** |
| Size (mm) [$\gamma_{1}$] | 0.02665437 | 0.0063795 | 17.46 | **<.0001** |
| Intercept ${[\alpha}_{2}]$ | 0.29839421 | 0.2763527 | 1.17 | 0.2802 |
| Sqrt(Size (mm)) $\beta_{2}$ | 0.16069904 | 0.0788434 | 4.15 | **0.0415** |
| Size (mm) [$\gamma_{2}$] | -0.015023 | 0.0055023 | 7.45 | **0.0063** |
| Intercept ${[\alpha}_{3}]$ | 0.41562304 | 0.2894097 | 2.06 | 0.1510 |
| Sqrt(Size (mm)) $\beta_{3}$ | 0.10670585 | 0.0833045 | 1.64 | 0.2002 |
| Size (mm) [$\gamma_{3}$] | -0.0169 | 0.0058694 | 8.29 | **0.0040** |
| **Effect Wald Tests** | | | | |
| **Source** | **Nparm** | **DF** | **Wald ChiSquare** | **Prob>ChiSq** |
| Sqrt(Size (mm)) | 3 | 3 | 242.02246 | **<.0001** |
| Size (mm) | 3 | 3 | 69.3294212 | **<.0001** |

**Table S35. Extrapolated counts of colorectal cancer size from three Poisson models for different rounding scenarios.** The number of size buckets was increased until ~1,453 overall observations were obtained (actual n = 1,484). Values rounded for visual simplicity. n/a = not applicable

| **Size (mm)** | **Counts from Poisson Model 1 (nearest centimeter)** | **Counts from**  **Poisson Model 2**  **(nearest half-centimeter)** | **Counts from Poisson Model 3**  **(nearest millimeter)** |
| --- | --- | --- | --- |
| **98** | n/a | n/a | 2 |
| **99** | n/a | n/a | 2 |
| **100** | 532 | n/a | n/a |
| **101** | n/a | n/a | 2 |
| **102** | n/a | n/a | 2 |
| **103** | n/a | n/a | 2 |
| **104** | n/a | n/a | 2 |
| **105** | n/a | 52 | n/a |
| **106** | n/a | n/a | 1 |
| **107** | n/a | n/a | 1 |
| **108** | n/a | n/a | 1 |
| **109** | n/a | n/a | 1 |
| **110** | 348 | n/a | n/a |
| **111** | n/a | n/a | 1 |
| **112** | n/a | n/a | 1 |
| **113** | n/a | n/a | 1 |
| **114** | n/a | n/a | 1 |
| **115** | n/a | 28 | n/a |
| **116** | n/a | n/a | 1 |
| **117** | n/a | n/a | 1 |
| **118** | n/a | n/a | 1 |
| **119** | n/a | n/a | 1 |
| **120** | 227 | n/a | n/a |
| **121** | n/a | n/a | 1 |
| **122** | n/a | n/a | 0 |
| **123** | n/a | n/a | 0 |
| **124** | n/a | n/a | 0 |
| **125** | n/a | 15 | n/a |
| **126** | n/a | n/a | 0 |
| **127** | n/a | n/a | 0 |
| **128** | n/a | n/a | 0 |
| **129** | n/a | n/a | 0 |
| **130** | 148 | n/a | n/a |
| **131** | n/a | n/a | 0 |
| **132** | n/a | n/a | 0 |
| **133** | n/a | n/a | 0 |
| **134** | n/a | n/a | 0 |
| **135** | n/a | 8 | n/a |
| **136** | n/a | n/a | 0 |
| **137** | n/a | n/a | 0 |
| **138** | n/a | n/a | 0 |
| **139** | n/a | n/a | 0 |
| **140** | 97 | n/a | n/a |

**Table S36. Screening outcomes per 1,000 individuals aged 40 free from diagnosed CRC based on colonoscopy screening strategies.** Outcomes generated using period life tables. Outcomes table modeled from Zauber et al.^1^ COL, colonoscopy; SIG, flexible sigmoidoscopy; CTC, computed tomographic colonography; CRC, colorectal cancer; LY, life-years; LYG, life-years gained compared with no screening. ^a^Maximum possible number with this strategy. ^b^Including deaths from complications of screening. ^c^Compared to no screening.

| **Strategy** | **Outcomes per 1,000 persons free of diagnosed cancer at age 40** | | | | | | | | | | | |  | **Reductions^c^ (%)** | |
| --- | --- | --- | --- | --- | --- | --- | --- | --- | --- | --- | --- | --- | --- | --- | --- |
| **Modality** | **Screening tests** | | | | **Follow-up COLs** | **Surveillance COLs** | **COLs for symptoms** | **Total COLs** | **Complications** | **CRC cases** | **CRC deaths^b^** | **LY with CRC** | **LYG** |  |  |
| **Age to begin-age to end, screening interval (# of tests^a^)** | **Stool tests** | **SIGs** | **CTCs** | **COLs** |  |  |  |  |  |  |  |  |  | **Incidence** | **Mortality** |
| No screening | 0 | 0 | 0 | 0 | 0 | 0 | 71 | 71 | 2 | 71.1 | 31.7 | 560.3 | 0.0 | 0.0% | 0.0% |
| COL 45-75, 5 (7) | 0 | 0 | 0 | 5,098 | 0 | 1,940 | 3 | 7,041 | 18 | 5.8 | 1.9 | 72.1 | 326.5 | 91.8% | 93.9% |
| COL 45-75, 10 (4) | 0 | 0 | 0 | 3,087 | 0 | 1,771 | 4 | 4,862 | 16 | 7.9 | 2.7 | 94.5 | 317.5 | 88.9% | 91.6% |
| COL 45-75, 15 (3) | 0 | 0 | 0 | 2,398 | 0 | 1,633 | 6 | 4,037 | 15 | 10.4 | 3.6 | 120.5 | 305.2 | 85.4% | 88.6% |
| COL 45-80, 5 (8) | 0 | 0 | 0 | 5,451 | 0 | 1,960 | 2 | 7,414 | 20 | 5.0 | 1.6 | 70.1 | 328.2 | 93.0% | 95.0% |
| COL 45-80, 10 (4) | 0 | 0 | 0 | 3,087 | 0 | 1,771 | 4 | 4,862 | 16 | 7.9 | 2.7 | 94.5 | 317.5 | 88.9% | 91.6% |
| COL 45-80, 15 (3) | 0 | 0 | 0 | 2,398 | 0 | 1,633 | 6 | 4,037 | 15 | 10.4 | 3.6 | 120.5 | 305.2 | 85.4% | 88.6% |
| COL 45-85, 5 (9) | 0 | 0 | 0 | 5,688 | 0 | 1,972 | 2 | 7,662 | 22 | 4.7 | 1.4 | 70.0 | 328.8 | 93.4% | 95.5% |
| COL 45-85, 10 (5) | 0 | 0 | 0 | 3,347 | 0 | 1,795 | 3 | 5,145 | 19 | 7.0 | 2.2 | 93.8 | 319.1 | 90.2% | 93.0% |
| COL 45-85, 15 (3) | 0 | 0 | 0 | 2,398 | 0 | 1,633 | 6 | 4,037 | 15 | 10.4 | 3.6 | 120.5 | 305.2 | 85.4% | 88.6% |
| COL 50-75, 5 (6) | 0 | 0 | 0 | 4,225 | 0 | 1,785 | 5 | 6,015 | 18 | 8.0 | 2.8 | 115.6 | 304.7 | 88.7% | 91.1% |
| COL 50-75, 10 (3) | 0 | 0 | 0 | 2,409 | 0 | 1,606 | 8 | 4,023 | 14 | 11.5 | 4.2 | 139.0 | 293.1 | 83.9% | 86.7% |
| COL 50-75, 15 (2) | 0 | 0 | 0 | 1,762 | 0 | 1,454 | 12 | 3,228 | 12 | 15.7 | 6.1 | 163.7 | 277.7 | 78.0% | 80.7% |
| COL 50-80, 5 (7) | 0 | 0 | 0 | 4,580 | 0 | 1,806 | 4 | 6,389 | 20 | 7.2 | 2.5 | 113.7 | 306.6 | 89.9% | 92.3% |
| COL 50-80, 10 (4) | 0 | 0 | 0 | 2,796 | 0 | 1,650 | 5 | 4,451 | 17 | 9.2 | 3.2 | 133.5 | 297.8 | 87.0% | 89.8% |
| COL 50-80, 15 (3) | 0 | 0 | 0 | 2,179 | 0 | 1,521 | 7 | 3,707 | 16 | 11.8 | 4.2 | 157.2 | 286.7 | 83.4% | 86.7% |
| COL 50-85, 5 (8) | 0 | 0 | 0 | 4,816 | 0 | 1,818 | 3 | 6,637 | 22 | 6.9 | 2.3 | 113.9 | 306.8 | 90.3% | 92.7% |
| COL 50-85, 10 (4) | 0 | 0 | 0 | 2,796 | 0 | 1,650 | 5 | 4,451 | 17 | 9.2 | 3.2 | 133.5 | 297.8 | 87.0% | 89.8% |
| COL 50-85, 15 (3) | 0 | 0 | 0 | 2,179 | 0 | 1,521 | 7 | 3,707 | 16 | 11.8 | 4.2 | 157.2 | 286.7 | 83.4% | 86.7% |
| COL 55-75, 5 (5) | 0 | 0 | 0 | 3,406 | 0 | 1,591 | 7 | 5,004 | 17 | 11.6 | 4.2 | 176.9 | 273.9 | 83.7% | 86.6% |
| COL 55-75, 10 (3) | 0 | 0 | 0 | 2,201 | 0 | 1,473 | 8 | 3,682 | 15 | 13.3 | 4.9 | 193.0 | 267.3 | 81.3% | 84.7% |
| COL 55-75, 15 (2) | 0 | 0 | 0 | 1,642 | 0 | 1,360 | 11 | 3,014 | 13 | 16.5 | 6.2 | 211.0 | 256.1 | 76.8% | 80.3% |
| COL 55-80, 5 (6) | 0 | 0 | 0 | 3,759 | 0 | 1,612 | 6 | 5,378 | 20 | 10.7 | 3.9 | 175.5 | 275.3 | 84.9% | 87.7% |
| COL 55-80, 10 (3) | 0 | 0 | 0 | 2,201 | 0 | 1,473 | 8 | 3,682 | 15 | 13.3 | 4.9 | 193.0 | 267.3 | 81.3% | 84.7% |
| COL 55-80, 15 (2) | 0 | 0 | 0 | 1,642 | 0 | 1,360 | 11 | 3,014 | 13 | 16.5 | 6.2 | 211.0 | 256.1 | 76.8% | 80.3% |
| COL 55-85, 5 (7) | 0 | 0 | 0 | 3,997 | 0 | 1,624 | 6 | 5,626 | 21 | 10.4 | 3.7 | 174.9 | 275.6 | 85.3% | 88.2% |
| COL 55-85, 10 (4) | 0 | 0 | 0 | 2,461 | 0 | 1,496 | 7 | 3,965 | 18 | 12.5 | 4.4 | 193.1 | 268.6 | 82.4% | 86.0% |
| COL 55-85, 15 (3) | 0 | 0 | 0 | 1,925 | 0 | 1,397 | 9 | 3,331 | 17 | 15.0 | 5.4 | 209.3 | 258.8 | 79.0% | 82.9% |

**Table S37. Screening outcomes per 1,000 individuals aged 40 free from diagnosed CRC based on multi-target stool DNA (mt-sDNA) screening strategies.** Outcomes generated using period life tables. Outcomes table modeled from Zauber et al.^1^ COL, colonoscopy; SIG, flexible sigmoidoscopy; CTC, computed tomographic colonography; CRC, colorectal cancer; LY, life-years; LYG, life-years gained compared with no screening. ^a^Maximum possible number with this strategy. ^b^Including deaths from complications of screening. ^c^Compared to no screening.

| **Strategy** | **Outcomes per 1,000 persons free of diagnosed cancer at age 40** | | | | | | | | | | | |  | **Reductions^c^ (%)** | |
| --- | --- | --- | --- | --- | --- | --- | --- | --- | --- | --- | --- | --- | --- | --- | --- |
| **Modality** | **Screening tests** | | | | **Follow-up COLs** | **Surveillance COLs** | **COLs for symptoms** | **Total COLs** | **Complications** | **CRC cases** | **CRC deaths^b^** | **LY with CRC** | **LYG** |  |  |
| **Age to begin-age to end, screening interval (# of tests^a^)** | **Stool tests** | **SIGs** | **CTCs** | **COLs** |  |  |  |  |  |  |  |  |  | **Incidence** | **Mortality** |
| No screening | 0 | 0 | 0 | 0 | 0 | 0 | 71 | 71 | 2 | 71.1 | 31.7 | 560.3 | 0.0 | 0.0% | 0.0% |
| mt-sDNA 45-75, 1 (31) | 13,168 | 0 | 0 | 0 | 1,592 | 1,502 | 8 | 3,103 | 12 | 13.2 | 4.4 | 154.2 | 305.3 | 81.4% | 86.3% |
| mt-sDNA 45-75, 3 (11) | 7,103 | 0 | 0 | 0 | 919 | 1,171 | 13 | 2,103 | 10 | 22.4 | 7.5 | 255.7 | 270.9 | 68.5% | 76.2% |
| mt-sDNA 45-75, 5 (7) | 5,212 | 0 | 0 | 0 | 702 | 989 | 18 | 1,708 | 9 | 29.0 | 10.2 | 314.3 | 236.9 | 59.3% | 67.7% |
| mt-sDNA 45-80, 1 (36) | 14,246 | 0 | 0 | 0 | 1,724 | 1,532 | 6 | 3,262 | 13 | 11.7 | 3.5 | 151.8 | 310.3 | 83.6% | 89.1% |
| mt-sDNA 45-80, 3 (12) | 7,704 | 0 | 0 | 0 | 999 | 1,201 | 10 | 2,210 | 11 | 20.7 | 6.3 | 255.2 | 278.8 | 70.8% | 80.0% |
| mt-sDNA 45-80, 5 (8) | 5,608 | 0 | 0 | 0 | 760 | 1,011 | 14 | 1,785 | 10 | 27.5 | 9.1 | 315.5 | 243.8 | 61.3% | 71.3% |
| mt-sDNA 45-85, 1 (41) | 15,022 | 0 | 0 | 0 | 1,820 | 1,547 | 4 | 3,371 | 15 | 11.0 | 3.0 | 153.5 | 312.6 | 84.6% | 90.6% |
| mt-sDNA 45-85, 3 (14) | 8,191 | 0 | 0 | 0 | 1,068 | 1,220 | 7 | 2,294 | 12 | 19.8 | 5.5 | 257.4 | 282.6 | 72.2% | 82.6% |
| mt-sDNA 45-85, 5 (9) | 5,880 | 0 | 0 | 0 | 800 | 1,022 | 12 | 1,834 | 11 | 27.2 | 8.6 | 318.6 | 245.6 | 61.8% | 73.0% |
| mt-sDNA 50-75, 1 (26) | 10,891 | 0 | 0 | 0 | 1,349 | 1,374 | 9 | 2,733 | 12 | 15.6 | 5.3 | 199.0 | 284.1 | 78.1% | 83.4% |
| mt-sDNA 50-75, 3 (9) | 5,958 | 0 | 0 | 0 | 797 | 1,055 | 14 | 1,866 | 10 | 25.0 | 8.5 | 298.4 | 250.2 | 64.8% | 73.3% |
| mt-sDNA 50-75, 5 (6) | 4,381 | 0 | 0 | 0 | 611 | 894 | 20 | 1,524 | 9 | 31.4 | 11.2 | 349.1 | 217.6 | 55.8% | 64.6% |
| mt-sDNA 50-80, 1 (31) | 11,979 | 0 | 0 | 0 | 1,483 | 1,402 | 7 | 2,892 | 13 | 14.1 | 4.4 | 197.6 | 288.4 | 80.2% | 86.2% |
| mt-sDNA 50-80, 3 (11) | 6,517 | 0 | 0 | 0 | 874 | 1,085 | 11 | 1,969 | 11 | 23.2 | 7.2 | 298.7 | 258.7 | 67.3% | 77.3% |
| mt-sDNA 50-80, 5 (7) | 4,781 | 0 | 0 | 0 | 670 | 914 | 16 | 1,599 | 10 | 30.0 | 10.1 | 351.6 | 224.4 | 57.9% | 68.2% |
| mt-sDNA 50-85, 1 (36) | 12,759 | 0 | 0 | 0 | 1,578 | 1,418 | 5 | 3,002 | 14 | 13.3 | 3.9 | 198.1 | 290.6 | 81.3% | 87.6% |
| mt-sDNA 50-85, 3 (12) | 6,957 | 0 | 0 | 0 | 933 | 1,099 | 9 | 2,041 | 12 | 22.5 | 6.5 | 301.7 | 261.6 | 68.3% | 79.4% |
| mt-sDNA 50-85, 5 (8) | 5,052 | 0 | 0 | 0 | 711 | 925 | 14 | 1,650 | 11 | 29.7 | 9.6 | 355.5 | 225.8 | 58.2% | 69.7% |
| mt-sDNA 55-75, 1 (21) | 8,752 | 0 | 0 | 0 | 1,120 | 1,207 | 12 | 2,339 | 12 | 19.4 | 6.7 | 259.8 | 253.1 | 72.7% | 78.8% |
| mt-sDNA 55-75, 3 (7) | 4,656 | 0 | 0 | 0 | 649 | 907 | 18 | 1,575 | 9 | 29.8 | 10.4 | 352.7 | 217.8 | 58.2% | 67.0% |
| mt-sDNA 55-75, 5 (5) | 3,578 | 0 | 0 | 0 | 520 | 770 | 23 | 1,313 | 9 | 35.3 | 12.8 | 394.3 | 190.3 | 50.4% | 59.5% |
| mt-sDNA 55-80, 1 (26) | 9,825 | 0 | 0 | 0 | 1,250 | 1,235 | 9 | 2,495 | 13 | 17.8 | 5.8 | 258.2 | 258.5 | 75.0% | 81.8% |
| mt-sDNA 55-80, 3 (9) | 5,411 | 0 | 0 | 0 | 753 | 950 | 13 | 1,716 | 11 | 27.2 | 8.7 | 352.5 | 227.7 | 61.8% | 72.4% |
| mt-sDNA 55-80, 5 (6) | 3,979 | 0 | 0 | 0 | 580 | 792 | 19 | 1,391 | 10 | 33.8 | 11.7 | 395.8 | 196.1 | 52.5% | 63.1% |
| mt-sDNA 55-85, 1 (31) | 10,604 | 0 | 0 | 0 | 1,347 | 1,252 | 8 | 2,607 | 14 | 17.1 | 5.3 | 258.8 | 260.9 | 76.0% | 83.2% |
| mt-sDNA 55-85, 3 (11) | 5,813 | 0 | 0 | 0 | 810 | 967 | 11 | 1,787 | 12 | 26.5 | 8.1 | 355.7 | 231.0 | 62.7% | 74.4% |
| mt-sDNA 55-85, 5 (7) | 4,252 | 0 | 0 | 0 | 622 | 805 | 16 | 1,443 | 11 | 33.5 | 11.1 | 401.9 | 199.2 | 52.9% | 65.1% |

**Table S38. Screening outcomes per 1,000 individuals aged 40 free from diagnosed CRC based on fecal immunochemical test (FIT) screening strategies.** Outcomes generated using period life tables. Outcomes table modeled from Zauber et al.^1^ COL, colonoscopy; SIG, flexible sigmoidoscopy; CTC, computed tomographic colonography; CRC, colorectal cancer; LY, life-years; LYG, life-years gained compared with no screening. ^a^Maximum possible number with this strategy. ^b^Including deaths from complications of screening. ^c^Compared to no screening.

| **Strategy** | **Outcomes per 1,000 persons free of diagnosed cancer at age 40** | | | | | | | | | | | |  | **Reductions^c^ (%)** | |
| --- | --- | --- | --- | --- | --- | --- | --- | --- | --- | --- | --- | --- | --- | --- | --- |
| **Modality** | **Screening tests** | | | | **Follow-up COLs** | **Surveillance COLs** | **COLs for symptoms** | **Total COLs** | **Complications** | **CRC cases** | **CRC deaths^b^** | **LY with CRC** | **LYG** |  |  |
| **Age to begin-age to end, screening interval (# of tests^a^)** | **Stool tests** | **SIGs** | **CTCs** | **COLs** |  |  |  |  |  |  |  |  |  | **Incidence** | **Mortality** |
| No screening | 0 | 0 | 0 | 0 | 0 | 0 | 71 | 71 | 2 | 71.1 | 31.7 | 560.3 | 0.0 | 0.0% | 0.0% |
| FIT 45-75, 1 (31) | 18,887 | 0 | 0 | 0 | 918 | 1,258 | 11 | 2,187 | 10 | 19.5 | 6.1 | 234.0 | 288.2 | 72.6% | 80.7% |
| FIT 45-75, 2 (16) | 11,480 | 0 | 0 | 0 | 607 | 983 | 16 | 1,606 | 9 | 28.3 | 9.2 | 324.2 | 254.8 | 60.3% | 70.9% |
| FIT 45-75, 3 (11) | 8,345 | 0 | 0 | 0 | 463 | 806 | 21 | 1,291 | 7 | 34.9 | 12.0 | 377.5 | 222.9 | 50.9% | 62.0% |
| FIT 45-80, 1 (36) | 20,544 | 0 | 0 | 0 | 1,003 | 1,288 | 7 | 2,298 | 11 | 17.5 | 4.9 | 231.6 | 295.6 | 75.4% | 84.7% |
| FIT 45-80, 2 (18) | 12,314 | 0 | 0 | 0 | 655 | 1,009 | 12 | 1,676 | 9 | 26.7 | 8.0 | 326.8 | 262.6 | 62.4% | 74.9% |
| FIT 45-80, 3 (12) | 8,929 | 0 | 0 | 0 | 501 | 828 | 18 | 1,346 | 8 | 33.6 | 10.9 | 380.7 | 228.9 | 52.7% | 65.4% |
| FIT 45-85, 1 (41) | 21,743 | 0 | 0 | 0 | 1,065 | 1,305 | 5 | 2,374 | 12 | 16.9 | 4.2 | 236.9 | 298.8 | 76.2% | 86.6% |
| FIT 45-85, 2 (21) | 13,219 | 0 | 0 | 0 | 710 | 1,026 | 8 | 1,744 | 11 | 26.0 | 7.0 | 334.3 | 267.2 | 63.4% | 78.0% |
| FIT 45-85, 3 (14) | 9,607 | 0 | 0 | 0 | 546 | 847 | 13 | 1,406 | 9 | 32.7 | 9.7 | 388.4 | 236.2 | 54.0% | 69.2% |
| FIT 50-75, 1 (26) | 15,563 | 0 | 0 | 0 | 789 | 1,136 | 12 | 1,937 | 10 | 22.1 | 7.2 | 277.9 | 265.3 | 68.9% | 77.4% |
| FIT 50-75, 2 (13) | 9,265 | 0 | 0 | 0 | 515 | 869 | 19 | 1,402 | 8 | 31.6 | 10.7 | 364.3 | 230.2 | 55.6% | 66.2% |
| FIT 50-75, 3 (9) | 6,865 | 0 | 0 | 0 | 402 | 709 | 24 | 1,135 | 7 | 37.9 | 13.3 | 411.1 | 201.4 | 46.8% | 57.9% |
| FIT 50-80, 1 (31) | 17,225 | 0 | 0 | 0 | 874 | 1,169 | 9 | 2,052 | 11 | 20.2 | 5.8 | 278.2 | 273.4 | 71.5% | 81.5% |
| FIT 50-80, 2 (16) | 10,520 | 0 | 0 | 0 | 589 | 908 | 13 | 1,510 | 9 | 29.3 | 8.9 | 367.0 | 241.7 | 58.9% | 72.0% |
| FIT 50-80, 3 (11) | 7,677 | 0 | 0 | 0 | 455 | 739 | 18 | 1,213 | 8 | 35.9 | 11.6 | 416.7 | 212.0 | 49.5% | 63.2% |
| FIT 50-85, 1 (36) | 18,423 | 0 | 0 | 0 | 937 | 1,184 | 7 | 2,128 | 12 | 19.6 | 5.2 | 280.9 | 276.2 | 72.4% | 83.5% |
| FIT 50-85, 2 (18) | 11,130 | 0 | 0 | 0 | 625 | 921 | 10 | 1,557 | 10 | 28.7 | 8.2 | 371.0 | 244.9 | 59.6% | 74.2% |
| FIT 50-85, 3 (12) | 8,105 | 0 | 0 | 0 | 483 | 751 | 16 | 1,250 | 9 | 35.3 | 10.9 | 419.8 | 215.3 | 50.4% | 65.4% |
| FIT 55-75, 1 (21) | 12,373 | 0 | 0 | 0 | 664 | 980 | 15 | 1,658 | 10 | 26.4 | 8.7 | 337.1 | 235.3 | 62.9% | 72.4% |
| FIT 55-75, 2 (11) | 7,593 | 0 | 0 | 0 | 449 | 749 | 21 | 1,218 | 8 | 35.3 | 12.0 | 413.8 | 203.5 | 50.4% | 62.2% |
| FIT 55-75, 3 (7) | 5,305 | 0 | 0 | 0 | 329 | 597 | 28 | 954 | 7 | 42.3 | 15.4 | 450.2 | 170.6 | 40.6% | 51.3% |
| FIT 55-80, 1 (26) | 14,050 | 0 | 0 | 0 | 749 | 1,015 | 11 | 1,776 | 11 | 24.4 | 7.4 | 337.1 | 243.0 | 65.7% | 76.6% |
| FIT 55-80, 2 (13) | 8,439 | 0 | 0 | 0 | 499 | 777 | 17 | 1,292 | 9 | 33.6 | 10.6 | 416.3 | 212.0 | 52.7% | 66.6% |
| FIT 55-80, 3 (9) | 6,263 | 0 | 0 | 0 | 393 | 642 | 22 | 1,057 | 8 | 39.6 | 13.2 | 454.3 | 184.4 | 44.3% | 58.2% |
| FIT 55-85, 1 (31) | 15,246 | 0 | 0 | 0 | 813 | 1,033 | 9 | 1,854 | 12 | 23.7 | 6.8 | 339.3 | 245.9 | 66.7% | 78.7% |
| FIT 55-85, 2 (16) | 9,352 | 0 | 0 | 0 | 555 | 798 | 13 | 1,366 | 10 | 32.7 | 9.5 | 422.2 | 217.2 | 54.0% | 69.9% |
| FIT 55-85, 3 (11) | 6,851 | 0 | 0 | 0 | 434 | 658 | 18 | 1,110 | 9 | 39.0 | 12.2 | 462.1 | 189.4 | 45.1% | 61.4% |

**Table S39. Screening outcomes per 1,000 individuals aged 40 free from diagnosed CRC based on guaiac-based fecal occult blood test (HSgFOBT) screening strategies.** Outcomes generated using period life tables. Outcomes table modeled from Zauber et al.^1^ COL, colonoscopy; SIG, flexible sigmoidoscopy; CTC, computed tomographic colonography; CRC, colorectal cancer; LY, life-years; LYG, life-years gained compared with no screening. ^a^Maximum possible number with this strategy. ^b^Including deaths from complications of screening. ^c^Compared to no screening.

| **Strategy** | **Outcomes per 1,000 persons free of diagnosed cancer at age 40** | | | | | | | | | | | |  | **Reductions^c^ (%)** | |
| --- | --- | --- | --- | --- | --- | --- | --- | --- | --- | --- | --- | --- | --- | --- | --- |
| **Modality** | **Screening tests** | | | | **Follow-up COLs** | **Surveillance COLs** | **COLs for symptoms** | **Total COLs** | **Complications** | **CRC cases** | **CRC deaths^b^** | **LY with CRC** | **LYG** |  |  |
| **Age to begin-age to end, screening interval (# of tests^a^)** | **Stool tests** | **SIGs** | **CTCs** | **COLs** |  |  |  |  |  |  |  |  |  | **Incidence** | **Mortality** |
| No screening | 0 | 0 | 0 | 0 | 0 | 0 | 71 | 71 | 2 | 71.1 | 31.7 | 560.3 | 0.0 | 0.0% | 0.0% |
| HSgFOBT 45-75, 1 (31) | 15,781 | 0 | 0 | 0 | 1,329 | 1,264 | 10 | 2,603 | 11 | 18.2 | 5.9 | 213.9 | 290.5 | 74.5% | 81.5% |
| HSgFOBT 45-75, 2 (16) | 10,436 | 0 | 0 | 0 | 911 | 1,013 | 15 | 1,939 | 9 | 26.3 | 8.7 | 299.7 | 258.9 | 63.0% | 72.4% |
| HSgFOBT 45-75, 3 (11) | 7,801 | 0 | 0 | 0 | 697 | 840 | 21 | 1,558 | 8 | 33.1 | 11.7 | 353.0 | 225.3 | 53.5% | 63.2% |
| HSgFOBT 45-80, 1 (36) | 17,149 | 0 | 0 | 0 | 1,444 | 1,294 | 7 | 2,745 | 12 | 16.4 | 4.7 | 213.3 | 297.2 | 77.0% | 85.3% |
| HSgFOBT 45-80, 2 (18) | 11,180 | 0 | 0 | 0 | 978 | 1,038 | 12 | 2,028 | 10 | 24.7 | 7.6 | 300.6 | 265.3 | 65.2% | 76.2% |
| HSgFOBT 45-80, 3 (12) | 8,434 | 0 | 0 | 0 | 757 | 865 | 17 | 1,639 | 9 | 31.6 | 10.4 | 356.1 | 233.3 | 55.6% | 67.1% |
| HSgFOBT 45-85, 1 (41) | 18,122 | 0 | 0 | 0 | 1,529 | 1,310 | 5 | 2,844 | 13 | 15.6 | 4.1 | 214.8 | 299.6 | 78.1% | 87.0% |
| HSgFOBT 45-85, 2 (21) | 11,992 | 0 | 0 | 0 | 1,053 | 1,057 | 8 | 2,118 | 11 | 23.9 | 6.6 | 305.6 | 270.3 | 66.4% | 79.3% |
| HSgFOBT 45-85, 3 (14) | 9,034 | 0 | 0 | 0 | 815 | 884 | 13 | 1,713 | 10 | 30.6 | 9.4 | 361.2 | 239.0 | 56.9% | 70.3% |
| HSgFOBT 50-75, 1 (26) | 13,084 | 0 | 0 | 0 | 1,124 | 1,148 | 12 | 2,285 | 11 | 20.9 | 6.9 | 261.0 | 268.2 | 70.6% | 78.3% |
| HSgFOBT 50-75, 2 (13) | 8,462 | 0 | 0 | 0 | 757 | 901 | 18 | 1,676 | 9 | 29.7 | 10.2 | 341.7 | 233.7 | 58.2% | 67.8% |
| HSgFOBT 50-75, 3 (9) | 6,502 | 0 | 0 | 0 | 597 | 746 | 23 | 1,366 | 8 | 35.7 | 12.8 | 388.3 | 205.2 | 49.7% | 59.7% |
| HSgFOBT 50-80, 1 (31) | 14,447 | 0 | 0 | 0 | 1,241 | 1,180 | 9 | 2,429 | 12 | 19.1 | 5.7 | 259.4 | 274.3 | 73.2% | 81.9% |
| HSgFOBT 50-80, 2 (16) | 9,583 | 0 | 0 | 0 | 859 | 940 | 13 | 1,812 | 10 | 27.3 | 8.5 | 343.3 | 244.8 | 61.6% | 73.3% |
| HSgFOBT 50-80, 3 (11) | 7,196 | 0 | 0 | 0 | 664 | 774 | 18 | 1,456 | 9 | 33.9 | 11.3 | 391.6 | 214.2 | 52.4% | 64.5% |
| HSgFOBT 50-85, 1 (36) | 15,430 | 0 | 0 | 0 | 1,325 | 1,197 | 7 | 2,528 | 13 | 18.2 | 5.1 | 261.4 | 277.5 | 74.4% | 84.0% |
| HSgFOBT 50-85, 2 (18) | 10,127 | 0 | 0 | 0 | 909 | 952 | 11 | 1,872 | 11 | 26.8 | 7.8 | 346.6 | 247.1 | 62.4% | 75.3% |
| HSgFOBT 50-85, 3 (12) | 7,661 | 0 | 0 | 0 | 709 | 785 | 16 | 1,510 | 9 | 33.4 | 10.6 | 396.8 | 217.3 | 53.1% | 66.6% |
| HSgFOBT 55-75, 1 (21) | 10,504 | 0 | 0 | 0 | 928 | 999 | 15 | 1,942 | 10 | 25.0 | 8.4 | 319.4 | 237.4 | 64.9% | 73.4% |
| HSgFOBT 55-75, 2 (11) | 6,969 | 0 | 0 | 0 | 644 | 780 | 20 | 1,444 | 9 | 33.5 | 11.5 | 394.6 | 206.2 | 53.0% | 63.6% |
| HSgFOBT 55-75, 3 (7) | 5,055 | 0 | 0 | 0 | 479 | 630 | 28 | 1,136 | 7 | 40.5 | 15.0 | 431.4 | 172.9 | 43.1% | 52.7% |
| HSgFOBT 55-80, 1 (26) | 11,872 | 0 | 0 | 0 | 1,044 | 1,030 | 11 | 2,086 | 12 | 23.1 | 7.2 | 318.9 | 244.5 | 67.6% | 77.3% |
| HSgFOBT 55-80, 2 (13) | 7,723 | 0 | 0 | 0 | 714 | 807 | 16 | 1,538 | 10 | 31.8 | 10.3 | 395.2 | 213.5 | 55.3% | 67.4% |
| HSgFOBT 55-80, 3 (9) | 5,941 | 0 | 0 | 0 | 565 | 672 | 21 | 1,259 | 8 | 37.8 | 12.8 | 435.2 | 187.2 | 46.8% | 59.6% |
| HSgFOBT 55-85, 1 (31) | 12,862 | 0 | 0 | 0 | 1,129 | 1,048 | 9 | 2,186 | 13 | 22.3 | 6.5 | 320.3 | 247.3 | 68.7% | 79.4% |
| HSgFOBT 55-85, 2 (16) | 8,542 | 0 | 0 | 0 | 789 | 828 | 13 | 1,630 | 11 | 30.8 | 9.2 | 400.3 | 219.4 | 56.7% | 70.9% |
| HSgFOBT 55-85, 3 (11) | 6,442 | 0 | 0 | 0 | 616 | 689 | 18 | 1,323 | 9 | 37.0 | 11.9 | 439.7 | 191.2 | 48.0% | 62.4% |

**Table S40. Screening outcomes per 1,000 individuals aged 40 free from diagnosed CRC based on sigmoidoscopy screening strategies.** Outcomes generated using period life tables. Outcomes table modeled from Zauber et al.^1^ COL, colonoscopy; SIG, flexible sigmoidoscopy; CTC, computed tomographic colonography; CRC, colorectal cancer; LY, life-years; LYG, life-years gained compared with no screening. ^a^Maximum possible number with this strategy. ^b^Including deaths from complications of screening. ^c^Compared to no screening.

| **Strategy** | **Outcomes per 1,000 persons free of diagnosed cancer at age 40** | | | | | | | | | | | |  | **Reductions^c^ (%)** | |
| --- | --- | --- | --- | --- | --- | --- | --- | --- | --- | --- | --- | --- | --- | --- | --- |
| **Modality** | **Screening tests** | | | | **Follow-up COLs** | **Surveillance COLs** | **COLs for symptoms** | **Total COLs** | **Complications** | **CRC cases** | **CRC deaths^b^** | **LY with CRC** | **LYG** |  |  |
| **Age to begin-age to end, screening interval (# of tests^a^)** | **Stool tests** | **SIGs** | **CTCs** | **COLs** |  |  |  |  |  |  |  |  |  | **Incidence** | **Mortality** |
| No screening | 0 | 0 | 0 | 0 | 0 | 0 | 71 | 71 | 2 | 71.1 | 31.7 | 560.3 | 0.0 | 0.0% | 0.0% |
| SIG 45-75, 5 (7) | 0 | 5,170 | 0 | 0 | 754 | 835 | 28 | 1,618 | 8 | 31.0 | 12.6 | 274.5 | 207.6 | 56.4% | 60.1% |
| SIG 45-75, 10 (4) | 0 | 3,275 | 0 | 0 | 538 | 743 | 30 | 1,311 | 8 | 33.9 | 13.8 | 302.3 | 193.2 | 52.4% | 56.5% |
| SIG 45-80, 5 (8) | 0 | 5,587 | 0 | 0 | 811 | 852 | 26 | 1,689 | 9 | 29.6 | 11.9 | 271.3 | 210.9 | 58.4% | 62.3% |
| SIG 45-80, 10 (4) | 0 | 3,275 | 0 | 0 | 538 | 743 | 30 | 1,311 | 8 | 33.9 | 13.8 | 302.3 | 193.2 | 52.4% | 56.5% |
| SIG 45-85, 5 (9) | 0 | 5,876 | 0 | 0 | 852 | 861 | 25 | 1,738 | 10 | 29.0 | 11.6 | 271.9 | 212.3 | 59.2% | 63.3% |
| SIG 45-85, 10 (5) | 0 | 3,606 | 0 | 0 | 594 | 760 | 28 | 1,382 | 9 | 32.7 | 13.0 | 302.6 | 196.3 | 54.0% | 58.8% |
| SIG 50-75, 5 (6) | 0 | 4,335 | 0 | 0 | 650 | 774 | 29 | 1,452 | 8 | 32.4 | 13.2 | 300.7 | 193.8 | 54.5% | 58.2% |
| SIG 50-75, 10 (3) | 0 | 2,527 | 0 | 0 | 430 | 670 | 33 | 1,133 | 7 | 36.9 | 15.3 | 327.5 | 176.1 | 48.1% | 51.7% |
| SIG 50-80, 5 (7) | 0 | 4,751 | 0 | 0 | 707 | 791 | 27 | 1,525 | 9 | 31.0 | 12.5 | 298.1 | 197.4 | 56.4% | 60.4% |
| SIG 50-80, 10 (4) | 0 | 3,003 | 0 | 0 | 509 | 700 | 29 | 1,239 | 8 | 34.2 | 13.9 | 324.6 | 182.7 | 51.9% | 56.2% |
| SIG 50-85, 5 (8) | 0 | 5,039 | 0 | 0 | 747 | 800 | 26 | 1,573 | 10 | 30.5 | 12.2 | 298.1 | 198.8 | 57.2% | 61.5% |
| SIG 50-85, 10 (4) | 0 | 3,003 | 0 | 0 | 509 | 700 | 29 | 1,239 | 8 | 34.2 | 13.9 | 324.6 | 182.7 | 51.9% | 56.2% |
| SIG 55-75, 5 (5) | 0 | 3,524 | 0 | 0 | 549 | 694 | 30 | 1,274 | 8 | 34.6 | 14.2 | 335.9 | 174.4 | 51.3% | 55.2% |
| SIG 55-75, 10 (3) | 0 | 2,337 | 0 | 0 | 414 | 624 | 32 | 1,070 | 7 | 37.1 | 15.2 | 356.6 | 163.3 | 47.8% | 52.1% |
| SIG 55-80, 5 (6) | 0 | 3,941 | 0 | 0 | 606 | 711 | 29 | 1,345 | 9 | 33.3 | 13.5 | 334.0 | 177.7 | 53.2% | 57.5% |
| SIG 55-80, 10 (3) | 0 | 2,337 | 0 | 0 | 414 | 624 | 32 | 1,070 | 7 | 37.1 | 15.2 | 356.6 | 163.3 | 47.8% | 52.1% |
| SIG 55-85, 5 (7) | 0 | 4,230 | 0 | 0 | 646 | 720 | 27 | 1,393 | 10 | 32.7 | 13.2 | 333.8 | 179.0 | 54.0% | 58.4% |
| SIG 55-85, 10 (4) | 0 | 2,668 | 0 | 0 | 470 | 642 | 30 | 1,142 | 9 | 36.0 | 14.5 | 357.3 | 166.6 | 49.4% | 54.4% |

**Table S41. Screening outcomes per 1,000 individuals aged 40 free from diagnosed CRC based on sigmoidoscopy plus fecal immunochemical test (SIG+FIT) screening strategies.** Outcomes generated using period life tables. Outcomes table modeled from Zauber et al.^1^ COL, colonoscopy; SIG, flexible sigmoidoscopy; CTC, computed tomographic colonography; CRC, colorectal cancer; LY, life-years; LYG, life-years gained compared with no screening. ^a^Maximum possible number with this strategy. ^b^Including deaths from complications of screening. ^c^Compared to no screening.

| **Strategy** | **Outcomes per 1,000 persons free of diagnosed cancer at age 40** | | | | | | | | | | | |  | **Reductions^c^ (%)** | |
| --- | --- | --- | --- | --- | --- | --- | --- | --- | --- | --- | --- | --- | --- | --- | --- |
| **Modality** | **Screening tests** | | | | **Follow-up COLs** | **Surveillance COLs** | **COLs for symptoms** | **Total COLs** | **Complications** | **CRC cases** | **CRC deaths^b^** | **LY with CRC** | **LYG** |  |  |
| **Age to begin-age to end, screening interval (# of tests^a^)** | **Stool tests** | **SIGs** | **CTCs** | **COLs** |  |  |  |  |  |  |  |  |  | **Incidence** | **Mortality** |
| No screening | 0 | 0 | 0 | 0 | 0 | 0 | 71 | 71 | 2 | 71.1 | 31.7 | 560.3 | 0.0 | 0.0% | 0.0% |
| SIG+FIT 45-75, 10_1 (4_31) | 16,639 | 2,281 | 0 | 0 | 1,149 | 1390.73 | 8.78 | 2,549 | 11 | 15.0 | 4.9 | 171.6 | 301.0 | 78.9% | 84.6% |
| SIG+FIT 45-75, 10_2 (4_16) | 10,081 | 2,626 | 0 | 0 | 928 | 1237.57 | 10.60 | 2,176 | 10 | 18.5 | 6.0 | 210.1 | 287.4 | 74.0% | 81.0% |
| SIG+FIT 45-75, 5_2 (7_16) | 9,327 | 4,216 | 0 | 0 | 1,091 | 1284.61 | 10.23 | 2,386 | 11 | 17.2 | 5.7 | 193.8 | 290.7 | 75.8% | 82.1% |
| SIG+FIT 45-75, 5_3 (7_11) | 6,772 | 4,469 | 0 | 0 | 1,009 | 1200.27 | 12.27 | 2,221 | 10 | 19.5 | 6.7 | 211.5 | 278.8 | 72.6% | 78.9% |
| SIG+FIT 45-80, 10_1 (4_36) | 17,984 | 2,424 | 0 | 0 | 1,238 | 1419.94 | 6.22 | 2,664 | 12 | 13.4 | 3.9 | 169.6 | 305.9 | 81.2% | 87.6% |
| SIG+FIT 45-80, 10_2 (4_18) | 10,711 | 2,709 | 0 | 0 | 974 | 1256.05 | 8.50 | 2,239 | 11 | 17.4 | 5.2 | 211.0 | 291.4 | 75.5% | 83.4% |
| SIG+FIT 45-80, 5_2 (8_18) | 10,005 | 4,534 | 0 | 0 | 1,174 | 1308.89 | 7.66 | 2,490 | 12 | 15.8 | 4.8 | 194.3 | 295.5 | 77.8% | 85.0% |
| SIG+FIT 45-80, 5_3 (8_12) | 7,309 | 4,779 | 0 | 0 | 1,083 | 1225.58 | 9.46 | 2,318 | 11 | 17.9 | 5.7 | 211.1 | 284.9 | 74.8% | 82.1% |
| SIG+FIT 45-85, 10_1 (5_41) | 19,068 | 2,512 | 0 | 0 | 1,307 | 1434.63 | 4.50 | 2,747 | 13 | 12.8 | 3.4 | 171.5 | 308.7 | 82.1% | 89.3% |
| SIG+FIT 45-85, 10_2 (5_21) | 11,510 | 2,874 | 0 | 0 | 1,046 | 1275.44 | 5.91 | 2,328 | 13 | 16.5 | 4.5 | 212.6 | 295.1 | 76.8% | 85.7% |
| SIG+FIT 45-85, 5_2 (9_21) | 10,654 | 4,723 | 0 | 0 | 1,238 | 1327.49 | 5.60 | 2,572 | 13 | 14.9 | 4.1 | 194.1 | 299.1 | 79.1% | 87.1% |
| SIG+FIT 45-85, 5_3 (9_14) | 7,783 | 5,003 | 0 | 0 | 1,144 | 1241.42 | 7.36 | 2,393 | 12 | 17.1 | 5.0 | 211.5 | 287.4 | 75.9% | 84.1% |
| SIG+FIT 50-75, 10_1 (3_26) | 13,586 | 1,936 | 0 | 0 | 984 | 1277.35 | 10.50 | 2,271 | 11 | 17.3 | 5.9 | 211.1 | 279.4 | 75.6% | 81.5% |
| SIG+FIT 50-75, 10_2 (3_13) | 8,073 | 2,132 | 0 | 0 | 775 | 1120.50 | 13.58 | 1,910 | 10 | 21.7 | 7.5 | 248.5 | 262.6 | 69.5% | 76.3% |
| SIG+FIT 50-75, 5_2 (6_13) | 7,568 | 3,567 | 0 | 0 | 936 | 1174.80 | 12.07 | 2,123 | 11 | 19.6 | 6.7 | 231.5 | 268.8 | 72.5% | 78.9% |
| SIG+FIT 50-75, 5_3 (6_9) | 5,625 | 3,752 | 0 | 0 | 870 | 1098.80 | 13.81 | 1,983 | 10 | 21.6 | 7.6 | 247.0 | 258.5 | 69.7% | 76.1% |
| SIG+FIT 50-80, 10_1 (4_31) | 15,094 | 2,101 | 0 | 0 | 1,085 | 1305.98 | 7.54 | 2,399 | 12 | 15.6 | 4.8 | 210.5 | 285.1 | 78.1% | 85.0% |
| SIG+FIT 50-80, 10_2 (4_16) | 9,179 | 2,403 | 0 | 0 | 881 | 1159.31 | 9.14 | 2,049 | 11 | 19.1 | 5.9 | 246.5 | 271.6 | 73.1% | 81.3% |
| SIG+FIT 50-80, 5_2 (7_16) | 8,503 | 3,849 | 0 | 0 | 1,028 | 1206.09 | 8.82 | 2,243 | 12 | 17.7 | 5.6 | 230.7 | 275.3 | 75.1% | 82.5% |
| SIG+FIT 50-80, 5_3 (7_11) | 6,201 | 4,080 | 0 | 0 | 951 | 1125.73 | 10.76 | 2,088 | 11 | 19.9 | 6.5 | 246.3 | 264.1 | 72.0% | 79.4% |
| SIG+FIT 50-85, 10_1 (4_36) | 16,050 | 2,209 | 0 | 0 | 1,152 | 1322.14 | 5.91 | 2,480 | 13 | 14.8 | 4.2 | 209.6 | 287.5 | 79.2% | 86.6% |
| SIG+FIT 50-85, 10_2 (4_18) | 9,636 | 2,466 | 0 | 0 | 916 | 1168.43 | 7.83 | 2,092 | 12 | 18.7 | 5.6 | 248.4 | 273.2 | 73.7% | 82.5% |
| SIG+FIT 50-85, 5_2 (8_18) | 8,993 | 4,070 | 0 | 0 | 1,088 | 1219.52 | 7.17 | 2,314 | 13 | 17.1 | 5.1 | 231.4 | 277.7 | 76.0% | 84.0% |
| SIG+FIT 50-85, 5_3 (8_12) | 6,591 | 4,294 | 0 | 0 | 1,005 | 1138.27 | 9.04 | 2,152 | 12 | 19.3 | 6.0 | 246.8 | 267.3 | 72.9% | 81.1% |
| SIG+FIT 55-75, 10_1 (3_21) | 10,820 | 1,686 | 0 | 0 | 845 | 1130.96 | 12.48 | 1,988 | 11 | 20.6 | 7.1 | 265.1 | 250.6 | 71.1% | 77.7% |
| SIG+FIT 55-75, 10_2 (3_11) | 6,646 | 1,907 | 0 | 0 | 695 | 1003.67 | 14.56 | 1,713 | 10 | 24.0 | 8.3 | 297.1 | 238.0 | 66.2% | 73.9% |
| SIG+FIT 55-75, 5_2 (5_11) | 6,180 | 2,913 | 0 | 0 | 797 | 1041.80 | 14.19 | 1,853 | 11 | 22.9 | 8.0 | 284.2 | 241.5 | 67.8% | 74.9% |
| SIG+FIT 55-75, 5_3 (5_7) | 4,399 | 3,059 | 0 | 0 | 728 | 968.88 | 16.89 | 1,713 | 10 | 25.3 | 9.1 | 297.4 | 230.0 | 64.4% | 71.3% |
| SIG+FIT 55-80, 10_1 (3_26) | 12,206 | 1,778 | 0 | 0 | 927 | 1158.79 | 10.16 | 2,096 | 12 | 19.1 | 6.2 | 263.7 | 255.7 | 73.2% | 80.5% |
| SIG+FIT 55-80, 10_2 (3_13) | 7,307 | 1,956 | 0 | 0 | 740 | 1022.34 | 12.56 | 1,775 | 11 | 23.0 | 7.6 | 297.8 | 242.3 | 67.7% | 76.2% |
| SIG+FIT 55-80, 5_2 (6_13) | 6,850 | 3,222 | 0 | 0 | 879 | 1068.12 | 11.54 | 1,959 | 12 | 21.3 | 7.0 | 282.0 | 246.2 | 70.1% | 78.0% |
| SIG+FIT 55-80, 5_3 (6_9) | 5,097 | 3,390 | 0 | 0 | 818 | 1003.52 | 13.22 | 1,834 | 11 | 23.2 | 7.8 | 295.5 | 237.3 | 67.4% | 75.3% |
| SIG+FIT 55-85, 10_1 (4_31) | 13,292 | 1,891 | 0 | 0 | 1,000 | 1174.91 | 8.36 | 2,184 | 13 | 18.4 | 5.6 | 265.9 | 258.3 | 74.1% | 82.2% |
| SIG+FIT 55-85, 10_2 (4_16) | 8,111 | 2,141 | 0 | 0 | 814 | 1043.96 | 9.78 | 1,868 | 12 | 22.0 | 6.7 | 298.6 | 245.4 | 69.1% | 78.7% |
| SIG+FIT 55-85, 5_2 (7_16) | 7,527 | 3,415 | 0 | 0 | 945 | 1083.62 | 9.54 | 2,038 | 13 | 20.6 | 6.4 | 284.7 | 249.0 | 71.0% | 79.7% |
| SIG+FIT 55-85, 5_3 (7_11) | 5,513 | 3,616 | 0 | 0 | 875 | 1018.11 | 11.25 | 1,904 | 12 | 22.6 | 7.3 | 298.0 | 239.5 | 68.3% | 77.0% |

**Table S42. Screening outcomes per 1,000 individuals aged 40 free from diagnosed CRC based on sigmoidoscopy plus guaiac-based fecal occult blood test (SIG+HSgFOBT) screening strategies.** Outcomes generated using period life tables. Outcomes table modeled from Zauber et al.^1^ COL, colonoscopy; SIG, flexible sigmoidoscopy; CTC, computed tomographic colonography; CRC, colorectal cancer; LY, life-years; LYG, life-years gained compared with no screening. ^a^Maximum possible number with this strategy. ^b^Including deaths from complications of screening. ^c^Compared to no screening.

| **Strategy** | **Outcomes per 1,000 persons free of diagnosed cancer at age 40** | | | | | | | | | | | |  | **Reductions^c^ (%)** | |
| --- | --- | --- | --- | --- | --- | --- | --- | --- | --- | --- | --- | --- | --- | --- | --- |
| **Modality** | **Screening tests** | | | | **Follow-up COLs** | **Surveillance COLs** | **COLs for symptoms** | **Total COLs** | **Complications** | **CRC cases** | **CRC deaths^b^** | **LY with CRC** | **LYG** |  |  |
| **Age to begin-age to end, screening interval (# of tests^a^)** | **Stool tests** | **SIGs** | **CTCs** | **COLs** |  |  |  |  |  |  |  |  |  | **Incidence** | **Mortality** |
| No screening | 0 | 0 | 0 | 0 | 0 | 0 | 71 | 71 | 2 | 71.1 | 31.7 | 560.3 | 0.0 | 0.0% | 0.0% |
| SIG+HSgFOBT 45-75, 10_1 (4_31) | 14,074 | 1,894 | 0 | 0 | 1,478 | 1,382 | 9 | 2,868 | 11 | 14.9 | 5.0 | 163.6 | 300.3 | 79.1% | 84.3% |
| SIG+HSgFOBT 45-75, 10_2 (4_16) | 9,216 | 2,312 | 0 | 0 | 1,160 | 1,242 | 11 | 2,413 | 11 | 18.2 | 6.0 | 201.3 | 286.9 | 74.5% | 80.9% |
| SIG+HSgFOBT 45-75, 5_2 (7_16) | 8,615 | 3,728 | 0 | 0 | 1,291 | 1,289 | 10 | 2,590 | 11 | 16.8 | 5.6 | 185.3 | 291.2 | 76.4% | 82.3% |
| SIG+HSgFOBT 45-75, 5_3 (7_11) | 6,416 | 4,087 | 0 | 0 | 1,162 | 1,209 | 12 | 2,383 | 11 | 18.9 | 6.5 | 202.8 | 280.1 | 73.5% | 79.4% |
| SIG+HSgFOBT 45-80, 10_1 (4_36) | 15,173 | 2,067 | 0 | 0 | 1,597 | 1,411 | 6 | 3,014 | 13 | 13.0 | 3.9 | 161.5 | 306.4 | 81.8% | 87.6% |
| SIG+HSgFOBT 45-80, 10_2 (4_18) | 9,747 | 2,440 | 0 | 0 | 1,228 | 1,264 | 9 | 2,500 | 11 | 16.7 | 5.2 | 198.7 | 291.5 | 76.5% | 83.5% |
| SIG+HSgFOBT 45-80, 5_2 (8_18) | 9,205 | 4,017 | 0 | 0 | 1,385 | 1,315 | 8 | 2,707 | 12 | 15.1 | 4.7 | 183.4 | 296.4 | 78.7% | 85.2% |
| SIG+HSgFOBT 45-80, 5_3 (8_12) | 6,973 | 4,359 | 0 | 0 | 1,250 | 1,234 | 9 | 2,494 | 12 | 17.3 | 5.6 | 200.5 | 285.7 | 75.7% | 82.4% |
| SIG+HSgFOBT 45-85, 10_1 (5_41) | 16,091 | 2,108 | 0 | 0 | 1,681 | 1,427 | 5 | 3,113 | 14 | 12.3 | 3.4 | 162.0 | 308.2 | 82.7% | 89.2% |
| SIG+HSgFOBT 45-85, 10_2 (5_21) | 10,477 | 2,547 | 0 | 0 | 1,311 | 1,282 | 6 | 2,599 | 13 | 15.9 | 4.5 | 201.7 | 294.8 | 77.7% | 85.9% |
| SIG+HSgFOBT 45-85, 5_2 (9_21) | 9,819 | 4,178 | 0 | 0 | 1,463 | 1,330 | 6 | 2,798 | 13 | 14.4 | 4.1 | 184.8 | 299.4 | 79.8% | 87.1% |
| SIG+HSgFOBT 45-85, 5_3 (9_14) | 7,404 | 4,564 | 0 | 0 | 1,320 | 1,252 | 7 | 2,579 | 13 | 16.5 | 5.0 | 201.5 | 288.4 | 76.9% | 84.2% |
| SIG+HSgFOBT 50-75, 10_1 (3_26) | 11,544 | 1,680 | 0 | 0 | 1,258 | 1,275 | 10 | 2,543 | 11 | 16.9 | 5.8 | 204.0 | 279.5 | 76.2% | 81.7% |
| SIG+HSgFOBT 50-75, 10_2 (3_13) | 7,399 | 1,940 | 0 | 0 | 969 | 1,132 | 13 | 2,115 | 10 | 20.9 | 7.3 | 238.0 | 263.9 | 70.6% | 76.9% |
| SIG+HSgFOBT 50-75, 5_2 (6_13) | 6,997 | 3,184 | 0 | 0 | 1,095 | 1,182 | 12 | 2,289 | 11 | 19.1 | 6.6 | 224.4 | 269.5 | 73.1% | 79.1% |
| SIG+HSgFOBT 50-75, 5_3 (6_9) | 5,385 | 3,447 | 0 | 0 | 1,000 | 1,110 | 14 | 2,123 | 10 | 21.1 | 7.4 | 239.9 | 259.5 | 70.4% | 76.6% |
| SIG+HSgFOBT 50-80, 10_1 (4_31) | 12,814 | 1,765 | 0 | 0 | 1,378 | 1,304 | 8 | 2,690 | 12 | 15.2 | 4.7 | 201.7 | 285.4 | 78.6% | 85.0% |
| SIG+HSgFOBT 50-80, 10_2 (4_16) | 8,410 | 2,132 | 0 | 0 | 1,089 | 1,168 | 9 | 2,266 | 12 | 18.5 | 5.8 | 236.8 | 272.6 | 74.0% | 81.5% |
| SIG+HSgFOBT 50-80, 5_2 (7_16) | 7,867 | 3,419 | 0 | 0 | 1,207 | 1,213 | 9 | 2,429 | 12 | 17.3 | 5.5 | 223.0 | 275.5 | 75.6% | 82.5% |
| SIG+HSgFOBT 50-80, 5_3 (7_11) | 5,890 | 3,745 | 0 | 0 | 1,088 | 1,135 | 11 | 2,234 | 11 | 19.4 | 6.4 | 238.8 | 265.8 | 72.8% | 79.7% |
| SIG+HSgFOBT 50-85, 10_1 (4_36) | 13,592 | 1,896 | 0 | 0 | 1,467 | 1,320 | 6 | 2,793 | 14 | 14.4 | 4.2 | 202.3 | 287.7 | 79.8% | 86.7% |
| SIG+HSgFOBT 50-85, 10_2 (4_18) | 8,789 | 2,227 | 0 | 0 | 1,140 | 1,179 | 8 | 2,327 | 13 | 18.0 | 5.4 | 237.5 | 273.9 | 74.8% | 82.8% |
| SIG+HSgFOBT 50-85, 5_2 (8_18) | 8,296 | 3,626 | 0 | 0 | 1,274 | 1,227 | 7 | 2,508 | 13 | 16.5 | 5.0 | 223.5 | 278.3 | 76.7% | 84.2% |
| SIG+HSgFOBT 50-85, 5_3 (8_12) | 6,291 | 3,932 | 0 | 0 | 1,152 | 1,149 | 9 | 2,311 | 12 | 18.7 | 5.9 | 240.2 | 267.5 | 73.7% | 81.3% |
| SIG+HSgFOBT 55-75, 10_1 (3_21) | 9,287 | 1,430 | 0 | 0 | 1,055 | 1,133 | 13 | 2,200 | 11 | 20.3 | 7.1 | 259.6 | 250.5 | 71.4% | 77.6% |
| SIG+HSgFOBT 55-75, 10_2 (3_11) | 6,144 | 1,708 | 0 | 0 | 847 | 1,014 | 15 | 1,875 | 11 | 23.5 | 8.3 | 288.6 | 237.9 | 66.9% | 73.9% |
| SIG+HSgFOBT 55-75, 5_2 (5_11) | 5,756 | 2,614 | 0 | 0 | 929 | 1,051 | 14 | 1,994 | 11 | 22.3 | 7.9 | 277.0 | 241.4 | 68.7% | 75.2% |
| SIG+HSgFOBT 55-75, 5_3 (5_7) | 4,229 | 2,823 | 0 | 0 | 828 | 980 | 17 | 1,825 | 10 | 24.8 | 9.0 | 290.4 | 230.0 | 65.1% | 71.5% |
| SIG+HSgFOBT 55-80, 10_1 (3_26) | 10,409 | 1,554 | 0 | 0 | 1,169 | 1,161 | 10 | 2,340 | 12 | 18.6 | 6.2 | 256.3 | 255.9 | 73.9% | 80.5% |
| SIG+HSgFOBT 55-80, 10_2 (3_13) | 6,714 | 1,786 | 0 | 0 | 911 | 1,035 | 13 | 1,959 | 11 | 22.3 | 7.5 | 288.4 | 242.5 | 68.6% | 76.3% |
| SIG+HSgFOBT 55-80, 5_2 (6_13) | 6,345 | 2,889 | 0 | 0 | 1,021 | 1,077 | 12 | 2,109 | 12 | 20.8 | 7.0 | 275.0 | 246.4 | 70.8% | 78.0% |
| SIG+HSgFOBT 55-80, 5_3 (6_9) | 4,885 | 3,124 | 0 | 0 | 931 | 1,014 | 13 | 1,958 | 11 | 22.7 | 7.8 | 289.6 | 236.9 | 68.1% | 75.4% |
| SIG+HSgFOBT 55-85, 10_1 (4_31) | 11,324 | 1,611 | 0 | 0 | 1,256 | 1,177 | 8 | 2,442 | 14 | 17.8 | 5.6 | 256.7 | 258.4 | 74.9% | 82.3% |
| SIG+HSgFOBT 55-85, 10_2 (4_16) | 7,449 | 1,917 | 0 | 0 | 997 | 1,054 | 10 | 2,061 | 13 | 21.4 | 6.7 | 290.0 | 246.1 | 70.0% | 78.9% |
| SIG+HSgFOBT 55-85, 5_2 (7_16) | 6,977 | 3,052 | 0 | 0 | 1,101 | 1,093 | 10 | 2,203 | 13 | 20.0 | 6.4 | 276.4 | 249.4 | 71.9% | 79.9% |
| SIG+HSgFOBT 55-85, 5_3 (7_11) | 5,249 | 3,332 | 0 | 0 | 996 | 1,028 | 11 | 2,035 | 12 | 22.0 | 7.2 | 290.9 | 239.9 | 69.0% | 77.2% |

**Table S43. Screening outcomes per 1,000 individuals aged 40 free from diagnosed CRC based on computed tomography colonography (CTC) screening strategies.** Outcomes generated using period life tables. Outcomes table modeled from Zauber et al.^1^ COL, colonoscopy; SIG, flexible sigmoidoscopy; CTC, computed tomographic colonography; CRC, colorectal cancer; LY, life-years; LYG, life-years gained compared with no screening. ^a^Maximum possible number with this strategy. ^b^Including deaths from complications of screening. ^c^Compared to no screening.

| **Strategy** | **Outcomes per 1,000 persons free of diagnosed cancer at age 40** | | | | | | | | | | | |  | **Reductions^c^ (%)** | |
| --- | --- | --- | --- | --- | --- | --- | --- | --- | --- | --- | --- | --- | --- | --- | --- |
| **Modality** | **Screening tests** | | | | **Follow-up COLs** | **Surveillance COLs** | **COLs for symptoms** | **Total COLs** | **Complications** | **CRC cases** | **CRC deaths^b^** | **LY with CRC** | **LYG** |  |  |
| **Age to begin-age to end, screening interval (# of tests^a^)** | **Stool tests** | **SIGs** | **CTCs** | **COLs** |  |  |  |  |  |  |  |  |  | **Incidence** | **Mortality** |
| No screening | 0 | 0 | 0 | 0 | 0 | 0 | 71 | 71 | 2 | 71.1 | 31.7 | 560.3 | 0.0 | 0.0% | 0.0% |
| CTC 45-75, 5 (7) | 0 | 0 | 5,031 | 0 | 805 | 1,175 | 11 | 1,991 | 10 | 17.0 | 6.1 | 174.4 | 285.0 | 76.1% | 80.8% |
| CTC 45-75, 10 (4) | 0 | 0 | 3,232 | 0 | 576 | 994 | 16 | 1,586 | 9 | 22.6 | 8.4 | 227.8 | 255.2 | 68.2% | 73.4% |
| CTC 45-80, 5 (8) | 0 | 0 | 5,405 | 0 | 869 | 1,198 | 9 | 2,076 | 11 | 15.3 | 5.2 | 172.9 | 289.5 | 78.4% | 83.6% |
| CTC 45-80, 10 (4) | 0 | 0 | 3,232 | 0 | 576 | 994 | 16 | 1,586 | 9 | 22.6 | 8.4 | 227.8 | 255.2 | 68.2% | 73.4% |
| CTC 45-85, 5 (9) | 0 | 0 | 5,662 | 0 | 914 | 1,209 | 7 | 2,130 | 12 | 14.7 | 4.7 | 174.5 | 291.7 | 79.3% | 85.1% |
| CTC 45-85, 10 (5) | 0 | 0 | 3,540 | 0 | 640 | 1,019 | 13 | 1,672 | 11 | 21.2 | 7.4 | 228.7 | 259.6 | 70.2% | 76.6% |
| CTC 50-75, 5 (6) | 0 | 0 | 4,204 | 0 | 705 | 1,099 | 13 | 1,817 | 10 | 19.0 | 7.0 | 210.4 | 264.9 | 73.3% | 78.0% |
| CTC 50-75, 10 (3) | 0 | 0 | 2,500 | 0 | 464 | 904 | 21 | 1,389 | 8 | 26.6 | 10.5 | 259.0 | 231.3 | 62.6% | 66.9% |
| CTC 50-80, 5 (7) | 0 | 0 | 4,580 | 0 | 769 | 1,122 | 10 | 1,902 | 11 | 17.2 | 6.0 | 207.5 | 270.7 | 75.8% | 81.1% |
| CTC 50-80, 10 (4) | 0 | 0 | 2,949 | 0 | 556 | 949 | 15 | 1,520 | 10 | 23.0 | 8.5 | 254.5 | 241.1 | 67.6% | 73.2% |
| CTC 50-85, 5 (8) | 0 | 0 | 4,837 | 0 | 813 | 1,133 | 8 | 1,955 | 12 | 16.7 | 5.6 | 209.2 | 271.6 | 76.6% | 82.5% |
| CTC 50-85, 10 (4) | 0 | 0 | 2,949 | 0 | 556 | 949 | 15 | 1,520 | 10 | 23.0 | 8.5 | 254.5 | 241.1 | 67.6% | 73.2% |
| CTC 55-75, 5 (5) | 0 | 0 | 3,412 | 0 | 608 | 992 | 15 | 1,615 | 10 | 22.2 | 8.3 | 260.1 | 237.1 | 68.8% | 73.8% |
| CTC 55-75, 10 (3) | 0 | 0 | 2,300 | 0 | 458 | 855 | 19 | 1,333 | 9 | 26.9 | 10.2 | 296.4 | 215.3 | 62.2% | 67.7% |
| CTC 55-80, 5 (6) | 0 | 0 | 3,788 | 0 | 672 | 1,016 | 12 | 1,700 | 11 | 20.4 | 7.3 | 257.6 | 242.1 | 71.4% | 76.9% |
| CTC 55-80, 10 (3) | 0 | 0 | 2,300 | 0 | 458 | 855 | 19 | 1,333 | 9 | 26.9 | 10.2 | 296.4 | 215.3 | 62.2% | 67.7% |
| CTC 55-85, 5 (7) | 0 | 0 | 4,045 | 0 | 717 | 1,028 | 11 | 1,756 | 12 | 19.7 | 6.8 | 258.0 | 244.0 | 72.2% | 78.4% |
| CTC 55-85, 10 (4) | 0 | 0 | 2,609 | 0 | 524 | 880 | 16 | 1,420 | 11 | 25.4 | 9.2 | 297.8 | 220.0 | 64.3% | 71.0% |

**Table S44. Efficiency outcomes for stool-based screening strategies per 1,000 individuals aged 40.** Outcomes generated using period life tables. Ages to start screening are 50 and 55 and ages to end screening are 75, 80, and 85. FIT, fecal immunochemical test; HSgFOBT, highly-sensitive guaiac-based fecal occult blood test; FIT-DNA, fecal immunochemical test with a DNA stool test; COL, colonoscopy; LYG, life-years gained compared with no screening; CRC, colorectal cancer; ΔCOL, incremental number of colonoscopies compared with the next-best non-dominated strategy; ΔLYG, incremental number of life-years gained compared with the next best non-dominated strategy. ^a^Maximum possible number with this strategy.

| **Model/strategy** | **Outcomes per 1,000 40-year-olds** | | | | | | | |
| --- | --- | --- | --- | --- | --- | --- | --- | --- |
| **Screening modality, age to begin-age to end, screening interval (# of tests^a^)** | **Stool tests** | **COLs** | **LYG** | **CRC deaths averted** | **ΔCOL** | **ΔLYG** | **Efficiency ratio (ΔCOL / ΔLYG)** | **Category** |
| FIT 55-75, 3 (7) | 5,305 | 954 | 170.6 | 16.3 |  |  |  | Efficient |
| FIT 55-80, 3 (9) | 6,263 | 1,057 | 184.4 | 18.4 |  |  |  | Weakly Dominated |
| FIT 55-85, 3 (11) | 6,851 | 1,110 | 189.4 | 19.5 |  |  |  | Weakly Dominated |
| FIT 50-75, 3 (9) | 6,865 | 1,135 | 201.4 | 18.3 | 181 | 31 | 6 | Efficient |
| HSgFOBT 55-75, 3 (7) | 5,055 | 1,136 | 172.9 | 16.7 |  |  |  | Strongly Dominated |
| FIT 50-80, 3 (11) | 7,677 | 1,213 | 212.0 | 20.0 | 78 | 11 | 7 | Efficient |
| FIT 55-75, 2 (11) | 7,593 | 1,218 | 203.5 | 19.7 |  |  |  | Strongly Dominated |
| FIT 50-85, 3 (12) | 8,105 | 1,250 | 215.3 | 20.7 | 37 | 3 | 11 | Near Efficient |
| HSgFOBT 55-80, 3 (9) | 5,941 | 1,259 | 187.2 | 18.9 |  |  |  | Strongly Dominated |
| FIT 55-80, 2 (13) | 8,439 | 1,292 | 212.0 | 21.1 |  |  |  | Strongly Dominated |
| mt-sDNA 55-75, 5 (5) | 3,578 | 1,313 | 190.3 | 18.8 |  |  |  | Strongly Dominated |
| HSgFOBT 55-85, 3 (11) | 6,442 | 1,323 | 191.2 | 19.8 |  |  |  | Strongly Dominated |
| FIT 55-85, 2 (16) | 9,352 | 1,366 | 217.2 | 22.1 |  |  |  | Weakly Dominated |
| HSgFOBT 50-75, 3 (9) | 6,502 | 1,366 | 205.2 | 18.9 |  |  |  | Strongly Dominated |
| mt-sDNA 55-80, 5 (6) | 3,979 | 1,391 | 196.1 | 20.0 |  |  |  | Strongly Dominated |
| FIT 50-75, 2 (13) | 9,265 | 1,402 | 230.2 | 21.0 | 189 | 18 | 10 | Near Efficient |
| mt-sDNA 55-85, 5 (7) | 4,252 | 1,443 | 199.2 | 20.6 |  |  |  | Strongly Dominated |
| HSgFOBT 55-75, 2 (11) | 6,969 | 1,444 | 206.2 | 20.2 |  |  |  | Strongly Dominated |
| HSgFOBT 50-80, 3 (11) | 7,196 | 1,456 | 214.2 | 20.4 |  |  |  | Strongly Dominated |
| HSgFOBT 50-85, 3 (12) | 7,661 | 1,510 | 217.3 | 21.1 |  |  |  | Strongly Dominated |
| FIT 50-80, 2 (16) | 10,520 | 1,510 | 241.7 | 22.8 | 297 | 30 | 10 | Efficient |
| mt-sDNA 50-75, 5 (6) | 4,381 | 1,524 | 217.6 | 20.5 |  |  |  | Strongly Dominated |
| HSgFOBT 55-80, 2 (13) | 7,723 | 1,538 | 213.5 | 21.4 |  |  |  | Strongly Dominated |
| FIT 50-85, 2 (18) | 11,130 | 1,557 | 244.9 | 23.5 | 46 | 3 | 15 | Efficient |
| mt-sDNA 55-75, 3 (7) | 4,656 | 1,575 | 217.8 | 21.2 |  |  |  | Strongly Dominated |
| mt-sDNA 50-80, 5 (7) | 4,781 | 1,599 | 224.4 | 21.6 |  |  |  | Strongly Dominated |
| HSgFOBT 55-85, 2 (16) | 8,542 | 1,630 | 219.4 | 22.4 |  |  |  | Strongly Dominated |
| mt-sDNA 50-85, 5 (8) | 5,052 | 1,650 | 225.8 | 22.1 |  |  |  | Strongly Dominated |
| FIT 55-75, 1 (21) | 12,373 | 1,658 | 235.3 | 22.9 |  |  |  | Strongly Dominated |
| HSgFOBT 50-75, 2 (13) | 8,462 | 1,676 | 233.7 | 21.5 |  |  |  | Strongly Dominated |
| mt-sDNA 55-80, 3 (9) | 5,411 | 1,716 | 227.7 | 22.9 |  |  |  | Strongly Dominated |
| FIT 55-80, 1 (26) | 14,050 | 1,776 | 243.0 | 24.3 |  |  |  | Strongly Dominated |
| mt-sDNA 55-85, 3 (11) | 5,813 | 1,787 | 231.0 | 23.6 |  |  |  | Strongly Dominated |
| HSgFOBT 50-80, 2 (16) | 9,583 | 1,812 | 244.8 | 23.2 |  |  |  | Strongly Dominated |
| FIT 55-85, 1 (31) | 15,246 | 1,854 | 245.9 | 24.9 |  |  |  | Weakly Dominated |
| mt-sDNA 50-75, 3 (9) | 5,958 | 1,866 | 250.2 | 23.2 |  |  |  | Weakly Dominated |
| HSgFOBT 50-85, 2 (18) | 10,127 | 1,872 | 247.1 | 23.8 |  |  |  | Strongly Dominated |
| FIT 50-75, 1 (26) | 15,563 | 1,937 | 265.3 | 24.5 | 381 | 20 | 19 | Near Efficient |
| HSgFOBT 55-75, 1 (21) | 10,504 | 1,942 | 237.4 | 23.2 |  |  |  | Strongly Dominated |
| mt-sDNA 50-80, 3 (11) | 6,517 | 1,969 | 258.7 | 24.5 |  |  |  | Strongly Dominated |
| mt-sDNA 50-85, 3 (12) | 6,957 | 2,041 | 261.6 | 25.1 |  |  |  | Strongly Dominated |
| FIT 50-80, 1 (31) | 17,225 | 2,052 | 273.4 | 25.8 | 495 | 29 | 17 | Efficient |
| HSgFOBT 55-80, 1 (26) | 11,872 | 2,086 | 244.5 | 24.5 |  |  |  | Strongly Dominated |
| FIT 50-85, 1 (36) | 18,423 | 2,128 | 276.2 | 26.4 | 76 | 3 | 27 | Efficient |
| HSgFOBT 55-85, 1 (31) | 12,862 | 2,186 | 247.3 | 25.1 |  |  |  | Strongly Dominated |
| HSgFOBT 50-75, 1 (26) | 13,084 | 2,285 | 268.2 | 24.8 |  |  |  | Strongly Dominated |
| mt-sDNA 55-75, 1 (21) | 8,752 | 2,339 | 253.1 | 25.0 |  |  |  | Strongly Dominated |
| HSgFOBT 50-80, 1 (31) | 14,447 | 2,429 | 274.3 | 25.9 |  |  |  | Strongly Dominated |
| mt-sDNA 55-80, 1 (26) | 9,825 | 2,495 | 258.5 | 25.9 |  |  |  | Strongly Dominated |
| HSgFOBT 50-85, 1 (36) | 15,430 | 2,528 | 277.5 | 26.6 | 401 | 1 | 295 | Near Efficient |
| mt-sDNA 55-85, 1 (31) | 10,604 | 2,607 | 260.9 | 26.4 |  |  |  | Strongly Dominated |
| mt-sDNA 50-75, 1 (26) | 10,891 | 2,733 | 284.1 | 26.4 | 605 | 8 | 77 | Near Efficient |
| mt-sDNA 50-80, 1 (31) | 11,979 | 2,892 | 288.4 | 27.3 | 764 | 12 | 63 | Near Efficient |
| mt-sDNA 50-85, 1 (36) | 12,759 | 3,002 | 290.6 | 27.8 | 875 | 14 | 60 | Efficient |

**Table S45. Efficiency outcomes for colonoscopy screening strategies per 1,000 individuals aged 40.** Outcomes generated using period life tables. Ages to start screening are 50 and 55 and ages to end screening are 75, 80, and 85. FIT, fecal immunochemical test; HSgFOBT, highly-sensitive guaiac-based fecal occult blood test; FIT-DNA, fecal immunochemical test with a DNA stool test; COL, colonoscopy; LYG, life-years gained compared with no screening; CRC, colorectal cancer; ΔCOL, incremental number of colonoscopies compared with the next-best non-dominated strategy; ΔLYG, incremental number of life-years gained compared with the next best non-dominated strategy. ^a^Maximum possible number with this strategy.

| **Model/strategy** | **Outcomes per 1,000 40-year-olds** | | | | | | | |
| --- | --- | --- | --- | --- | --- | --- | --- | --- |
| **Screening modality, age to begin-age to end, screening interval (# of tests^a^)** | **Stool tests** | **COLs** | **LYG** | **CRC deaths averted** | **ΔCOL** | **ΔLYG** | **Efficiency ratio (ΔCOL / ΔLYG)** | **Category** |
| COL 55-75, 15 (2) | 0 | 3,014 | 256.1 | 25.4 |  |  |  | Efficient |
| COL 55-80, 15 (2) | 0 | 3,014 | 256.1 | 25.4 |  |  |  | Strongly Dominated |
| COL 50-75, 15 (2) | 0 | 3,228 | 277.7 | 25.6 | 215 | 22 | 10 | Efficient |
| COL 55-85, 15 (3) | 0 | 3,331 | 258.8 | 26.3 |  |  |  | Strongly Dominated |
| COL 55-75, 10 (3) | 0 | 3,682 | 267.3 | 26.8 |  |  |  | Strongly Dominated |
| COL 55-80, 10 (3) | 0 | 3,682 | 267.3 | 26.8 |  |  |  | Strongly Dominated |
| COL 50-80, 15 (3) | 0 | 3,707 | 286.7 | 27.5 | 478 | 9 | 54 | Near Efficient |
| COL 50-85, 15 (3) | 0 | 3,707 | 286.7 | 27.5 |  |  |  | Strongly Dominated |
| COL 55-85, 10 (4) | 0 | 3,965 | 268.6 | 27.2 |  |  |  | Strongly Dominated |
| COL 50-75, 10 (3) | 0 | 4,023 | 293.1 | 27.5 | 795 | 15 | 52 | Efficient |
| COL 50-80, 10 (4) | 0 | 4,451 | 297.8 | 28.5 | 428 | 5 | 91 | Efficient |
| COL 50-85, 10 (4) | 0 | 4,451 | 297.8 | 28.5 |  |  |  | Strongly Dominated |
| COL 55-75, 5 (5) | 0 | 5,004 | 273.9 | 27.4 |  |  |  | Strongly Dominated |
| COL 55-80, 5 (6) | 0 | 5,378 | 275.3 | 27.8 |  |  |  | Strongly Dominated |
| COL 55-85, 5 (7) | 0 | 5,626 | 275.6 | 27.9 |  |  |  | Strongly Dominated |
| COL 50-75, 5 (6) | 0 | 6,015 | 304.7 | 28.8 | 1564 | 7 | 227 | Near Efficient |
| COL 50-80, 5 (7) | 0 | 6,389 | 306.6 | 29.2 | 1939 | 9 | 221 | Efficient |
| COL 50-85, 5 (8) | 0 | 6,637 | 306.8 | 29.3 | 248 | 0 | 1492 | Efficient |
